# The genomic imprint of chromosomal inversions and demographic history in island populations of deer mice

**DOI:** 10.1101/2025.04.18.649517

**Authors:** Emma K. Howell, Felix Baier, Hopi E. Hoekstra, Bret A. Payseur

## Abstract

Populations that colonize islands experience novel selective pressures, fluctuations in size, and changes to their connectivity. Owing to their unique geographic setting, islands can function as natural laboratories in which to examine the interactions between demographic history and natural selection replicated across isolated populations. We used whole genome sequences of wild-caught deer mice (*Peromyscus maniculatus*) from two islands (Saturna and Pender) and one mainland location (Maple Ridge) in the Gulf Islands region of coastal British Columbia to investigate two primary determinants of genome-wide diversity: chromosomal inversions and non-equilibrium demographic history. We found that segregating inversions produce characteristic, large-scale distortions in allele frequencies and linkage disequilibrium that make it possible to identify and characterize them from short-read sequence data. Patterns of variation within and between karyotypes indicate that six inversion polymorphisms have been maintained by a shared history of balancing selection in both island and mainland populations. Whereas the estimated timing of contemporary population splits is consistent with the isolation of island populations from each other following the Last Glacial Maximum, ancestral island and mainland lineages are inferred to have diverged much earlier. These aspects of demographic history suggest that shared inversions existed long ago in a common ancestor or spread via limited gene flow between ancestral island and mainland lineages. Our results raise the possibility that inversions segregating among Gulf Islands populations are on similar evolutionary trajectories, providing a contrast to previous findings in mainland *P. maniculatus* and contributing to the emerging portrait of inversion evolution in this species.

## INTRODUCTION

Island colonization alters the evolutionary trajectory of a population in multiple ways. Ecological differences between island and mainland environments in variables such as species composition, habitat diversity, resource availability, and climate impart novel selective challenges on colonizing populations (Losos and Ricklefs 2009). Coincident with these ecological shifts, founder effect bottlenecks amplify the effects of genetic drift and reductions in gene flow driven by island isolation increase differentiation between populations (Losos and Ricklefs 2009). Together, these unique selective and demographic changes that accompany evolution on islands shape the fates of standing variants and new mutations alike.

One source of genetic variation that may be particularly sensitive to these features of island evolution is structural changes to the genome. Compared to single nucleotide mutations, structural variants can be orders of magnitude larger in their genomic scope and change the collinearity of chromosome copies. Chromosomal inversions, which occur when a DNA segment is reinserted in the reverse orientation, can disrupt genes that stretch across breakpoints and alter the proximity of genetic elements within and outside of breakpoints, achieving substantial shifts in genome function and architecture in a single mutational step (reviewed in Berdan et al. 2023). Since Alfred Sturtevant’s pioneering discovery of inversions in *Drosophila* (Sturtevant 1921), this class of structural variation has been the focus of intensive evolutionary study aimed at understanding the origin, spread, and maintenance of inversions in populations (Hoffmann and Rieseberg 2008).

Central to the evolutionary significance of inversions is their effect on recombination. Crossing over that occurs within inversion breakpoints can produce unbalanced gametes in individuals that carry both the inverted and standard orientation (McClintock 1931). This underdominant behavior in heterokaryotypes, in addition to potentially deleterious positional and breakpoint effects, suggests that inversions may be more likely to spread in populations experiencing increased genetic drift (Lande 1979; Hedrick 1981; Bush et al. 1977), such as those inhabiting islands. Indeed, accelerated rates of inversion-driven karyotypic evolution in gibbons (Carbone et al. 2014) and annual sunflowers (Burke et al. 2004) have been attributed to dramatic fluctuations in population size. At the same time, inversions can promote adaptive evolution through their suppression of recombination. By preserving associations between alleles with epistatic fitness effects, inversions can maintain co-adapted variation (Dobzhansky 1948; Dobzhansky 1949; Dobzhansky 1950). Even in the absence of such epistatic interactions, inversions that capture locally adaptive haplotypes can spread when there is gene flow between populations experiencing divergent selective regimes (Kirkpatrick and Barton 2006). Empirical evidence from diverse taxa suggests that inversions are an important source of adaptive divergence between populations (reviewed in Wellenreuther and Bernatchez 2018).

*Peromyscus* provides a powerful system for examining chromosomal evolution on islands. As one of the most abundant, widespread rodents in North America (Osgood 1909; Bedford and Hoekstra 2015), the range of *Peromyscus* extends to a wide array of coastal islands that line the continent (Redfield 1976; Gill 1980; Adler and Tamarin 1983; Adler et al. 1986; Lucid and Cook 2004; Álvarez-Castañeda et al. 2010; Nolfo-Clements et al. 2017). Although all species share a 2n = 48 karyotype, there is substantial interspecific and intraspecific variation among chromosomes driven by pericentric inversions and heterochromatic additions (Ohno et al 1966; Robbins and Baker 1981; Stangl and Baker 1984; Greenbaum et al. 1994). In the Pacific Northwest, this karyotypic variation has helped to clarify taxonomic relationships between the deer mouse, *Peromyscus maniculatus*, and Keen’s mouse, *P. keeni*, where cytogenetic analyses have revealed differences both within and between species in the fundamental number of chromosome arms (Gunn and Greenbaum 1986; Allard and Greenbaum 1988; Gunn 1987; Hogan et al. 1993). Our understanding of such variability is being refined and expanded in the genomics era. Using a combination of long-read sequencing and population genomic data, Harringmeyer and Hoekstra (2022) provide a detailed characterization of 21 large-scale inversions in *P. maniculatus*, including evidence for substantial geographic variability among mainland populations in both the presence and frequencies of inversion polymorphisms. This genomic profile of inversions provides a useful framework for investigating how this important source of variation is partitioned among island and mainland populations.

Deer mice inhabiting the Gulf Islands of British Columbia offer compelling subjects for studying the interaction between demographic history and inversion evolution in island populations. Pleistocene-era glacial activity played an important role in shaping the contemporary distributions of species occupying higher latitudes (Hewitt 2000; Lessa et al. 2003). Glacial cover of the Gulf Islands during this period (Clague 1980; Clague and James 2002; Hutchinson et al. 2004; Fedje et al. 2009; James et al. 2009) suggests that island colonization could have been accompanied by population size fluctuations as mice dispersed from unglaciated regions or glacial refugia. For low vagility species like *P. maniculatus*, subsequent isolation of the islands from each other and from nearby mainland areas could impose formidable barriers to gene flow (Redfield 1976) that may enable inversions to evolve independently among populations.

Using whole genome sequences from two island populations and one mainland population, we show that segregating inversions have dramatic effects on patterns of variation that span a large portion of the genome. We leverage these genomic signatures to detect polymorphic inversions in short-read population genomic data and to draw inferences about their evolutionary history. By reconstructing the joint demographic history of island and mainland populations, we provide context for our evolutionary interpretations. Together, our genomic characterization of the Gulf Islands populations enriches and expands our current understanding of inversion evolution and population history in *P. maniculatus*.

## RESULTS

### Population sampling and structure

The Gulf Islands are situated at the southern end of the Strait of Georgia between Vancouver Island (to the west) and mainland British Columbia (to the east). Our sampling of *Peromyscus maniculatus* spans two of the southern Gulf Islands, Saturna (31 km^2^) and Pender (34 km^2^), and one nearby mainland site at Maple Ridge (Figure 1). Subsequent genomic analyses are based on a subset of unrelated individuals identified among mice sampled from each location (n=20 for Saturna Island, n=10 for Pender Island, and n=17 for mainland Maple Ridge; see Materials and Methods).

**Figure 1.**
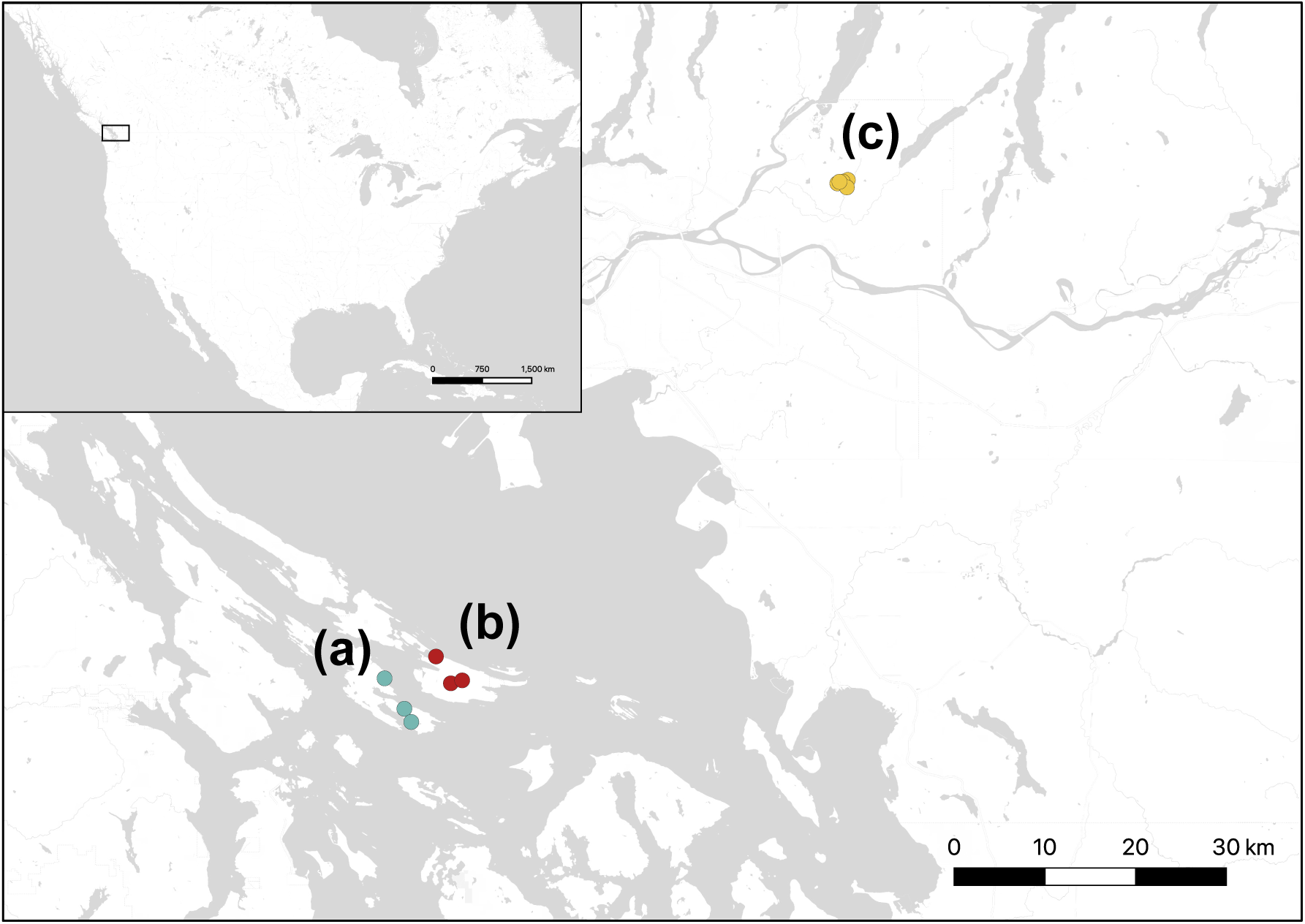
Map of the Gulf Islands in southwestern British Columbia. Land areas are depicted in white and water areas are depicted in gray. Points indicate sampled areas on Pender Island (a; blue), Saturna Island (b; red), and mainland Maple Ridge (c; yellow). Inset depicts location of the Gulf Islands region in the Pacific Northwest.

To characterize genetic relationships within and among island and mainland sites, we conducted principal component analysis (PCA) and global ancestry inference using genome-wide, unlinked single nucleotide polymorphisms (SNPs). PCA revealed three distinct clusters along the first and second principal component axes that separated mice according to sampling location (Figure 2A). Within this combined sample, cross-validation analyses performed with ADMIXTURE (Alexander et al. 2009) recovered three ancestral components (k=3) that almost entirely delineate individuals by sampling location (Figure 2F). In contrast, cross-validation analyses performed separately for each site supported a single ancestral component (k=1) (Figure 2B-D). Together, these findings provide evidence for structure between, but not within, each of the sampled locations.

**Figure 2.**
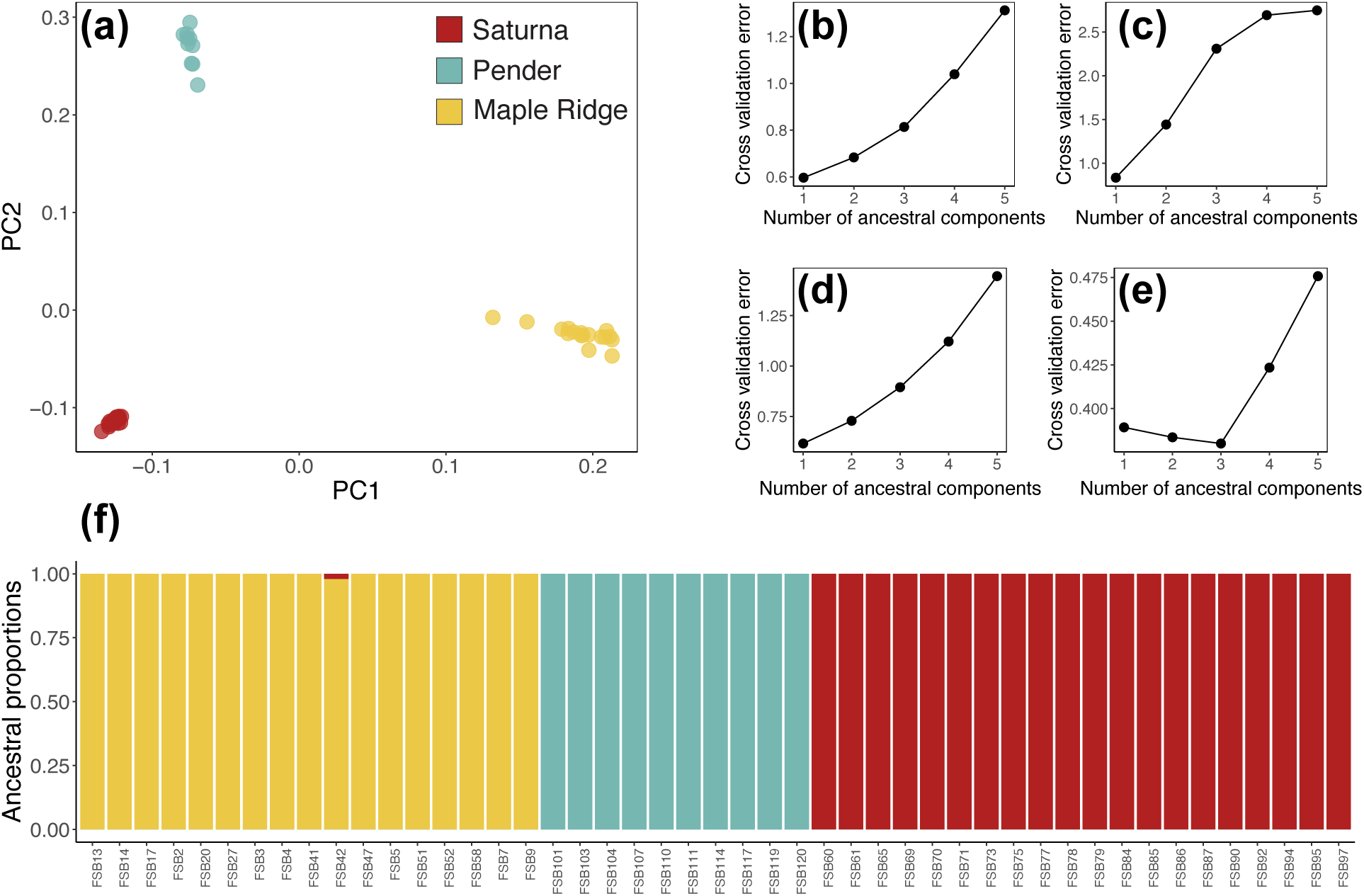
Evidence for structure among island and mainland sampling locations. Panel a depicts the results of PCA conducted using genome-wide unlinked SNPs. Each point represents an individual’s position along the first (x-axis) and second (y-axis) principal component. Individuals are colored according to sampling location (yellow for mainland Maple Ridge, blue for Pender Island, and red for Saturna Island). Percentage of variance explained by PC1=7.73% and by PC2=4.65%. Panels b-e present the results of five-fold cross validation analyses performed with ADMIXTURE (Alexander et al. 2009) to identify the best-fit number of ancestral components within the Saturna (b), Pender (c), Maple Ridge (d), and combined (e) samples. Panel f depicts global ancestry proportions estimated for the combined sample, assuming k=3 ancestral components. Each individual is represented by a colored bar where the height of each colored segment within the bar gives the proportion of ancestry an individual derives from each component. Ancestral component colors reflect sampling location (as in Panel a).

### Inversions drive large-scale distortions in allele frequencies and linkage disequilibrium

Leveraging existing knowledge about chromosomal rearrangements in *P. maniculatus*, we investigated how segregating inversions affect patterns of variation. Due to the suppression of recombination in heterokaryotypes, variation accumulated after the emergence of the inversion may remain private to either class, driving divergence between karyotypes in a manner analogous to local population subdivision. In Gulf Islands *P. maniculatus*, we found that this effect is sufficient to distort the site frequency spectrum (SFS) at the genome-wide level. Within each population, the folded SFS constructed from genome-wide variant calls exhibits subtle elevations in SNP counts at intermediate minor allele frequencies (Figure 3A-C; Supplementary Figure S1). Partitioning the SFS by chromosome, we found that these genome-wide irregularities are driven by extreme distortions in the chromosome-wide SFS for five autosomes: 7, 14, 15, 21, and 22 (Figure 3A-C; Supplementary Figure S1). In each case, the chromosome-wide SFS exhibits an unusually high density of SNPs at a specific minor allele frequency (MAF) bin, consistent with an excess of fixed differences between inverted and standard karyotypes.

**Figure 3.**
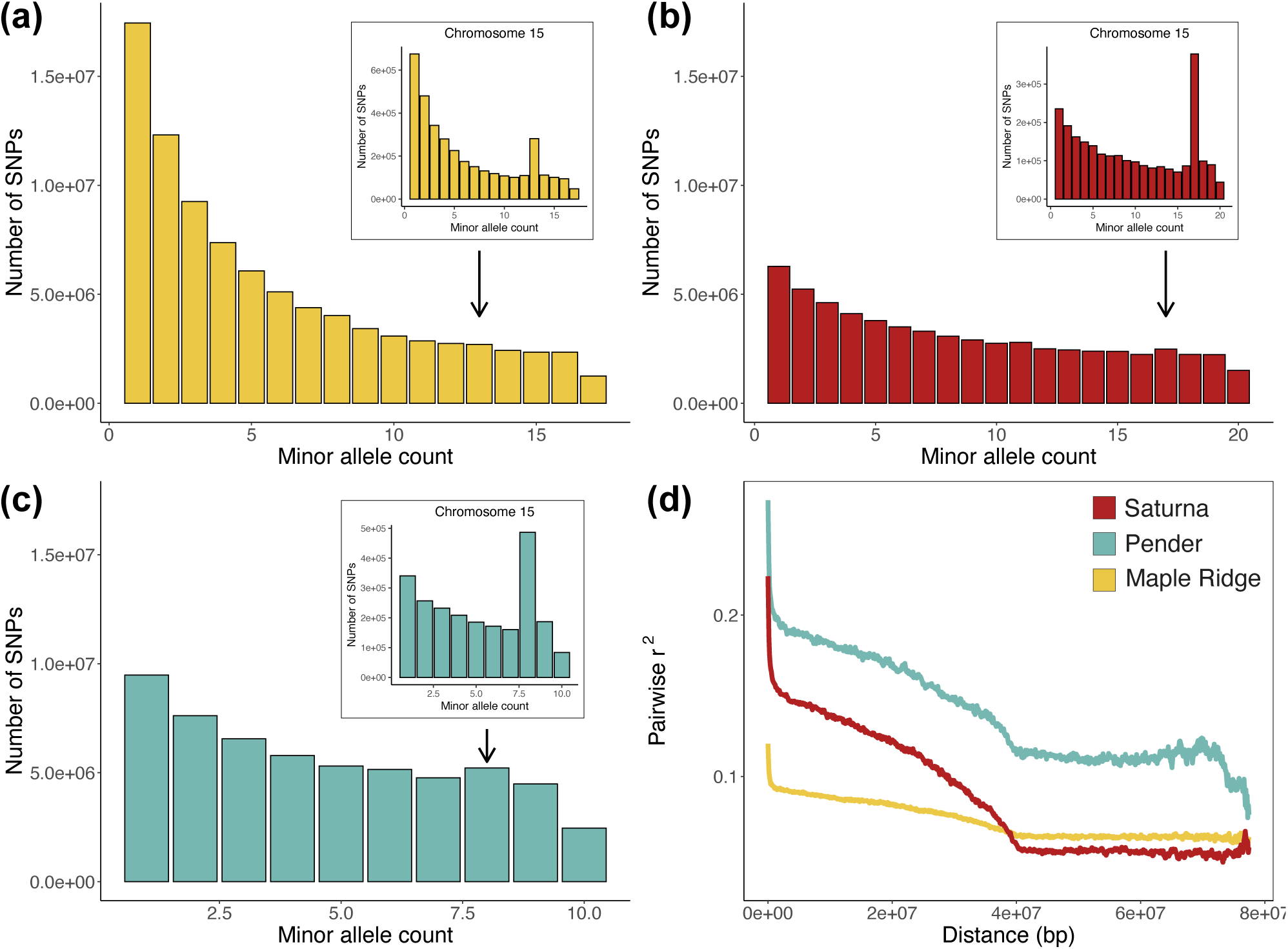
Inversion polymorphisms drive irregularities in the SFS and cause distortions in LD decay. Panels a-c provide an example of how the inversion polymorphism on chromosome 15 (inv15.0) affects both local and genome-wide allele frequencies in the Maple Ridge (a), Saturna Island (b), and Pender Island (c) samples. Main histograms plot the folded SFS constructed from genome-wide SNPs. Arrows indicate minor allele frequency bins in the genome-wide SFS that exhibit elevated SNP counts. Insets illustrate how these minor distortions in the genome-wide SFS correspond to major distortions in the local SFS for chromosome 15 SNPs. Panel d plots average pairwise r^2^ (y-axis) between chromosome 15 SNPs separated by increasing distance (x-axis). Each line represents LD decay within a different population sample (yellow for Maple Ridge, red for Saturna, and blue for Pender).

These same chromosomes also exhibit unusual patterns of linkage disequilibrium (LD). We found that pairwise r^2^ across these five chromosomes is elevated over intervals as large as 40 Mbp (Figure 3D; Supplementary Figure S1). Such distortions in LD decay could result from the mis-orientation of sampled sequences during their alignment to the reference assembly: when short reads derived from within an inversion are mapped to an inversion-free reference, the re-assignment of genomic coordinates causes nearby SNPs to appear farther apart (and vice versa). In addition to this artificial elevation in LD, the observed patterns of LD decay may also reflect true reductions in recombination across regions harboring polymorphic inversions.

SNPs that contribute to the aberrant MAF bins in each chromosome-wide SFS localize to discrete genomic intervals that span the breakpoints identified by Harringmeyer and Hoekstra (2022) for inversions inv7.2, inv14.0, inv15.0, inv21.0, and inv22.0 (Figure 4A-E; Supplementary Figures S2-S3). These MAF-partitioned SNPs exhibit extreme genotype continuity across the entirety of each interval, such that each mouse almost exclusively carries heterozygous or homozygous genotypes at this subset of variants (Figure 4A-E; Supplementary Figures S2-S3). Together, these findings illustrate how the unique evolutionary dynamics induced by non-collinearity can cause patterns of variation spanning polymorphic inversions to depart substantially from genome-wide expectations.

**Figure 4.**
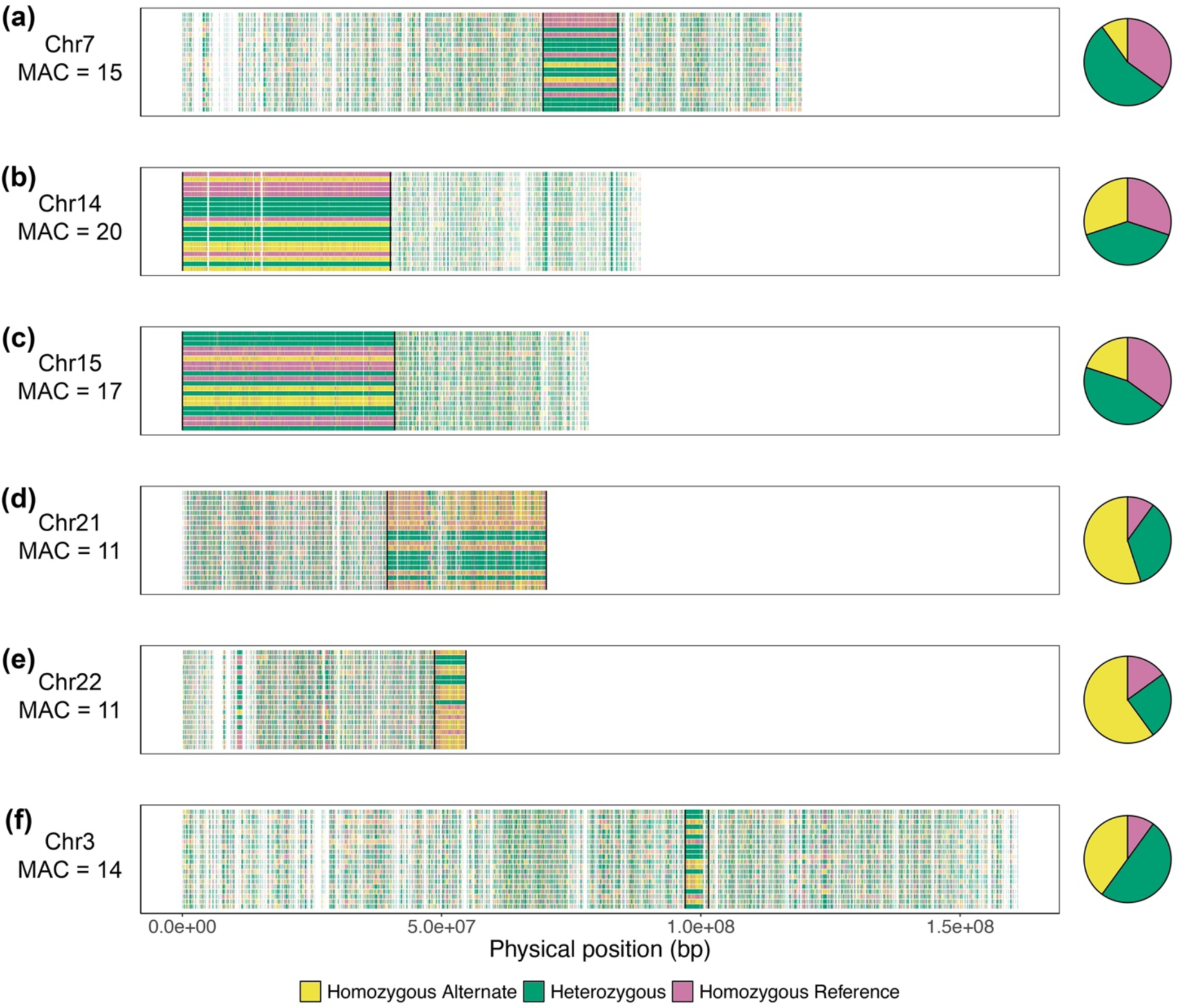
Partitioning SNP genotypes by minor allele frequency enables the detection and genotyping of polymorphic inversions. Plots illustrate the positive results for the inversion screen conducted in the Saturna Island sample. Each horizontal panel represents a different autosome: chr7 (a), chr14 (b), chr15 (c), chr21 (d), chr22 (e), and chr3 (f). Labels beneath each autosome name indicate the corresponding minor allele count of the inversion in the sample. X-axes measure the physical position along each autosome. Within a panel, each row represents a sampled individual, and each vertical tick mark indicates the location of a SNP with the specified minor allele count. SNPs are colored according to their diploid genotype in each individual (yellow for homozygous alternate, green for heterozygous, and pink for homozygous reference). Vertical black lines denote the breakpoints defined by Harringmeyer and Hoekstra (2022) for inv7.2 (a), inv14.0 (b), inv15.0 (c), inv21.0 (d), inv22.0 (e), and inv3.0 (f). Pie charts adjacent to each panel denote the genotype proportions of the arrangement in each sample. See Supplementary Figures S2 and S3 for corresponding plots for the Pender Island and mainland Maple Ridge samples.

We leveraged this diagnostic pattern to screen our short-read sequence data for additional segregating inversions by partitioning autosomal genotypes in each population by all possible minor allele frequencies. This approach revealed that a sixth locus on chromosome 3, inv3.0 (Harringmeyer and Hoekstra 2022), is also polymorphic (Figure 4F; Supplementary Figures S2-S3). The shorter span of this inversion is likely insufficient to drive the chromosome-wide distortions in the SFS and LD decay that were observed for the loci on chromosomes 7, 14, 15, 21, and 22. In total, our inversion screen indicates that the same six inversion polymorphisms are segregating in all three Gulf Islands populations.

Local PCA conducted using SNPs spanning the breakpoints of inv3.0, inv7.2, inv14.0, inv15.0, inv21.0, and inv22.0 yielded three distinct clusters along the first principal component axis (Figure 5). Whereas genome-wide SNPs group individuals by sampling location (Figure 5A), the clustering pattern observed at these loci instead reflects the three possible inversion genotypes (homozygous inversion, heterozygous, or homozygous standard; Figure 5B-G). We find little evidence that additional polymorphisms beyond the six described above are segregating in the Gulf Islands populations. With the possible exception of inv9.0 in the mainland Maple Ridge population, PCA conducted using SNPs within the 21 intervals defined by Harringmeyer and Hoekstra (2022) failed to produce clustering patterns consistent with distinct inversion genotypes (Supplementary Figure S4).

**Figure 5.**
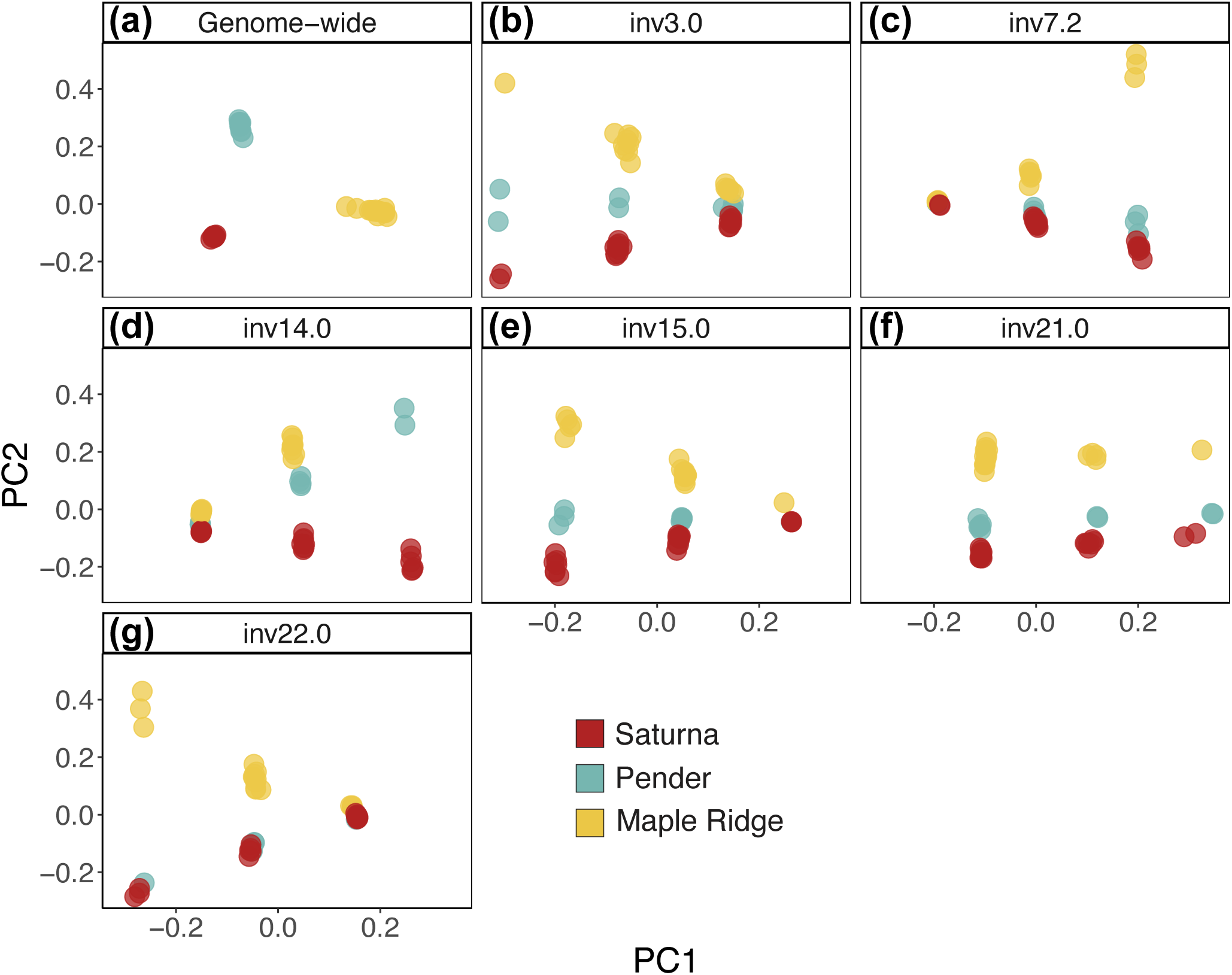
Individuals from distinct populations cluster according to inversion genotype at each polymorphic locus. Panels compare the results of PCA conducted on genome-wide SNPs falling outside of known inversions (a) to PCA performed on SNPs within each polymorphic inversion locus: inv3.0 (b), inv7.2 (c), inv14.0 (d), inv15.0 (e), inv21.0 (f), and inv22.0 (g). Each point denotes an individual’s position along the first (x-axis) and second (y-axis) principal component. Points are colored according to population sample (yellow for Maple Ridge, blue for Pender Island, and red for Saturna Island).

### Polarizing inversion polymorphisms

In order to draw inferences about the evolutionary dynamics of segregating inversions, we reconstructed the ancestral orientation at each locus by conducting pairwise whole genome alignments between the *P. maniculatus* reference assembly and genomes from four *Peromyscus* species: *P. polionotus*, *P. leucopus*, *P. eremicus*, and *P. californicus* (see Materials and Methods). Comparisons between *P. maniculatus* and these additional species revealed an abundance of large-scale inversions, duplications, and translocations (Supplementary Figures S5-S8). In some cases, these inter-species rearrangements span over 50 Mbp (Supplementary Figures S5-S8). By assessing collinearity across inversion intervals, we determined that the *P. maniculatus* reference genome carries the derived (i.e., inverted) orientation at inv14.0 and the ancestral (i.e., standard) orientation at inv3.0, inv7.2, inv15.0, inv21.0, and inv22.0 (Supplementary Figures S5-S8). For each locus, we assigned individuals to one of three genotypic classes with respect to the *P. maniculatus* reference genome (homozygous ALT, heterozygous, and homozygous REF) based on their proportion of ALT alleles at (Supplementary Figure S9) or approaching (Supplementary Figures S10-S12) the MAF of the inversion (see Materials and Methods). Based on these polarizations, Table 1 summarizes the frequencies of the inversions at each polymorphic locus in the Gulf Islands populations.

**Table 1.** Polymorphic inversions in the Gulf Islands. Table 1. **All three Gulf Islands populations are polymorphic at the same inversion loci.** Column three summarizes observed frequencies of the inverted karyotype at each polymorphic inversion locus. Individual genotypes at each inversion locus were obtained by partitioning SNP genotypes by minor allele frequency (see Figure 4 and Supplementary Figures S2-S3). The orientation of each arrangement was then determined based on alignments between the *P. maniculatus* reference assembly and those of *P. polionotus*, *P. leucopus*, *P. eremicus*, and *P. californicus*. Columns four through six tabulate mean nucleotide diversity within standard karyotype (𝜋̅_std_), mean nucleotide diversity within inverted karyotype (𝜋̅_inv_) and mean divergence between karyotypes (𝑑̅_XY_^karyotype^) for each locus. Means are computed across 5 kbp windows. Columns eight and nine summarize the physical length and average continent-wide frequency of each inversion, as defined by Harringmeyer and Hoekstra (2022).

| **Table 1. Polymorphic inversions in the Gulf Islands** |  |  |  |  |  |  |  |
| --- | --- | --- | --- | --- | --- | --- | --- |
| Locus | Population | Inversion frequency | $\bar{\pi}_{\text{std}}$ | $\bar{\pi}_{\text{inv}}$ | $\bar{d}_{\text{XY}}^{\text{karyotype}}$ | Inversion length | Continent-wide frequency |
| inv3.0 | Saturna | 0.65 | 0.0049 | 0.0037 | 0.0150 | 4.5 Mbp | 0.183 |
|  | Pender | 0.7 | 0.0078 | 0.0040 | 0.0151 |  |  |
|  | Maple Ridge | 0.676 | 0.0108 | 0.0053 | 0.0154 |  |  |
| inv7.2 | Saturna | 0.375 | 0.0084 | 0.0010 | 0.0176 | 14.4 Mbp | 0.110 |
|  | Pender | 0.5 | 0.0086 | 0.0006 | 0.0178 |  |  |
|  | Maple Ridge | 0.647 | 0.0102 | 0.0010 | 0.0173 |  |  |
| inv14.0 | Saturna | 0.5 | 0.0019 | 0.0061 | 0.0168 | 40.1 Mbp | 0.862 |
|  | Pender | 0.4 | 0.0016 | 0.0066 | 0.0169 |  |  |
|  | Maple Ridge | 0.235 | – | – | – |  |  |
| inv15.0 | Saturna | 0.425 | 0.0092 | 0.0005 | 0.0192 | 40.9 Mbp | 0.161 |
|  | Pender | 0.4 | 0.0085 | 0.0005 | 0.0192 |  |  |
|  | Maple Ridge | 0.382 | 0.0119 | 0.0015 | 0.0186 |  |  |
| inv21.0 | Saturna | 0.725 | 0.0041 | 0.0069 | 0.0141 | 30.7 Mbp | 0.275 |
|  | Pender | 0.7 | 0.0019 | 0.0065 | 0.0147 |  |  |
|  | Maple Ridge | 0.824 | 0.0028 | 0.0085 | 0.0144 |  |  |
| inv22.0 | Saturna | 0.725 | 0.0073 | 0.0027 | 0.0171 | 6.1 Mbp | 0.571 |
|  | Pender | 0.6 | 0.0068 | 0.0016 | 0.0171 |  |  |
|  | Maple Ridge | 0.529 | 0.0109 | 0.0025 | 0.0164 |  |  |

Surprisingly, we also uncovered evidence that certain segregating inversions are shared across *Peromyscus* species. Collinearity between the *P. maniculatus* and *P. polionotus* genomes along the majority of the inv14.0 interval suggests that this rearrangement is shared between these sister species (Supplementary Figure S5). In addition, although the *P. maniculatus* assembly carries the standard orientation at inv22.0, evidence for this inversion within *P. leucopus* suggests that this rearrangement is shared across greater phylogenetic distances (Supplementary Figure S6).

### High divergence between karyotypes and variable levels of diversity within inversions

To gain insight into how the evolutionary histories of segregating inversions depart from colinear regions of the genome, we compared levels of nucleotide diversity measured at putatively neutral, non-inverted loci (𝜋_genome_), to three summaries of diversity spanning inversion loci: divergence between standard and inverted karyotypes (*d_XY_*^karyotype^), diversity within inverted karyotypes (𝜋_inv_), and diversity within the standard karyotypes (𝜋_std_). Table 1 summarizes the means of these distributions calculated in 5 kbp windows across each segregating locus in each population.

Each locus exhibits high levels of divergence between karyotypes (*d_XY_*^karyotype^). At all loci, mean *d_XY_*^karyotype^ measured in each population far exceeds corresponding levels of diversity measured within either karyotypic class (mean 𝜋_inv_ and mean 𝜋_std_) (Table 1). Divergence between karyotypes is also high relative to background levels of diversity. Across all populations and inversion loci, mean *d_XY_*^karyotype^ exceeds mean nucleotide diversity measured genome-wide at putatively neutral, non-inverted loci (𝜋_genome_) (Figure 6). This departure is greatest for inv15.0, which yields a mean *d_XY_*^karyotype^ that falls within the right tail of the 𝜋_genome_ distribution, and lowest for inv21.0 (Figure 6).

**Figure 6.**
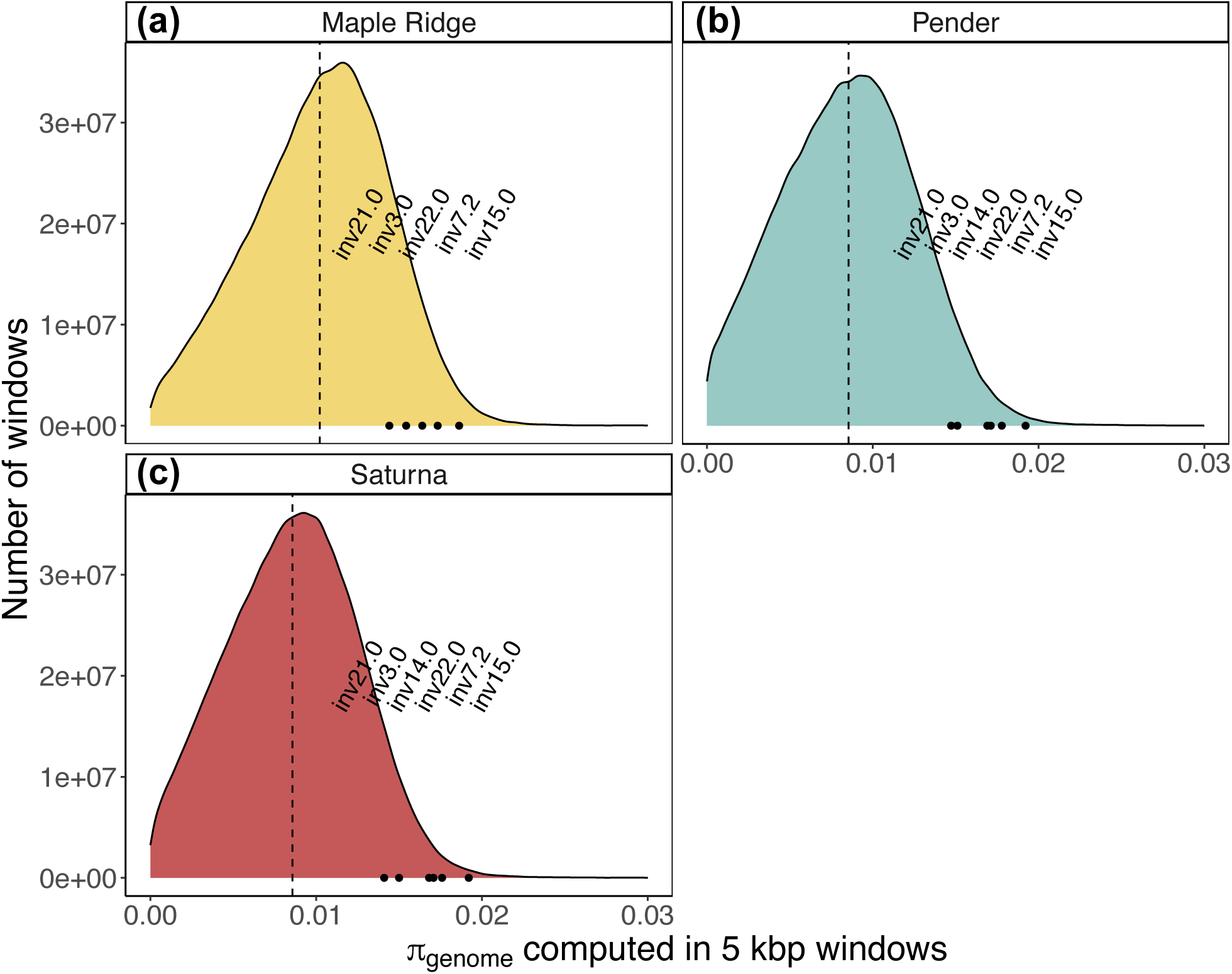
Divergence between karyotypes at polymorphic inversions exceeds mean genome-wide diversity. Density plots illustrate the distribution of nucleotide diversity computed in genome-wide non-overlapping 5 kbp windows outside of known inversion loci (𝜋_genome_) for the Maple Ridge (a), Pender Island (b), and Saturna Island (c) samples. Dashed black lines denote the mean 𝜋_genome_ observed in each population sample. Red points demonstrate where mean divergence between karyotypes (*d_XY_*^karyotype^) falls along the distribution of genome-wide nucleotide diversity for each polymorphic inversion.

Relative levels of nucleotide diversity within inverted (𝜋_inv_) and standard (𝜋_std_) karyotypes exhibit interesting differences across loci. For inv3.0, inv7.2, inv15.0, and inv22.0, inverted karyotypes exhibit reduced nucleotide diversity compared to their corresponding standard karyotypes (i.e., mean 𝜋_inv_ < mean 𝜋_std_) (Table 1). These reductions are most extreme for inversions at loci inv7.2 and inv15.0, where mean 𝜋_inv_ is less than 14% of mean 𝜋_std_. Both of these inversions feature broad-scale reductions in nucleotide diversity that span nearly the entirety of each locus (Figure 7). In contrast, we observed the opposite trend for inv14.0 and inv21.0, where the inverted karyotype exhibits higher nucleotide diversity (i.e., mean 𝜋_inv_ > mean 𝜋_std_) (Table 1). Despite this heterogeneity among inversion loci, all three Gulf Islands populations show remarkable similarity in relative levels of *d_XY_*^karyotype^, 𝜋_std_, and 𝜋_inv_ at any given locus.

**Figure 7.**
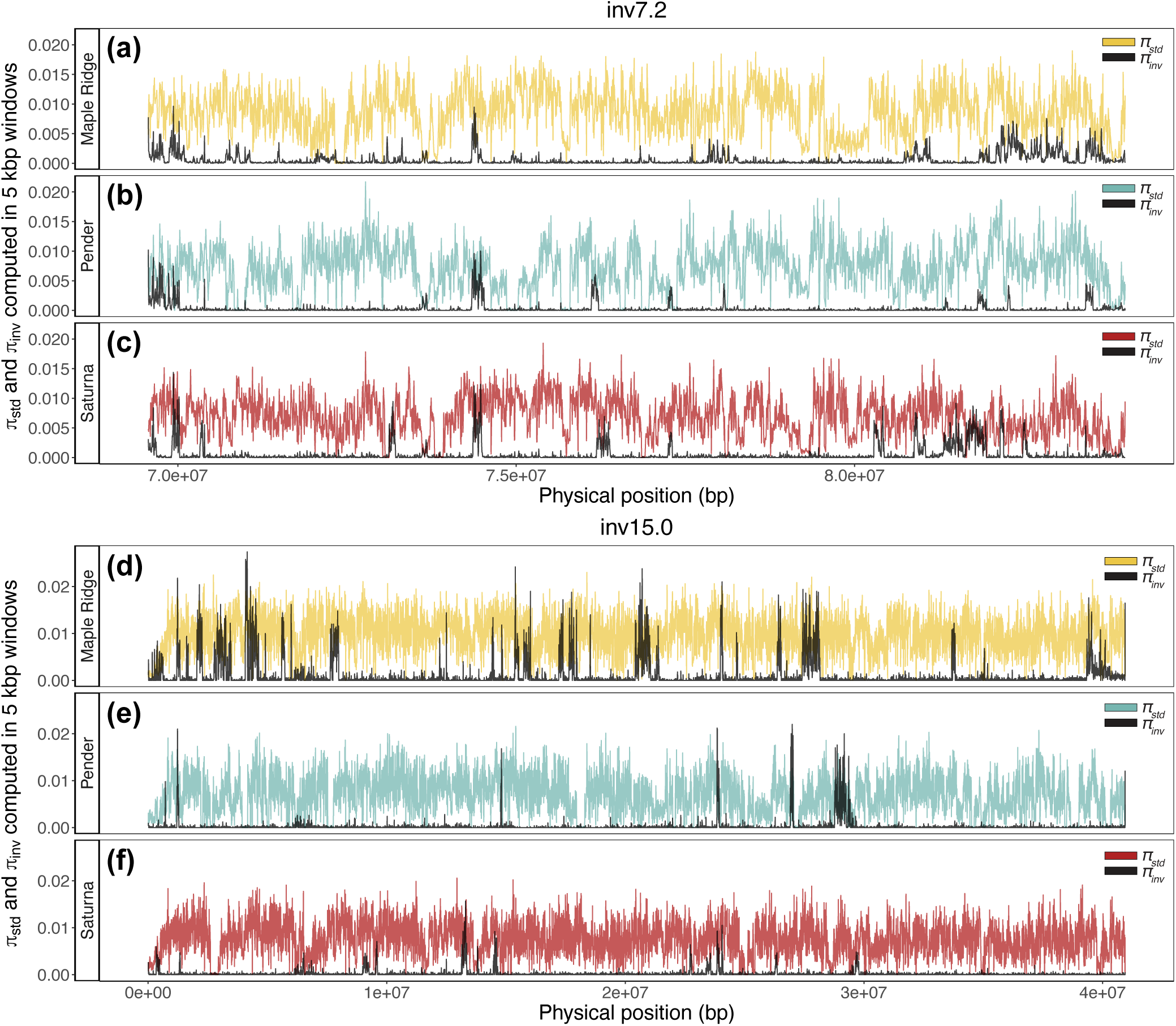
Polymorphic inversions on chromosomes 7 and 15 exhibit extreme reductions in nucleotide diversity. Horizontal panels compare nucleotide diversity among inverted karyotypes (𝜋_inv_) and nucleotide diversity among standard karyotypes (𝜋_std_) across the inv7.2 (a-c) and inv15.0 (d-f) loci. Each panel represents a different population: Maple Ridge (a and d), Pender Island (b and e), and Saturna Island (c and f). X-axes represent genomic position along the chromosome, bounded by inversion breakpoints. Y-axes plot each summary of diversity computed in non-overlapping 5 kbp windows. Yellow, blue, and red lines represent nucleotide diversity measured among standard karyotypes (𝜋_std_) sampled from Maple Ridge, Pender, and Saturna, respectively. Black lines represent nucleotide diversity within the corresponding sample of inverted karyotypes (𝜋_inv_).

### Demographic history

To provide context for the similar patterns of variation we observed at shared polymorphic inversions, we reconstructed the joint demographic history of the Saturna Island, Pender Island, and mainland Maple Ridge populations. Our aim was to estimate population split times, the timing and magnitude of population size changes, and migration rates between populations.

Using *Moments* (Jouganous et al. 2017), we fit a variety of simplified demographic models to joint SFS constructed from putatively neutral SNPs falling outside the boundaries of known inversion polymorphisms. Figure 8 depicts the best-fitting three-population demographic model. Maximum likelihood parameter estimates from this model suggest that ancestral island and mainland populations diverged 2,565,592 generations ago and were connected by a low rate of migration (m = 6.75e-6) until 5,090 generations ago (Figure 8). At this time, the contemporary Saturna Island and Pender Island populations split from their shared ancestor, and the mainland Maple Ridge population underwent an over eight-fold size reduction (Figure 8). Although this best-fit model provides evidence for migration between the contemporary Saturna Island and Pender Island populations (m = 1.63e-7), it does not support migration between the islands and mainland during this recent epoch (Figure 8). Effective population size (N_e_) estimates from this model suggest a large N_e_ for both the ancestral island (N_e_ = 255,958) and ancestral mainland (N_e_ = 579,644) populations and smaller N_e_ for contemporary populations (N_e_ = 18,607 for Saturna Island, 14,984 for Pender Island, and 69,313 for mainland Maple Ridge) (Figure 8).

**Figure 8.**
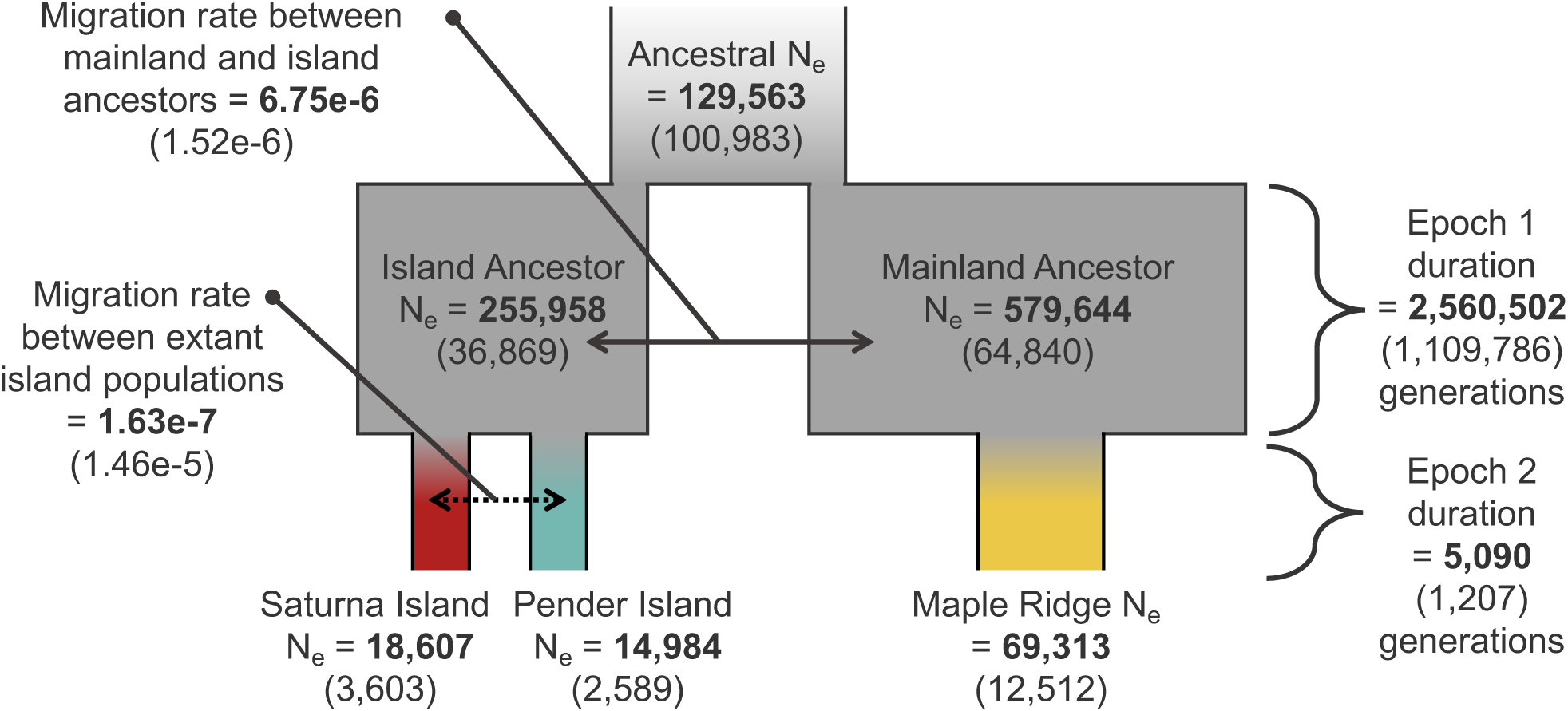
Parameter estimates for the candidate demographic model. Diagram reflects the structure of the best-fit three population demographic model. Boxes reflect population size changes through time (not drawn to scale). Gray shading reflects ancestral populations included in the model. Red, blue, and yellow shading reflects the contemporary Saturna Island, Pender Island, and mainland Maple Ridge populations, respectively. Arrows denote symmetric migration parameters included in the model. Maximum likelihood estimates for each parameter are bolded and standard deviations are given in parentheses. Standard deviations were obtained using the Godambe Information Matrix (Coffman et al. 2016) as an alternative to conventional bootstrapping.

Coalescent simulations conducted under this candidate model yielded a close fit to the means of empirical distributions of 𝜋, Tajima’s *D*, and F_ST_ (Supplementary Figure S13). Variance among loci in these summary statistic estimates is greater for the empirical data than for the simulated data, which could reflect heterogeneity in the recombination rate and mutation rate not captured by simulations. In contrast to the close agreement between empirical data and simulated data for diversity-related summaries, our best-fit demographic model underestimates levels of LD. In each population, average pairwise r^2^ decays slower than predicted by simulated data (Supplementary Figure S13). One explanation for this disagreement is that recent population size histories are more dynamic than suggested by our SFS-derived demographic model. To investigate this possibility, we used levels of LD to estimate contemporary N_e_, following the approach implemented in *currentNe* (Santiago et al. 2024). Our results indicated that contemporary N_e_ in each population is more than an order of magnitude smaller than suggested by our SFS-based model (N_e_ = 679 for Saturna Island, 219 for Pender Island, and 278 for mainland Maple Ridge) (Supplementary Table S1).

### Patterns of divergence at inversion loci mirror demographic history

Leveraging the shared history of the three Gulf Islands populations, we used patterns of divergence between populations to shed further light on the evolutionary relationships among inverted and standard karyotypes. We extended our comparisons of divergence between karyotypes sampled within the same population (*d_XY_*^karyotype^) to include three additional contrasts: inverted karyotypes sampled from different populations (*d_XY_*^inv^), standard karyotypes sampled from different populations (*d_XY_*^std^), and opposite karyotypes sampled from different populations (*d_XY_*^karyotype (between)^). We used these pairwise summaries of divergence as the basis for distance-based comparisons of inverted and standard karyotypes from all three populations. Figure 9 depicts the resulting trees constructed for each locus using the neighbor-joining algorithm of Saitou and Nei (1987).

**Figure 9.**
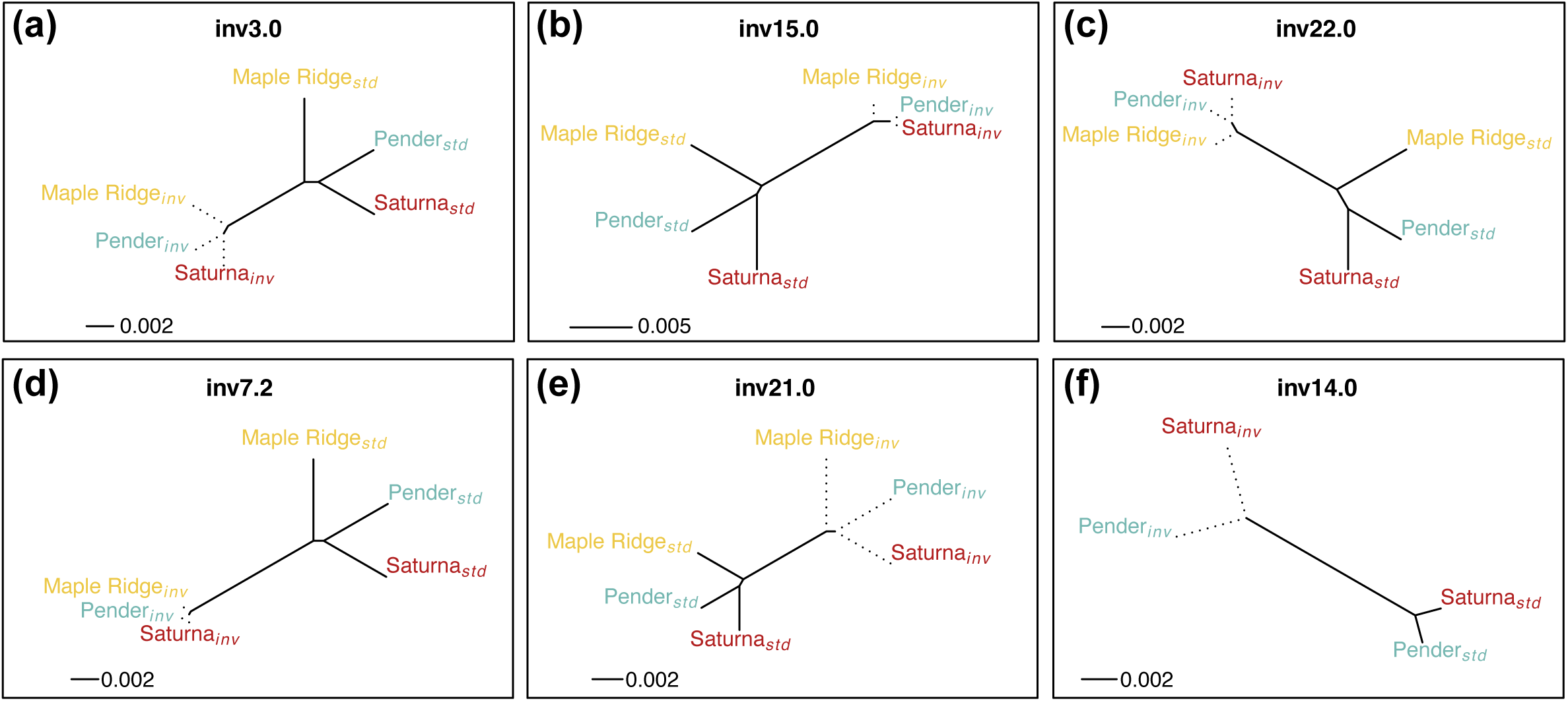
Divergence within and between karyotypes provide insights about the emergence of inversion polymorphisms. Panels depict unrooted, neighbor-joining trees relating standard and inverted karyotypes sampled from each population at each polymorphic locus: inv3.0 (a), inv15.0 (b), inv22.0 (c), inv7.2 (d), inv21.0 (e), and inv14.0 (f). Labels are colored according to population (red for Saturna Island, blue for Pender Island, and yellow for Maple Ridge) and subscripts denote karyotype. Dotted edges represent terminal branches for inverted karyotypes. Distance matrices were constructed using mean divergence (*d_XY_*) measured among karyotypic classes sampled within and between populations. Trees were constructed using the neighbor-joining algorithm of Saitou and Nei (1987). Source data is provided in Supplementary Table S5. Maple Ridge karyotypes are excluded from the inv14.0 tree (f) due to a lack of inversion homozygotes.

Surprisingly, despite the old split times we inferred between island and mainland lineages, karyotypes at all polymorphic loci cluster predominantly by orientation rather than by population (Figure 9). This suggests that inversions share a more recent common ancestor with inverted karyotypes in other populations than with their corresponding standard karyotype. Looking within either karyotypic class, branching patterns between populations mirror our best-fit demographic model (i.e., indicating a more recent split between the Saturna Island and Pender Island populations) (Figure 8 and Figure 9). Interestingly, there are major differences among loci in the relative branch lengths subtending inverted versus standard karyotypes. Whereas the inversions at inv3.0, inv15.0, inv22.0, and inv7.2 exhibit lower divergence among populations than their corresponding standard karyotype (Figure 9A-D), we find that the opposite pattern is true for inv21.0 and inv14.0, where inversions exhibit longer branch lengths than standard karyotypes (Figure 9E-F).

### Factors shaping divergence between karyotypes

Observed levels of divergence between standard and inverted karyotypes reflect the interplay of two opposing forces. Whereas the gradual accumulation of mutations increases divergence between them, rare “genetic flux” due to gene conversion and double crossover events within heterokaryotypic individuals erodes these differences (Navarro et al. 1997; Guerrero et al. 2012). In the absence of genomic heterogeneity in recombination, rates of genetic flux can depend on the length of the inversion and its historical frequency trajectory, both of which determine the propensity for exchange between karyotypes to occur. Longer inversions are more likely to experience the double crossover and gene conversion events that facilitate exchange between karyotypes. Similarly, inversions that have achieved higher frequencies will have spent a greater amount of time in the heterokaryotypic state, increasing the opportunity for genetic flux. Despite these expectations, *d_XY_*^karyotype^ is not strongly associated with inversion length or inversion frequency (Supplementary Figure S14; Supplementary Table S2).

In the absence of selection targeting regions located within inverted intervals, rates of genetic flux are expected to be greatest in the center of the inverted locus and lowest near the breakpoints, predicting a negative correlation between *d_XY_*^karyotype^ and distance from breakpoints (Navarro et al. 1997). Treating each inversion locus individually, we find little support for this trend. With the exception of a weak negative correlation at inv21.0 in the Saturna Island sample (Pearson correlation coefficient r = -0.15; p-value = 3.24e-3), all other significant correlations between *d_XY_*^karyotype^ and distance from breakpoints are weakly positive (Supplementary Figure S15; Supplementary Table S3).

## DISCUSSION

Our genomic characterization of *P. maniculatus* from the Gulf Islands revealed widespread effects of both polymorphic inversions and demographic history on patterns of variation within and between populations. In the following sections, we discuss how these results enrich our understanding of inversion evolution and the evolution of island populations in this unique system.

### A shared history for inversions segregating in island and mainland populations

Our profile of inversions in the Gulf Islands revealed surprising similarities between populations. Despite our estimates that ancestral island and mainland lineages diverged over 2.5 million generations ago, all three Gulf Islands populations segregate the same six inversion polymorphisms. These congruencies stand in stark contrast to the geographic variability among polymorphic inversions observed across mainland populations of *P. maniculatus* (Harringmeyer and Hoekstra 2022).

Inversion polymorphisms could be shared among populations because they were inherited from a common ancestor, dispersed by gene flow, or generated by distinct mutations at the same loci. Recent analyses of breakpoint content using long-read sequences have revealed an enrichment of segmental duplications and a lineage-specific centromeric satellite sequence flanking large-scale inversions in *P. maniculatus* (Gozashti et al. 2025). This observation, combined with evidence for breakpoint re-use in *P. maniculatus* (Gozashti et al. 2025), indicates that certain genomic regions are more susceptible to rearrangement than others, as has been observed in diverse species (Cáceres et al. 1999; Pevzner and Tesler 2003; Corbett-Detig et al. 2019; Maggiolini et al. 2020). Although the shared inversion polymorphisms in the Gulf Islands populations could have originated independently at such “fragile” sites, between-population comparisons revealed that inversions sampled from different populations have a lower time to most recent common ancestor (TMRCA) than opposite karyotypes sampled from the same population, indicating that they instead derive from single shared origins. Based on the demographic history we inferred, these inversions either arose in the ultimate island-mainland ancestor or were spread among lineages during the long period of connectivity between ancestral island and ancestral mainland populations. In addition, although we observed distinct trends across inversion loci, levels of within-karyotype diversity and between-karyotype divergence are broadly similar across all three populations within each locus, providing further support for a shared history.

### Differences among inversion loci

The distinct patterns of diversity and divergence we observed across inversion loci could reflect differences in the selective regime acting on inversions (discussed in the next section), differences in inversion age, or a combination of both properties. Although levels of divergence between karyotypes can provide useful insights about the age of an inversion (Hasson and Eanes 1996; Corbett-Detig and Hartl 2012), direct estimation of inversion age is complicated by genetic flux between karyotypes and departures from demographic equilibrium (Hasson and Eanes 1996; Corbett-Detig and Hartl 2012; Charlesworth 2023). Surprisingly, we found that inversions at loci with the highest levels of between-karyotype divergence, inv15.0 and inv7.2 (Figure 6), also exhibit the lowest levels of divergence across Gulf Islands populations, as illustrated by the shorter branch lengths in Figure 9. In contrast, inv21.0 exhibits the lowest levels of between-karyotype divergence (Figure 6), yet inversions at this locus show high levels of divergence across populations (Figure 9). Though these within-population and between-population patterns appear to paint contrasting portraits of inversion age, they likely reflect the complex interaction between rates of genetic flux, inversion age, and selection. Future efforts aimed at jointly inferring the strength of selection, rates of genetic flux, and inversion age under explicit demographic scenarios will enable clearer insights about inversion dynamics.

### Evidence for selection on inversions

Multiple findings suggest that the inversions segregating in Gulf Islands populations have been maintained longer than expected under neutrality. First, all six inversion polymorphisms are shared by the three Gulf Islands populations. Second, whole genome alignments of diverse *Peromyscus* reference assemblies suggest at least two inversions, inv14.0 and inv22.0, may pre-date species splits. Finally, levels of divergence between karyotypes far exceed genome-wide levels of diversity measured at putatively neutral, non-inverted loci, consistent with an elevated TMRCA.

Models of long-term balancing selection predict an unusually old TMRCA at the targeted locus and a genealogy characterized by elongated internal branches (Takahata 1990; Hudson 1990; Charlesworth et al. 1997; Bitarello et al. 2023). These features lead to increased neutral variation at the locus, elevated linkage disequilibrium between neutral variants and the selected site, and shifts in the site frequency spectrum toward intermediate-frequency alleles (Bitarello et al 2023; Charlesworth 2023). In most genomic regions, recombination narrows the span of these signals, challenging efforts to detect balancing selection. In regions harboring polymorphic inversions, by contrast, recombination suppression in heterokaryotypes extends the signatures of balancing selection over much greater distances (Charlesworth 2023). At the same time, genetic flux between karyotypes can cause patterns of neutral diversity to vary with distance from breakpoints and selected sites (Hasson and Eanes 1996; Navarro et al. 1997; Navarro et al. 2000; Corbett-Detig and Hartl 2012; Guerrero et al. 2012; Charlesworth 2023; Kapun et al. 2023).

Population genetic models that account for some of these complexities have provided insights into the effects of balancing selection on inversion loci. Navarro et al. (2000), Guerrero et al. (2012), and Charlesworth (2023) studied the behavior of coalescence times within and between karyotypes under different models of balancing selection and population structure. Their findings indicated that expected patterns of diversity and divergence strongly depend on whether the system has reached equilibrium and the equilibrium frequency at which the inversion is maintained (Navarro et al. 2000; Guerrero et al. 2012; Charlesworth 2023). After a selectively favored inversion arises, its approach to equilibrium occurs in two phases (Navarro et al. 2000; Guerrero et al. 2012; Charlesworth 2023). First, the inversion quickly rises to its balanced frequency, eliminating variation within the inverted class in a manner analogous to a selective sweep (Maynard Smith and Haigh 1974). Once this balanced frequency is reached, variation within the inverted class recovers through the accumulation of new mutations and genetic flux with the standard karyotype. In a symmetric fashion, variation within the standard class is lost because of its reduced frequency in the population. After equilibrium is established, relative levels of diversity within each karyotypic class should reflect the frequencies at which they are maintained.

We observed surprising variability among loci in relative levels of nucleotide diversity within inverted karyotypes. Whereas inversions at most loci (inv3.0, inv7.2, inv15.0, and inv22.0) exhibit reductions in diversity, inversions at two loci (inv14.0 and inv21.0) exhibit higher nucleotide diversity than the standard karyotype. In keeping with theoretical expectations, these contrasting levels of relative nucleotide diversity could reflect the long-term equilibrium frequencies at which each inversion has been maintained. Given the dynamic population size histories we infer, the equilibrium status of these loci among the Gulf Islands populations is uncertain. However, looking across loci, relative levels of 𝜋_inv_ do not obviously match the population-specific or continent-wide frequencies of inverted karyotypes (Table 1).

Alternatively, reductions in relative 𝜋_inv_ could reflect recent selective sweeps of inversions to their balanced frequencies. We found that inv15.0 and inv7.2 exhibit marked reductions in diversity within the inverted karyotype that span nearly the entire locus (Figure 7). Though striking, the extent to which these patterns reflect selective sweeps is unclear. Such broad reductions in diversity would require extremely strong selection, such that there has been little opportunity for recombination within inversion homokaryotypes to narrow the sweep signature. Increases in relative 𝜋_inv_ for inv14.0 and inv21.0 also seem inconsistent with neutral evolution, but are difficult to reconcile with existing theory. Current models assume that selection on inversions is sufficiently strong that their equilibrium frequency is quickly achieved, and that the selective regime remains constant through time (Navarro et al. 2000; Guerrero et al. 2012; Charlesworth 2023). In reality, the dynamics could depend on both the mechanism of balancing selection (Fijarczyk and Babik 2015; Bitarello et al. 2023) and its targets (inversion breakpoints or alleles captured by the rearrangement) (Said et al. 2018; Kapun et al. 2023), features that may differ from inversion to inversion. Alternatively, the trends we observed at these two loci could reflect errors in our classification of homozygous genotypes or in our determination of ancestral states. The accuracy of these assignments is influenced by factors including the age of the inversion, the quality of reference assemblies, and the probability of incomplete lineage sorting.

More broadly, our findings suggest that there could be important regional differences in the role these inversions play in local adaptation. The same inversions we examined show strong evidence for selection in mainland populations of *P. maniculatus* from the Cascades region, where they exhibit clines across a forest-prairie gradient and associations with phenotypic differences between forest and prairie ecotypes (Hager et al. 2022; Harringmeyer and Hoekstra 2022). In contrast to their role in ecotype differentiation among Cascades populations, none of the inversion loci exhibit allele frequency differentiation among Gulf Islands populations, nor do their frequencies in the Gulf Islands mirror continent-wide averages measured by Harringmeyer and Hoekstra (2022) (Table 1). Furthermore, the joint minor allele frequencies of the six inversion polymorphisms across populations (computed by treating the loci as a group) are not unusual with respect to the joint frequencies of neutral, non-inverted SNPs (Supplementary Figure S16).

Important differences between the Gulf Islands and Cascades could lead to contrasting evolutionary roles for the same inversions in these two systems. Although island and mainland environments may present divergent selective regimes, the selective gradient responsible for establishing inversion clines among Cascades populations might not exist in the Gulf Islands. In addition, whereas forest and prairie populations in the Cascades region exhibit high rates of migration across divergent environments (Hager et al. 2022), we found little support for migration between island and mainland populations during the contemporary epoch of our demographic model. By linking together adaptive allele combinations, inversions may facilitate local adaptation in the face of high gene flow (Kirkpatrick and Barton 2005; Yeaman and Otto 2011; Yeaman and Whitlock 2011). Though this phenomenon may position inversions as an important adaptive mechanism in the Cascades region, our demographic inferences indicate that gene flow is not an impediment to adaptive divergence between island and mainland populations in the Gulf Islands. Hager et al. (2022) suggest that inv15.0 pre-dates formation of the current forest and prairie habitat distribution, raising the possibility that its role in local adaptation has resulted from subsequent accumulation of adaptive alleles (Said et al. 2018; Kapun et al. 2023). Future comparisons of the genetic variation carried by these inversions in the Gulf Islands, Cascades, and other populations of *P. maniculatus* could help clarify their roles in local adaptation.

### Prospects for inversion characterization

Our work highlights the valuable insights that can be gained by interpreting patterns of variation spanning inversion polymorphisms within the context of an explicit demographic model. At the same time, we illustrate how the unique evolutionary dynamics experienced by inversions generate characteristic departures in allele frequencies and LD that could have severe consequences for reconstructing demographic history. Indeed, recent demographic analyses using genomic data from the inversion-rich spruce bark beetle demonstrated that segregating inversions affect inferences derived from both SFS and sequentially Markovian coalescent (SMC)-based frameworks depending on the relative contributions of co-linear and non-colinear sequences to the data used for analysis (Zieliński et al. 2025). The six inversions we observed segregating in Gulf Islands *P. maniculatus* span a large portion of the autosomal genome (∼6%), have accumulated substantial divergence between karyotypes, and are shared across populations, positioning them in a zone of parameter space with great potential to bias demographic inference. Fortunately, these same properties that are likely to mislead demographic inferences also make it straightforward to detect segregating inversions based on the distortions they produce in the SFS and patterns of LD. Our findings demonstrate that partitioning SNP genotypes by their minor allele frequency provides a simple framework for detecting and genotyping inversions in the absence of long-read sequencing data.

Despite recent progress in the characterization of *P. maniculatus* inversions (Harringmeyer and Hoekstra 2022; Gozashti et al. 2025), much remains to be uncovered about inversion evolution in this system. Knowledge of inversion polymorphisms in *P. maniculatus* and the broader karyotypic diversity of the genus pre-dates the genomic era (Ohno et al 1966; Robbins and Baker 1981; Stangl and Baker 1984; Greenbaum et al. 1994), yet important biological properties of these loci are unknown. Uncertainty about local mutation rates and rates of genetic flux between karyotypes preclude more detailed model-based inferences about inversion age and selection. Estimating these key parameters is an important next step for extracting deeper insights about inversion dynamics in *P. maniculatus*.

### Insights into the evolution of island populations

The demographic history we reconstructed from genome-wide patterns of variation provides new insights into the population dynamics of *P. maniculatus*. Assuming 1-2 generations per year (Wolff 1985), the estimated split time of 5,090 generations between Pender Island and Saturna Island is broadly consistent with the post-glacial establishment of these populations. During the last glacial period, the southern Strait of Georgia and Gulf Islands region was covered by the Cordilleran ice sheet (Clague 1980; Clague and James 2002; Hutchinson et al. 2004; Fedje et al. 2009; James et al. 2009). Following its peak extent 14,500 years ago, the retreat of glacial ice drove rapid uplift of underlying land areas, with relative sea level in the southern Strait of Georgia reaching its lowest point around 11,500 years ago (James et al. 2009). Subsequent rises in global sea level driven by glacial melt established the present-day distribution of land area in the region by 6,000 years ago (James et al. 2009). There is conflicting evidence about the extent of land exposure between southeastern Vancouver Island and mainland British Columbia during this period of sea-level adjustment (Gowan 2005; James et al. 2009), leaving unclear whether mainland *P. maniculatus* could have dispersed to the islands via emergent land bridges. In the absence of such overland routes, it is possible that *P. maniculatus* may have dispersed from mainland areas by rafting on floating ice or debris (Steffan 2016). Notably, our island divergence time estimate also post-dates human inhabitation of coastal British Columbia (Mackie et al. 2011; Mackie et al. 2018), raising the possibility that island populations could have instead been founded by inadvertent human introductions.

Given its closer proximity and greater connectivity to the Gulf Islands during the post-glacial sea level low stand, Vancouver Island may be a more likely source for the Saturna Island and Pender Island populations than mainland British Columbia. Radiocarbon dating of *Peromyscus* fossils from northern Vancouver Island suggest that *P. maniculatus* could have been present as early as 14,000 years ago (Steffan 2016). As this estimate pre-dates deglaciation of the region (Clague and James 2002; Hutchinson et al. 2004; Fedje et al. 2009; James et al. 2009), it indicates that Vancouver Island *Peromyscus* fossils might reflect dispersals from nearby glacial refugia (Steffan 2016). Population genetic analysis of *P. maniculatus* from Vancouver Island and other potential source populations could increase the resolution of demographic inferences for mice from Saturna Island and Pender Island.

The ancient divergence time between ancestral island and ancestral mainland populations we recovered pre-dates the last inter-glacial period substantially (Clague 1980), leaving its connection to the geological history of the region unclear. Though there is phylogeographic evidence for deep divergence among certain geographic clades of *P. maniculatus* (Sawyer et al. 2017), our results are surprising given the proximity of the island and mainland sites we sampled. Demographic analysis of *P. maniculatus* from across the Cascade mountain range produced similar divergence time estimates (2.2 million generations ago) between western populations of the forest ecotype and eastern prairie-ecotype populations (Hager et al. 2022). The similarity in divergence times and the broad proximity of the Gulf Islands and Cascades regions suggest that island-mainland divergence and forest-prairie divergence in these two systems were driven by shared geographic features. Indeed, the Fraser River, which bisects south central British Columbia and northern Washington along its eastern margin, could have shaped population structure among *P. maniculatus* in the Pacific Northwest (Zheng et al. 2003).

Our results also speak to demographic processes operating more recently. Analyses of LD in the Gulf Islands populations indicated that values of contemporary N_e_ are 1-2 orders of magnitude lower than historical estimates based on the SFS. The causes of these recent, severe declines in population size are unknown, but they suggest a reduced capacity for selection to remove deleterious mutations and to spread beneficial mutations in these populations. In addition, our findings serve as a reminder that different aspects of genomic variation respond to evolutionary processes happening on disparate timescales (Beichman et al. 2018; Nadachowska-Brzyska 2022). We found a similar discordance between N_e_ estimates based on allele frequencies and patterns of linkage disequilibrium in island populations of the white-footed mouse, *P. leucopus* (Howell et al. 2025). Commonalities between these systems, such as the fragmentation and urbanization of habitat areas (Eis and Craigdallie 1980; Richburg and Patterson 2005), might suggest that fluctuations in effective population size on contemporary timescales are driven by shared processes. Similar comparisons in natural populations of other species could reveal the extent to which these disparities are a universal reflection of human activity.

Finally, our inferred demographic history provides context for interpreting patterns of phenotypic evolution on islands. Mice on Saturna Island exhibit multiple hallmarks of the “island syndrome” (Adler and Levins 1994), including larger body sizes, higher population densities, lower dispersal, and reduced intraspecific aggression (Redfield 1976; Sullivan 1977; Halpin and Sullivan 1978). Saturna Island mice are 35% heavier than mainland mice when raised in a common laboratory environment, indicating a heritable difference in body size (Baier and Hoekstra 2019). The ancient split time between island and mainland lineages substantially extends the temporal window during which the mice that now inhabit Saturna Island and Pender Island could have evolved island syndrome phenotypes. Comparisons of molar sizes between ancient and modern specimens suggest that Pleistocene-era *P. maniculatus* from Vancouver Island were significantly larger than their modern counterparts (Steffan 2016). This inference raises the intriguing possibility that the large body sizes of Saturna Island and Pender Island mice (Redfield 1976; Sullivan 1977; Baier and Hoekstra 2019) are relics of this ancient characteristic.

## MATERIALS AND METHODS

### Fieldwork and sample collection

We sampled deer mice (n=120) in the Gulf Island National Park Reserve on Saturna Island and Pender Island (*P. m. saturatus*) and in the Malcolm Knapp Research Forest (MKRF) on the mainland (*P. m. austerus and P. keeni*) in October 2014. Of these, we collected 101 individuals, and either sacrificed and froze them in the field or transported them alive to Harvard University. Specimens were accessioned in the Harvard Museum of Comparative Zoology (MCZ 69713– 69813) and liver samples were stored in ethanol. We verified species identity of specimens using a fragment of the cytochrome b gene following Zheng et al. (2003) (see electronic supplementary material, Figure S1, in Baier and Hoekstra (2019) for details). The samples in the present study were previously examined by Baier and Hoekstra (2019). We sequenced 79 of the 101 mice for which liver tissue was collected (n=31 for Saturna Island, n=16 for Pender Island, and n=32 for mainland Maple Ridge). Fieldwork was approved by the Harvard University Faculty of Arts and Sciences Institutional Animal Care and Use Committee (protocol #14-08-211). Fieldwork was conducted with permits from the Gulf Island National Park Reserve (GINP-2014-17276), the Pender Island Parks and Recreation Commission, the Malcolm Knapp Research Forest at the University of British Columbia and the Ministry of Forests, Lands and Natural Resource Operations (NASU14-154228, NASU-154230).

### Population sampling and sequencing

DNA was extracted from liver samples using Gentra Puregene Tissue Kits (Qiagen, Germantown, MD, USA). Sample purity was assessed by measuring 260/280 absorbance ratios using a NanoDrop Spectrophotometer (Thermo Fisher Scientific, Waltham, MA, USA). DNA concentrations were measured using the Qubit dsDNA HS Assay (Life Technologies, Carlsbad, CA, USA). We used agarose gel electrophoresis to assess DNA sample quality and confirm the presence of high-molecular weight DNA prior to sequencing. Library preparation and sequencing were conducted by the University of Wisconsin-Madison Biotechnology Center. The Illumina DNA Library Prep Kit was used to construct sequencing libraries. Purified and pooled libraries were sequenced on the Illumina NovaSeq X Plus platform to generate paired-end 150 bp reads. Samples were sequenced to a targeted coverage of 30X per individual. As a quality control measure, we re-sequenced four mice: one male each from Saturna (FSB85), Pender (FSB114), and Maple Ridge (FSB42) as well as a single female from Maple Ridge (FSB6). These technical replicates were sequenced using unique libraries, allowing us to capture the effects of sequencing/genotyping error introduced at all laboratory and bioinformatic steps downstream of PCR-based library preparation.

### Sequence data processing, variant calling, and variant filtering

Prior to alignment and variant calling, raw sequencing reads were pre-processed to remove adapter sequences and low-quality base calls. BBDuk v38.96 (https://sourceforge.net/projects/bbmap/) was used to remove Illumina universal adapter sequences and Trimmomatic v0.39 (Bolger et al. 2014) was used to trim low-quality base calls. Processed sequencing reads were then aligned to the *P. maniculatus* NCBI RefSeq assembly (HU_Pman_2.1.3; GCF_003704035.1) using BWA-MEM v0.7.17 (Li and Durbin 2009).

We based our variant calling approach on the Genome Analysis Toolkit (GATK) Best Practices recommendations for germline short variant discovery (McKenna et al. 2010; Van der Auwera and O’Connor 2020). Prior to variant identification, we used GATK v4.2.0.0 MarkDuplicates to identify and exclude duplicate reads from downstream variant calling. We then computed genotype likelihoods on a per-individual basis using GATK v4.2.0.0 HaplotypeCaller with the Reference Confidence Model set to “GVCF” mode (Poplin et al. 2017). The resulting GVCF call sets were combined using GATK v4.2.0.0 CombineGVCFs. GATK v4.2.0.0 GenotypeGVCFs was then used to perform joint variant calling separately for the Saturna (n=32), Pender (n=17), and Maple Ridge (n=34) cohorts (Poplin et al. 2017). This approach identified a total of 110,458,672 SNPs in the Saturna cohort, 100,102,299 SNPs in the Pender cohort, and 151,071,903 SNPs in the Maple Ridge cohort.

We filtered our variant calls with GATK v4.2.0.0 VariantFiltration using site-level annotation thresholds based on GATK Best Practices recommendations for hard filtering (“FisherStrand” > 60.0, “StrandOddsRatio” > 3.0, “RMSMappingQuality” < 40.0, “MappingQualityRankSumTest” < -12.5, and “ReadPosRankSumTest” < -8.0; Van der Auwera and O’Connor 2020) with modifications (“QualByDepth” < 5.0). We then used GATK v4.2.0.0 SelectVariants to remove filtered sites and restrict each call set to autosomal, biallelic, single-nucleotide variants. We retained 78,214,290 SNPs in the Saturna cohort, 72,836,785 SNPs in Pender, and 106,361,479 SNPs in Maple Ridge following these basic variant filtering and selection procedures.

The population-specific joint calling approach we employed can yield incongruent call sets when there is sufficient allele frequency differentiation between populations because variants private to one cohort will not be represented among the variant calls of other cohorts. Variants that are “missing” between call sets could be monomorphic for the reference allele within their cohort, or they could be sites for which high-quality genotype calls could not be obtained. To avoid this ambiguity and ensure that joint allele frequencies were properly calibrated for downstream analyses, we “rescued” genotypes at these missing sites. Briefly, we used the bcftools v1.8 (Danecek et al. 2021) “isec” function to extract the locations of private variants observed in each call set. We then used the “-all-sites” mode of GATK v4.2.0.0 GenotypeGVCFs with the “- stand-call-conf” threshold set to 0 to emit monomorphic reference calls at these private variant sites in each of the other cohorts. We used GATK v4.2.0.0 VariantFiltration and SelectVariants to apply the same site-level filtering thresholds to these “rescued” calls (excluding “QualByDepth”). The resulting high-quality invariant calls were then combined with the polymorphic calls of their respective cohort and resulting monomorphic + polymorphic call sets were merged across cohorts, yielding a single multi-population call set that represented the union of variant sites observed across the three populations.

### Identification and exclusion of close relatives

Using KING v2.2.8 (Manichaikul et al. 2010), we estimated pairwise kinship coefficients among individuals within each population cohort. Prior to kinship analysis, each of the filtered single-population autosomal call sets described above was pruned for LD by removing SNPs with pairwise r^2^ > 0.1 in 50 bp sliding windows using the “--indep-pairwise 50 10 0.1” option in plink v1.90 (Chang et al. 2015). This LD-pruning left a total of 2,853,858 SNPs in the Saturna cohort, 2,079,267 in Pender, and 5,503,047 in Maple Ridge. We applied the KING-Robust algorithm to each LD-pruned call set and determined relationships from the resulting kinship coefficient estimates using the bounds suggested by the KING documentation (Manichaikul et al. 2010).

We identified a large number of closely related individuals within all three cohorts, with 4.44% of Saturna Island pairs, 10.29% of Pender Island pairs, and 6.95% of mainland Maple Ridge pairs exceeding the lower bound for 2^nd^ degree relatives. We did not find close relatives among individuals sampled from different locations. To mitigate the potential biases that close relatives can introduced to population genetic analyses (Wang 2017), we excluded related individuals from downstream analyses. To do so, we used the “relatednessFilter” function of the R package plinkQC v0.3.4 to identify the maximum set of unrelated individuals (defined here as pairs ≥ 3^rd^ degree relatives) within each sample. This left us with unrelated sample sizes of n=20 for Saturna, n=10 for Pender, and n=17 for Maple Ridge.

### Population structure analysis

We evaluated population structure within and between sampling locations by assessing PCA clustering patterns and conducting model-based inference of global ancestry proportions with ADMIXTURE (Alexander et al. 2009). Since both approaches require unlinked SNPs, we conducted the same LD-pruning described above on the filtered multi-population autosomal call set, restricting our analyses to the unrelated subset of individuals identified in each cohort. PCA was performed on this combined call set using the “--pca” option in plink v1.90 (Chang et al. 2015). Ancestry proportions were inferred either for each population sample separately or for the combined cohort with ADMIXTURE v1.3.0 (Alexander et al. 2009). Five-fold cross-validation was performed for a range of ancestral components (k=1 to k=5) using the “--cv” option. Individual ancestry proportions were then inferred for the k value that produced the lowest cross-validation error.

### Inversion identification

We used GATK v4.2.0.0 VariantsToTable to extract information about position, allele count, allele number, allelic states, and individual genotypes at each SNP in the filtered autosomal call sets described above. The resulting genotype tables were used to inspect properties of the genome-wide folded SFS within each population. To investigate the source of “bumps” observed in each population’s SFS, we partitioned genotype tables by chromosome in order inspect chromosome-wide allele frequencies. To assess LD decay along each chromosome, we used the “--r2” option in plink v1.90 (Chang et al. 2015) to measure pairwise r^2^ among SNPs sampled from each chromosome (set by the “--chr” argument) in the filtered VCF call sets. Singleton variants and sites with missing data were excluded from these LD calculations by setting the “-- mac” argument to 1 and the “--geno” argument to 0.

In order to “screen” each population sample for evidence of segregating inversions, we looked for intervals that exhibited a high density of either homozygous or heterozygous SNPs in each individual at a specific MAF, consistent with an excess of fixed differences between inverted and standard karyotypes. To perform this screen, we partitioned each population’s genotype table by chromosome and by every possible MAF. Coloring SNPs based on their genotype in each individual, we then constructed maps of these MAF-partitioned SNPs and assessed whether regions exhibiting high SNP density and genotype continuity overlapped the boundaries of previously characterized inversion polymorphisms (Harringmeyer and Hoekstra 2022). MAF-partitioned genotype maps were constructed and visualized using ggplot v3.5.1 (Wickham 2016) in R v4.2.1 (R Core Team 2022).

To confirm the presence of the inversion polymorphisms identified by our screen and to determine whether additional loci may be segregating in any of the Gulf Islands populations, we conducted PCA within each of the 21 inversion loci characterized by Harringmeyer and Hoekstra (2022). Starting with the filtered multi-population call set, we used the “--extract” function in plink v1.90 (Chang et al. 2015) to extract SNPs falling within each inversion’s breakpoints, then performed PCA on each subset of SNPs using the “--pca” function. Because segregating loci carry an abundance of fixed differences that distinguish standard and inverted karyotypes, we did not prune SNPs within inversion intervals for LD prior to analysis. For comparison, we also conducted PCA on genome-wide SNPs falling outside of known inversion intervals. This subset of SNPs was pruned for LD using the same parameters described previously. For each locus, we evaluated whether PCA clustering patterns were consistent with three genotypic classes: homozygous ALT, heterozygous, and homozygous REF. Though inv7.3 exhibits patterns consistent with a segregating inversion (Supplementary Figure S4), we note that this is likely driven by polymorphism at the larger inv7.2 locus in which it is nested, and as such do not treat it as a separate polymorphism.

For each polymorphic locus, we assigned individuals to one of three genotypic classes (homozygous ALT, heterozygous, and homozygous REF) by inspecting each mouse’s genotype proportions for SNPs segregating at the MAF of the rearrangement. We reasoned that mice with an enrichment of homozygous reference (REF/REF) genotypes carry the same orientation as the reference assembly while those enriched for homozygous alternate (ALT/ALT) genotypes carry the opposite orientation. For inv3.0, inv7.2, inv14.0, inv15.0, and inv22.0, the homozygous classes are clearly distinguished by their proportion of ALT/ALT genotypes (Supplementary Figure S9), and these assignments are visually apparent among MAF-partitioned genotypes (Figure 4 and Supplementary Figures S2-S3). In contrast, neither homozygous class shows a consistent enrichment of ALT/ALT genotypes across all three populations at the inv21.0 locus (Supplementary Figure S9). Instead, one of the homozygous classes (hereby assumed to carry the alternate orientation) shows an enrichment of ALT alleles at MAFs approaching that of the rearrangement (Supplementary Figures S10-S12). This pattern might indicate that although private variants have achieved high frequencies within either karyotypic subpopulation, the inversion is sufficiently young that few of these private differences have fixed. Alternatively, this lack of fixed differences could indicate that the inversion is sufficiently old such that substantial genetic exchange has occurred between karyotypes, impeding the fixation of private mutations. Though we find little evidence for a major role of genetic flux in shaping patterns of divergence between karyotypes (see Results), distinguishing between these explanations remains difficult given our lack of knowledge about rates of exchange.

### Inversion polarization

In order to connect the genotype assignments described above (homozygous REF, heterozygous, and homozygous ALT) to rearrangement karyotype in each individual (inv/inv, inv/std, and std/std), we polarized segregating inversion loci to determine whether the *P. maniculatus* assembly carries the inverted (i.e., derived) or standard (i.e., ancestral) orientation at each locus. To assess evolutionary relationships among existing *Peromyscus* reference genomes, we ran Mashtree v 1.4.6 (Katz et al. 2019) on all of the existing *Peromyscus* NCBI RefSeq assemblies: *P. maniculatus* (HU_Pman_2.1.3; GCF_003704035.1), *P. polionotus* (HU_Ppol_1.3.3; GCA_003704135.2), *P. leucopus* (UCI_PerLeu_2.1; GCF_004664715.2), *P. californicus* (ASM782708v3; GCF_007827085.1), *P. eremicus* (PerEre_H2_v1; GCF_949786415.1), *P. nudipes* (Pnud_10x_v1; GCA_902168325.1), *P. melanophrys* (Pmel_10x_v1; GCA_902168415.1), *P. attwateri* (Patt_10x_v1; GCA_902168425.1), and *P. aztecus* (Pazt_10x_v1; GCA_902168405.1). The resulting distance-based tree (Supplementary Figure S17) recovers previously characterized phylogenetic relationships by Bradley et al. (2007) and Platt et al. (2015). We focused our polarization analysis on four of these species: *P. polionotus*, *P. leucopus*, *P. californicus*, and *P. eremicus*, which collectively span 1.5 to 5 million years of divergence from *P. maniculatus* (Bradley et al. 2007; Platt et al. 2015).

We used the nucmer module of the MUMmer v4.0.0 alignment package (Marçais et al. 2018) to conduct pairwise, whole genome alignments between *P. maniculatus* and each of the four focal *Peromyscus* species. Each pair of reference assemblies was aligned using the following parameters: “--maxmatch”, “-c 500”, “-b 500”, and “-l 100”. The resulting alignments were filtered using the delta-filter module with parameters “-m”, “-i 90”, and “-l 100”. We then used the show-coords module with parameters “-THrd” to summarize the filtered alignments.

Supplementary Figures S18-S21 depict the resulting dotplots from our initial whole genome alignments. Though the closely related *P. polionotus* assembly exhibits one-to-one homology at the chromosome level (Supplementary Figure S18), we identified major inconsistencies between chromosome name and order between *P. maniculatus* and the more diverged *P. leucopus* (Supplementary Figure S19) and *P. eremicus* assemblies (Supplementary Figure S20). Both the *P. leucopus* and *P. eremicus* assemblies support the fusion of chromosomes 16 and 21 into a single scaffold (chr16_21) and the splitting of chromosome 8 into two separate scaffolds (chr8a and chr8b). According to Long et al. (2019), chromosome assignments for the *P. leucopus* reference were based on the mapping of markers from the *P. maniculatus* linkage map (Kenney-Hunt et al. 2014; Brown et al. 2018) and based on the results of their Hi-C contact analysis, which supported two separate chromosome 8 scaffolds and a fused chromosome 16 and chromosome 21 scaffold. Neither of these fusions or splits are reflected in the *P. maniculatus* assembly. Instead, we found that chr16_21 in both *P. eremicus* and *P. leucopus* is homologous with chr21 in *P. maniculatus* (Supplementary Figures S20 and S19). For the split chromosome 8, we find that chr8a and chr8b in *P. eremicus* align with chr8 and chr16 in *P. maniculatus*, respectively (Supplementary Figure S20). In contrast, chr8a and chr8b in *P. leucopus* align with chr16 and chr8 in *P. maniculatus* (Supplementary Figure S19). We also find that the assignments of chr11 and chr15 in both *P. eremicus* and *P. leucopus* are switched in *P. maniculatus* (Supplementary Figures S20 and S19). We used the “replace” function in seqkit v2.9.0 (Shen et al. 2016) to re-assign chromosome names in both *P. eremicus* and *P. leucopus* reference FASTAs to reflect their homology with *P. maniculatus*. For the scaffold-level *P. californicus* assembly FASTA, we assigned names to the 24 largest scaffolds to reflect their alignment to *P. maniculatus* chromosomes (Supplementary Figure S21).

In addition to these chromosome naming inconsistencies, we found that pairwise alignments with *P. leucopus*, *P. eremicus*, and *P. californicus* all exhibit reverse-strand errors in which homologous chromosomes in each pair are represented by the forward strand in one species and the reverse strand in the other (Supplementary Figures S19-S21). These errors manifest as whole-chromosome inversions and can confound the identification of intra-chromosomal rearrangements. To correct these errors, we used the “seq -r -p” option in seqkit v2.9.0 (Shen et al. 2016) to reverse-complement chr1, chr2, chr5, chr8, chr9, chr13, chr16, chr17, chr22, and chrX in the alignment with *P. californicus* and chr1, chr5, chr9, chr11, chr12, chr17, chr18, chr20, and chrX in the alignments with both *P. eremicus* and *P. leucopus*.

Following these operations on the reference FASTAs, we re-ran each alignment with the same MUMmer parameters described above. We used SyRI v1.7.1 (Goel et al. 2019) to identify large-scale structural rearrangements between species from these revised alignments and visualized the resulting synteny plots using plotsr v1.1.1 (Goel and Schneeberger 2022). Using established phylogenetic relationships among these species (Bradley et al. 2007; Platt et al. 2015), we then assessed collinearity across inversion intervals to determine the orientation (i.e., standard versus inverted) of each inversion polymorphism the *P. maniculatus* reference genome.

### Population genomic analyses of inversions

Taken together, the genotype assignments described above, which reflect the status of each arrangement relative to the *P. maniculatus* reference assembly (i.e., homozygous ALT, heterozygous, or homozygous REF), and our polarization results, which establish the orientation of each locus in the reference, enabled us to determine the inversion genotype carried by each mouse (i.e., inv/inv, inv/std, or std/std). These designations are summarized in Supplementary Table S4. To avoid switch errors that could be introduced by the phasing of inverted and standard haplotypes in heterozygous individuals, we restricted our analyses of diversity and divergence to the two homozygous classes of individuals (inv/inv and std/std).

Using these homozygous subsets, we computed nucleotide diversity π (Tajima 1983) within each karyotype (π_inv_ and π_std_) and divergence *d_XY_* (Nei and Li 1979) measured either between opposite karyotypes sampled from the same population (*d_XY_*^karyotype^), between populations within each karyotypic class (*d_XY_*^inv^ and *d_XY_*^std^), or between opposite karyotypes sampled from different populations (*d_XY_*^karyotype (between)^). We used scikit-allel v1.3.6 (https://github.com/cggh/scikit-allel) to compute each of these summaries in non-overlapping 5 kbp physical windows along each of the six segregating inversion loci: inv3.0, inv7.2, inv14.0, inv15.0, inv21.0, inv22.0. Because the Maple Ridge sample did not include any inversion homozygotes for inv14.0, we excluded within-and between-karyotype summary statistic comparisons involving this population at this locus. To examine spatial patterns of diversity and divergence along each locus, we additionally filtered variant calls to exclude sites exhibiting excess heterozygosity in order to mitigate the effects of repetitive DNA. The GATK commands used to conduct this filtering are described in the next section.

We used the means of the pairwise divergence measures described above to construct distance matrices relating inverted and standard karyotypes sampled from all three Gulf Islands populations at each polymorphic inversion locus (Supplementary Table S5). We then used the algorithm of Saitou and Nei (1987), implemented in the ape v5.7-1 R package (Paradis and Schliep 2019), to construct neighbor-joining trees for each locus.

### Obtaining a high-confidence, putatively neutral SNP call set

To obtain a high-confidence SNP call set for demographic analyses, we leveraged the technical replicates of re-sequenced mice to adjust SNP filtering thresholds in a manner that achieved low genotyping error rates while maintaining high SNP retention. In addition to the *site*-level filtering described previously, we focused our more rigorous quality filtering on two *individual*-level annotations: coverage depth (DP), which measures individual read count at a given SNP, and genotype quality score (GQ), which measures the difference between the second lowest and the lowest phred-scaled genotype likelihood. Whereas a low coverage depth can indicate poor support for an individual’s genotype call, artificially inflated coverage could reflect copy number variation that is poorly resolved in the reference assembly (Li 2014; Pfeifer 2017). By measuring pairwise genotype concordance between technical replicates, we used the rate of genotype mismatch within each pair of duplicate samples as a proxy for genotyping/sequencing error. To achieve an optimum balance of sensitivity and specificity, we tested a range of maximum DP (41X, 46.5X, 52X, and 57.5X), minimum DP (2.5X, 8X, 13.5X, and 19X), and minimum GQ (0, 25, and 50) thresholds. The impact of these filtering thresholds on SNP retention and error rates is given in Supplementary Table S6. Based on these findings, we imposed the following thresholds using GATK v4.2.0.0 VariantFiltration: “DP” > 8, “DP” < 52, and “GQ” > 25. These individual-level filters left a total of 46,287,217 SNPs in Saturna, 50,060,722 SNPs in Pender, and 57,424,563 SNPs in Maple Ridge while reducing the error rates in each sample (measured as one-half the rate of genotype mismatch within each pair of duplicate samples) to 0.06%, 0.05%, and 0.04%, respectively.

We also restricted downstream analyses to variants falling outside of annotated repetitive elements, which can cause errors in read mapping and variant calling (Pfeifer 2017). We used the union of the RepeatMasker (hub_2100979_repeatMasker; https://repeatmasker.org), WindowMasker (hub_2100979_windowMasker; Morgulis et al. 2006), TandemRepeatFinder (hub_2100979_simpleRepeat; Benson 1999), and Tandem Duplicates (hub_2100979_tandemDups) annotation tracks of the UCSC Genome Browser (http://genome.ucsc.edu; Karolchik et al 2004) to exclude SNPs within these regions. Repetitive DNA that is poorly resolved in the reference genome can also manifest as regions of high heterozygosity when reads derived from diverged repeats map to the same position. To mitigate this issue, we used GATK v4.2.0.0 VariantFiltration to remove sites with “ExcessHet” > 15.0, corresponding to a p-value less than 0.05 for the Hardy-Weinberg exact test of excess heterozygosity (Wigginton et al. 2005).

To reduce the impact of direct and linked selection, we restricted our analysis to putatively neutral SNPs by masking annotated genes in the *P. maniculatus* reference assembly. Using the NCBI RefSeq track (hub_2100979_ncbiRefSeq) of the UCSC Genome Browser (http://genome.ucsc.edu; Karolchik et al 2004), we excluded SNPs falling within 1 kbp of predicted and curated genes. Because inversions may experience distinct recombination and selective landscapes, SNPs falling within these intervals also violate the assumptions of SFS-based demographic inference frameworks. Using the breakpoints defined by Harringmeyer and Hoekstra (2022), we excluded sites within each of the 21 inversion polymorphisms they characterized from downstream analyses. Following these additional masking and filtering procedures, we were left with a total of 2,470,844 SNPs in Saturna, 2,315,952 SNPs in Pender, and 3,455,232 SNPs in Maple Ridge to use for demographic inference.

### Demographic inference

We employed the SFS-based inference approach of *Moments* (Jouganous et al. 2017) to reconstruct the joint demographic history of the Gulf Islands populations. Using the high-confidence, putatively neutral variant call set described above, we constructed 2D joint SFS (jSFS) for each pair of populations and a 3D jSFS for the combined set of populations. jSFS were folded to avoid errors in ancestral state identification and all sites containing missing genotype information for one or more individuals were excluded from analysis. To estimate demographic parameters from these jSFS, we used the maximum likelihood framework implemented in *Moments* v1.1.15.

We began by fitting simplified two-population demographic histories to the 2D jSFS for each pair of populations (Supplementary Figures S22-S24). Each model featured the divergence of contemporary populations from a shared ancestor, followed by either constant population sizes, discrete changes in size, continuous changes in size, or a combination of discrete and continuous changes in size (Supplementary Figure S22). On top of these basic population size histories, we fit additional parameters that allowed for either symmetric (Supplementary Figure S23) or asymmetric (Supplementary Figure S24) migration between populations following their split.

To fit each of the tested two-population models, we used the BFGS optimization routine in *Moments* v1.1.15 to find the parameter values that maximized the likelihood of each observed 2D jSFS. To ensure thorough exploration of the parameter space, starting parameter values were permuted across 10,000 independent searches. For each tested model, we retained the top three combinations of parameter values that yielded the highest likelihood. After fitting each model, we inspected parameter convergence, model likelihoods, and residual fit to select between contrasting models. First, we eliminated models where the coefficient of variation for one or more parameters exceeded 0.2 across the top three highest likelihood estimates. Next, we compared model likelihoods either directly (for models with equivalent numbers of parameters), or through adjusted likelihood ratio tests (for models with nested parameters; Coffman et al. 2016). For remaining models, we inspected their fit to both the joint and marginal SFS using Anscombe residuals and selected the simplest model that yielded the best fit to all partitions of the jSFS.

Our two-population model fitting and selection procedure indicated that the island-island history was best captured by a model involving recent population divergence (Supplementary Figure S25). In contrast, both island-mainland comparisons yielded best-fit models that involved ancient population divergence followed by recent discrete population size reductions (Supplementary Figures S26 and S27). All three comparisons supported a history of symmetric gene flow between populations (Supplementary Figures S25-S27).

To reconcile the extreme differences in divergence times suggested by the island-island and island-mainland comparisons, we used these candidate two-population histories to construct a model that reflects the shared history of all three Gulf Islands populations. This three-population model featured an older epoch in which the ancestral island and mainland lineages diverge from each other, followed by a more recent epoch reflecting the establishment of the contemporary island and mainland populations (Supplementary Figure S28). Given the evidence for gene flow, we fit all possible parameterizations of symmetric migration among ancestral and contemporary populations (Supplementary Figure S28) to the 3D jSFS. We used the same model fitting and model selection approach described above to obtain demographic parameter estimates and select among contrasting models. Residual differences between the joint and marginal SFS observed for each population/population-pair and the expected SFS under the best-fitting three-population demographic model are illustrated in Supplementary Figure S29.

For both two- and three-population candidate models, the ancestral effective population sizes were calculated as N_Anc_ = θ/(4*µ*L) using an assumed mutation rate, µ, of 5e-9 per-site, per-generation and an effective sequence length, L, that represents the total length of “callable” sequence. To approximate L, we multiplied the total length of the *P. maniculatus* autosomal genome (2,302,074,966 bp) by the average percentage of SNPs (2.34%) that were retained following both the site- and individual-level filtering as well as the masks we applied to repetitive, genic, and inverted DNA. This yielded an approximate L of 53,870,000 bp. Using the formula above, estimates of population size and time parameters were scaled by N_Anc_ and 2*N_Anc_ to convert them into units of individuals and generations, respectively. Parameter uncertainties were estimated using the Godambe Information Matrix methods described in Coffman et al. (2016). We used uncertainty propagation techniques (Ku 1966) to obtain the standard deviations for parameters that are functions of multiple estimated quantities.

We conducted predictive simulations with *msprime* v1.2.0 (Baumdicker et al. 2022) to evaluate the fit of the inferred three-population demographic history. Using the maximum likelihood parameter estimates obtained for the best-fit three-population model, we conducted 2,500 independent simulations of a 1 Mbp genomic element for each component population assuming uniform recombination and mutation rates of 5e-9 per-site, per-generation. Simulated sample sizes were set to match those of the empirical data for each population. We computed nucleotide diversity 𝜋 (Tajima 1983), Tajima’s *D* (Tajima 1989), and F_ST_ (Hudson et al. 1992) in non-overlapping 5 kbp physical windows using scikit-allel v1.3.6 (https://github.com/cggh/scikit-allel) for both the simulated and empirical data. Together, the quality-based filtering we conducted on our empirical SNP call set and the masks we applied to repetitive, genic, and inverted DNA introduced variability in the number of sites “observed” among the 5 kbp physical windows that we used to compute each summary statistic. To account for this variability, we used the “is_accessible” argument in scikit-allel to compute per-site statistics based on the total number of observable sites. To measure the decay of LD in both simulated and empirical data, we used the “--r2” option in plink v1.90 (Chang et al. 2015) to compute pairwise r^2^. We restricted this analysis to SNPs separated by at most 1 Mbp by setting the “--ld-window” and “-- ld-window-kb” arguments to 1,000,000 and 1,000, respectively. Singleton variants were excluded from these calculations by setting the “--mac” argument to 1.

Using the same jSFS employed for demographic inference, we assessed whether the joint minor allele frequencies of the six segregating inversion loci were unusual compared to neutral, non-inverted loci. We summarized joint frequencies by computing the Euclidean distance between each population’s minor allele frequencies at the six inversion loci. To obtain the null distributions of this distance measure, we constructed 1,000 random samples of six SNPs from the jSFS and computed the Euclidean distance between minor allele frequencies for each population pair using the dist() function in R v4.2.1 (R Core Team 2022).

### Contemporary effective population size estimation

We used the LD-based approach of *currentNe* (Santiago et al. 2024) to estimate contemporary effective population sizes among Gulf Islands populations. Excluding close relatives in a random sample of individuals can artificially inflate estimates of contemporary effective population size obtained through LD-based methods (Waples and Anderson 2017; Santiago et al. 2024). Since the excess of close relatives we found in each sample could be driven by small contemporary effective population sizes or nonrandom trapping, we chose to conduct these analyses on both the full sample (excluding technical replicates) and on the subset of unrelated individuals from each population (obtained by excluding relationships ≤ 2^nd^ degree). The same SNP call set employed for demographic inference was used as input to these analyses. We used the “-s” argument to restrict each analysis to 50,000 randomly sampled SNPs and restricted LD calculations to pairs of SNPs located on different chromosomes. When conducting analyses on samples that included close relatives, we used the “-k” option to jointly estimate contemporary effective population size and k, which represents the average number of full siblings each individual has in the sample (Santiago et al. 2024). We use the artificial neutral network approach implemented in *currentNe* to obtain 90% confidence intervals for each point estimate of contemporary effective population size (Santiago et al. 2024).

## ACKNOWLEDGEMENTS

We thank members of the Payseur lab for their helpful input on this work. This research was funded by National Institutes of Health (NIH) grants R01GM100426 and R35GM139412 (to B.A.P.). E.K.H. was partially supported by the NIH Graduate Training Grant in Genetics at the University of Wisconsin-Madison (T32GM007133). Computational analyses were conducted using resources provided by the University of Wisconsin-Madison’s Center for High Throughput Computing. Fieldwork was supported by a Putnam Expedition Grant of the Harvard Museum of Comparative Zoology. F.B. was supported by an HHMI International Student Research Fellowship, a Joan Brockman Williamson Graduate Research Fellowship, and a Herchel Smith Graduate Fellowship. This work was supported by the Howard Hughes Medical Institute, of which H.E.H. was an Investigator.

## DATA ACCESSIBILITY

Upon submission, raw sequencing reads will be deposited in the SRA. Code used to conduct the analyses is available on GitHub (https://github.com/PayseurLabUWMadison/gi_demography_inversions).

## AUTHOR CONTRIBUTIONS

E.K.H. and B.A.P. designed the study. F.B. and H.E.H acquired the samples used for the study. E.K.H. conducted the research with supervision from B.A.P. E.K.H. and B.A.P. wrote the manuscript with input from F.B. and H.E.H.

