## Supplementary Figure for "The genomic imprint of chromosomal inversions and demographic history in island populations of deer mice"

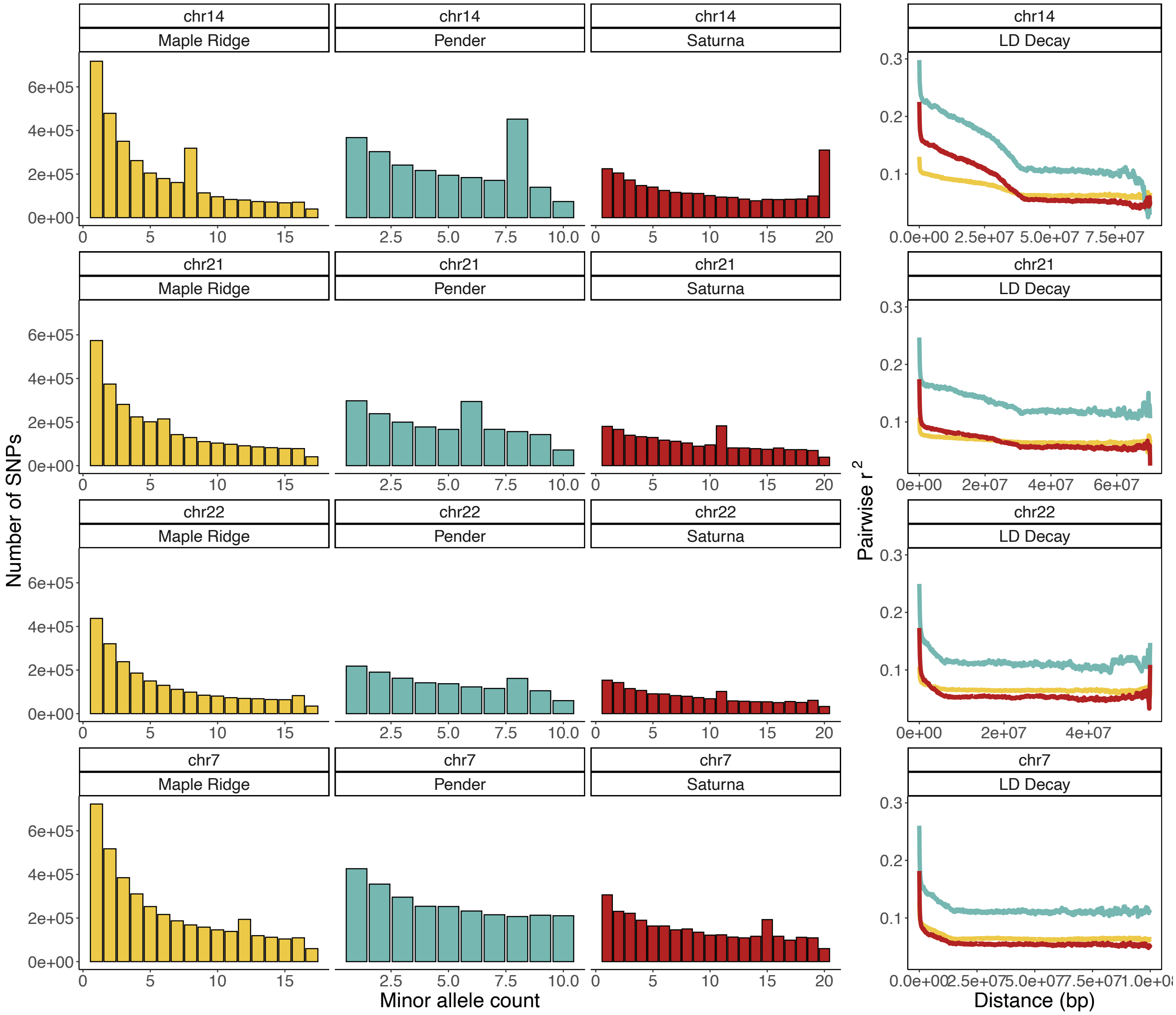


**Figure S1. Inversion polymorphisms drive irregularities in the SFS and cause distortions in LD decay.** Each row illustrates how segregating inversions affect chromosome-wide allele frequencies and linkage disequilibrium on chromosomes 14, 21, 22, and 7 (in order from top to bottom). Histograms in each row plot the folded, chromosome-wide SFS for Maple Ridge (yellow), Pender (blue), and Saturna (red). Curves in each row plot average pairwise $r^2$ (y-axis) between SNPs separated by increasing distance (x-axis) in each population. LD decay curves for each population are colored according to their SFS.

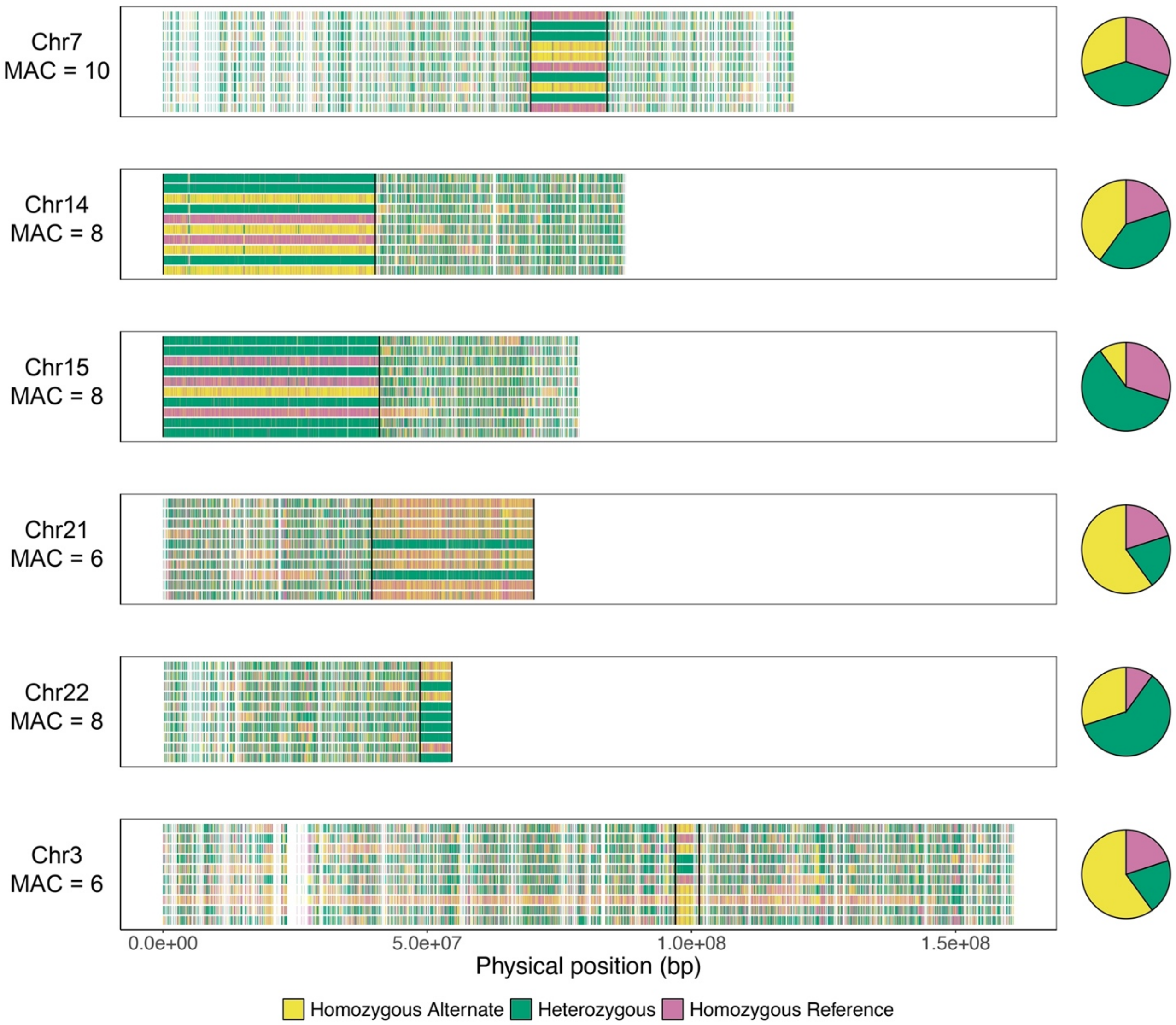


**Figures S2. SNP genotypes partitioned by minor allele frequency in the Pender Island sample.** Plots illustrate the positive results for the inversion screen conducted in the Pender Island sample. Each horizontal panel represents a different autosome: chr7, chr14, chr15, chr21, chr22, and chr3 (in order from top to bottom). Labels beneath each autosome name indicate the corresponding minor allele count of the inversion in the sample. X-axes measure the physical position along each autosome. Within a panel, each row represents a sampled individual, and each vertical tick mark indicates the location of a SNP with the specified minor allele count. SNPs are colored according to their diploid genotype in each individual (yellow for homozygous alternate, green for heterozygous, and pink for homozygous reference). Vertical black lines denote the breakpoints defined by Harringmeyer and Hoekstra (2022) for inv7.2 (a), inv14.0 (b), inv15.0 (c), inv21.0 (d), inv22.0 (e), and inv3.0 (f). Pie charts adjacent to each panel denote the genotype proportions of the arrangement in each sample.

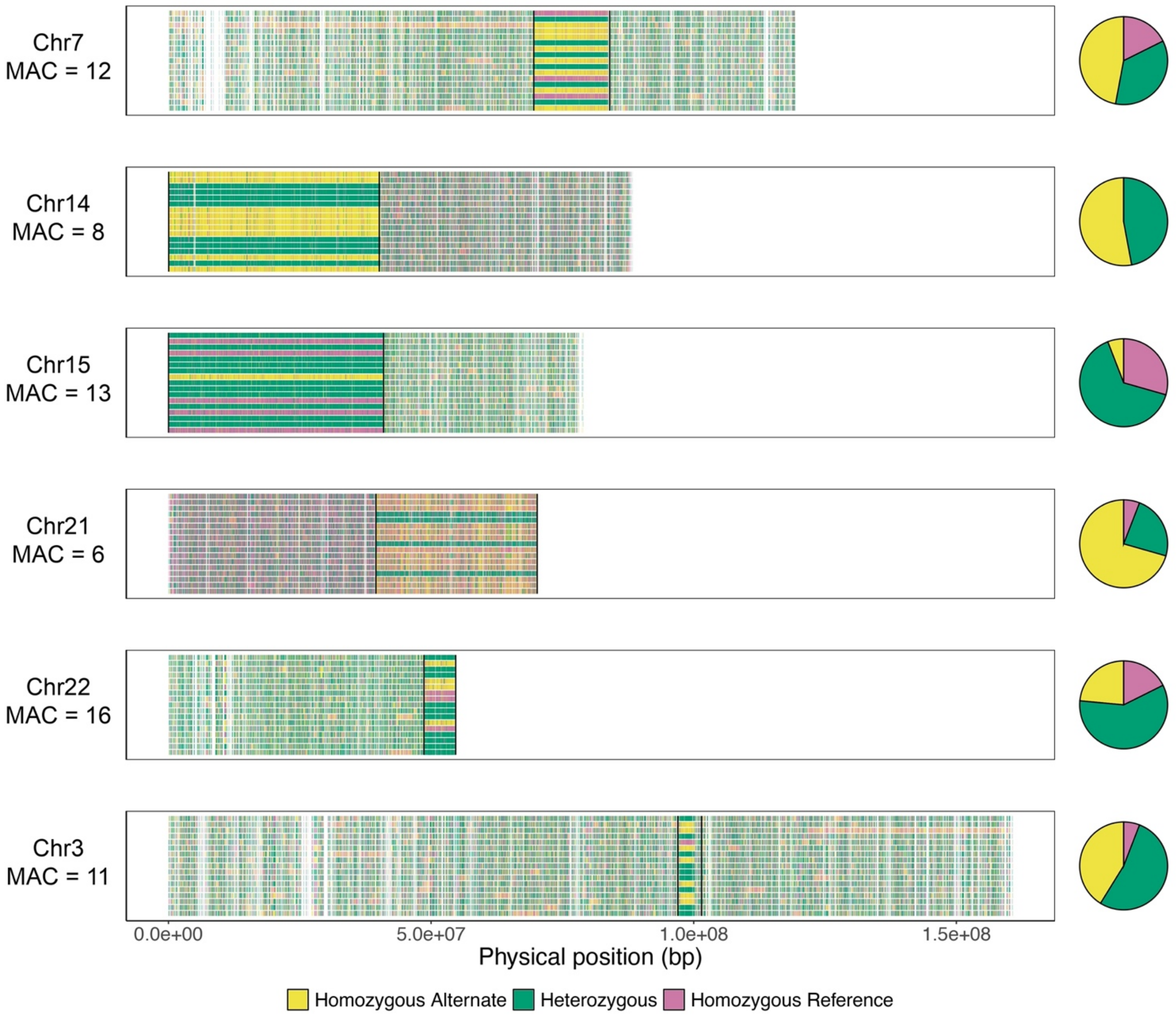


**Figures S3. SNP genotypes partitioned by minor allele frequency in the mainland Maple Ridge sample.** Plots illustrate the positive results for the inversion screen conducted in the mainland Maple Ridge sample. Each horizontal panel represents a different autosome: chr7, chr14, chr15, chr21, chr22, and chr3 (in order from top to bottom). Labels beneath each autosome name indicate the corresponding minor allele count of the inversion in the sample. X-axes measure the physical position along each autosome. Within a panel, each row represents a sampled individual, and each vertical tick mark indicates the location of a SNP with the specified minor allele count. SNPs are colored according to their diploid genotype in each individual (yellow for homozygous alternate, green for heterozygous, and pink for homozygous reference). Vertical black lines denote the breakpoints defined by Harringmeyer and Hoekstra (2022) for inv7.2 (a), inv14.0 (b), inv15.0 (c), inv21.0 (d), inv22.0 (e), and inv3.0 (f). Pie charts adjacent to each panel denote the genotype proportions of the arrangement in each sample.

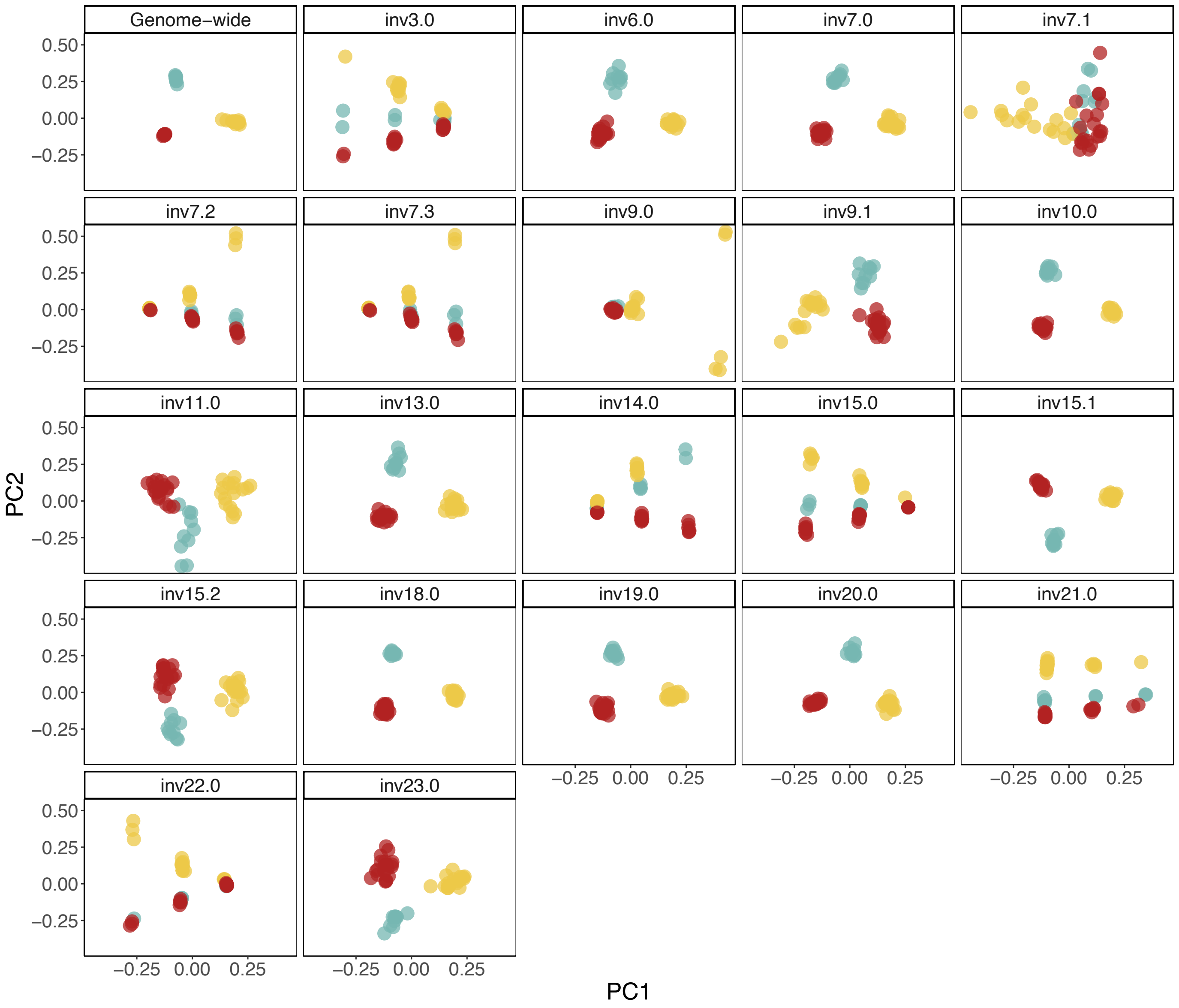


**Figure S4. Individuals from distinct populations cluster according to inversion genotype at six polymorphic loci.** Panels compare the results of PCA conducted on genome-wide SNPs falling outside of known inversions to PCA performed on SNPs within each polymorphic inversion locus identified by Harringmeyer and Hoekstra (2022) in their broad geographic survey of mainland *P. maniculatus*. Each point denotes an individual's position along the first (x-axis) and second (y-axis) principal component. Points are colored according to population sample (yellow for Maple Ridge, blue for Pender Island, and red for Saturna Island). Whereas most of the loci surveyed cluster individuals according to population (as observed for genome-wide, non-inverted SNPs), individuals instead cluster according to inversion genotype at the six segregating loci we identified: inv3.0, inv7.2, inv14.0, inv15.0, inv21.0, and inv22.0, with the exception of inv9.0. At the inv9.0 locus, Maple Ridge individuals form two clusters along PC1, raising the possibility that two genotypic classes are present. However, we find no further evidence for this polymorphism based on allele frequencies or patterns of linkage disequilibrium in Maple Ridge. We also note that, though inv7.3 produces a clustering pattern consistent with a polymorphic inversion, it partitions individuals in an identical manner to inv7.2 (in which it is nested), indicating that the inv7.3 locus itself is not segregating.

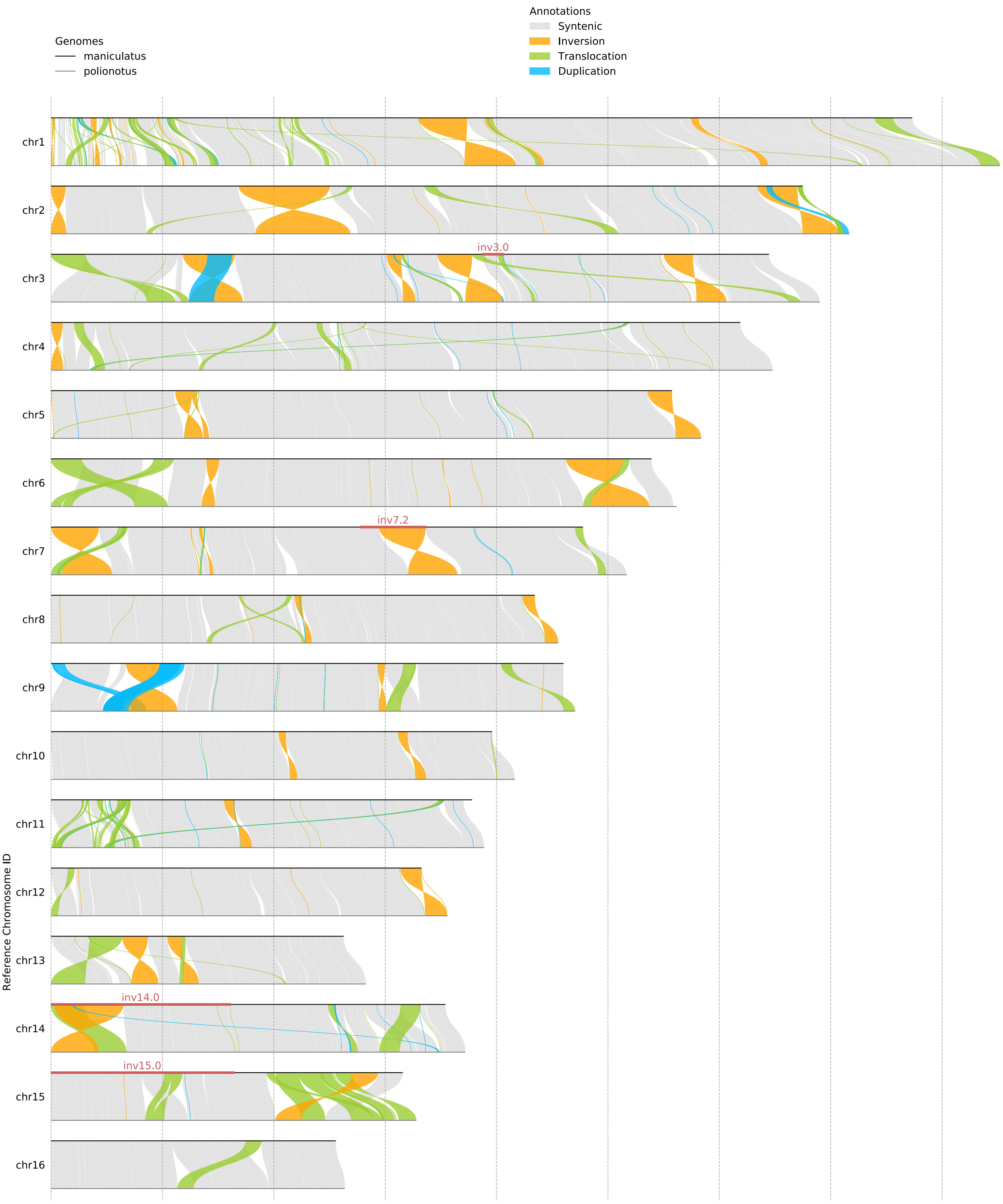
Genomes
maniculatus
polionotus
Annotations
Syntenic
Inversion
Translocation
Duplication
Reference Chromosome ID
chr1
chr2
chr3
chr4
chr5
chr6
chr7
chr8
chr9
chr10
chr11
chr12
chr13
chr14
chr15
chr16
inv3.0
inv7.2
inv14.0
inv15.0

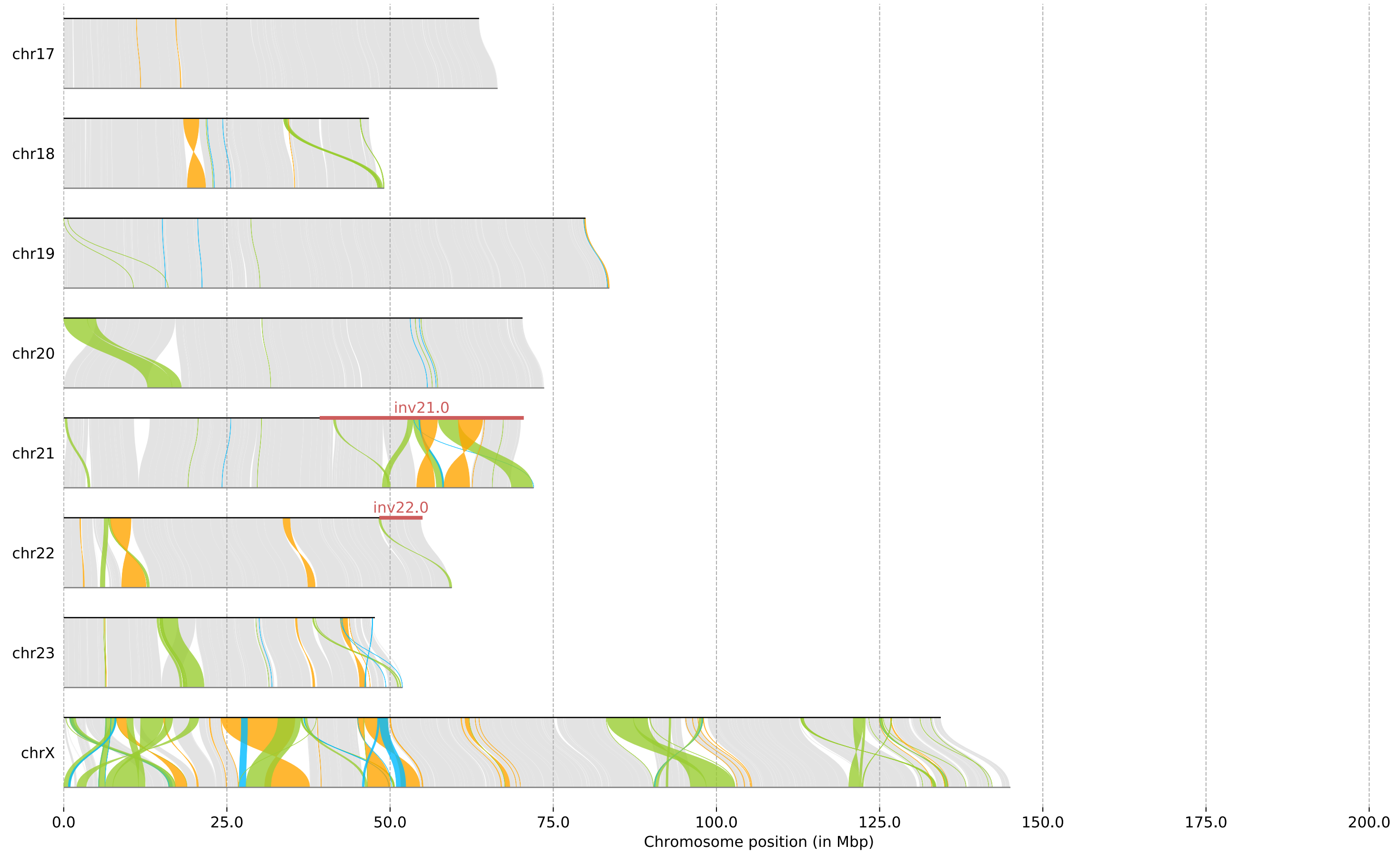


**Figure S5. Large-scale structural rearrangements between the *P. maniculatus* and *P. polionotus* reference assemblies.** Each horizontal facet represents a different chromosome. Chromosome IDs on the y-axis reflect chromosome assignments in the *P. maniculatus* reference. *P. maniculatus* chromosomes are represented at the top of each synteny plot by a black line and homologous *P. polionotus* chromosomes are represented at the bottom of each plot by a gray line. Gray ribbons between the two chromosomes represent aligned regions of synteny between the two species. Structural rearrangements are represented by orange ribbons (inversions), green ribbons (translocations), and blue ribbons (duplications). The genomic locations of the six inversions segregating within the Gulf Islands populations are represented by red bars along the *P. maniculatus* chromosome.

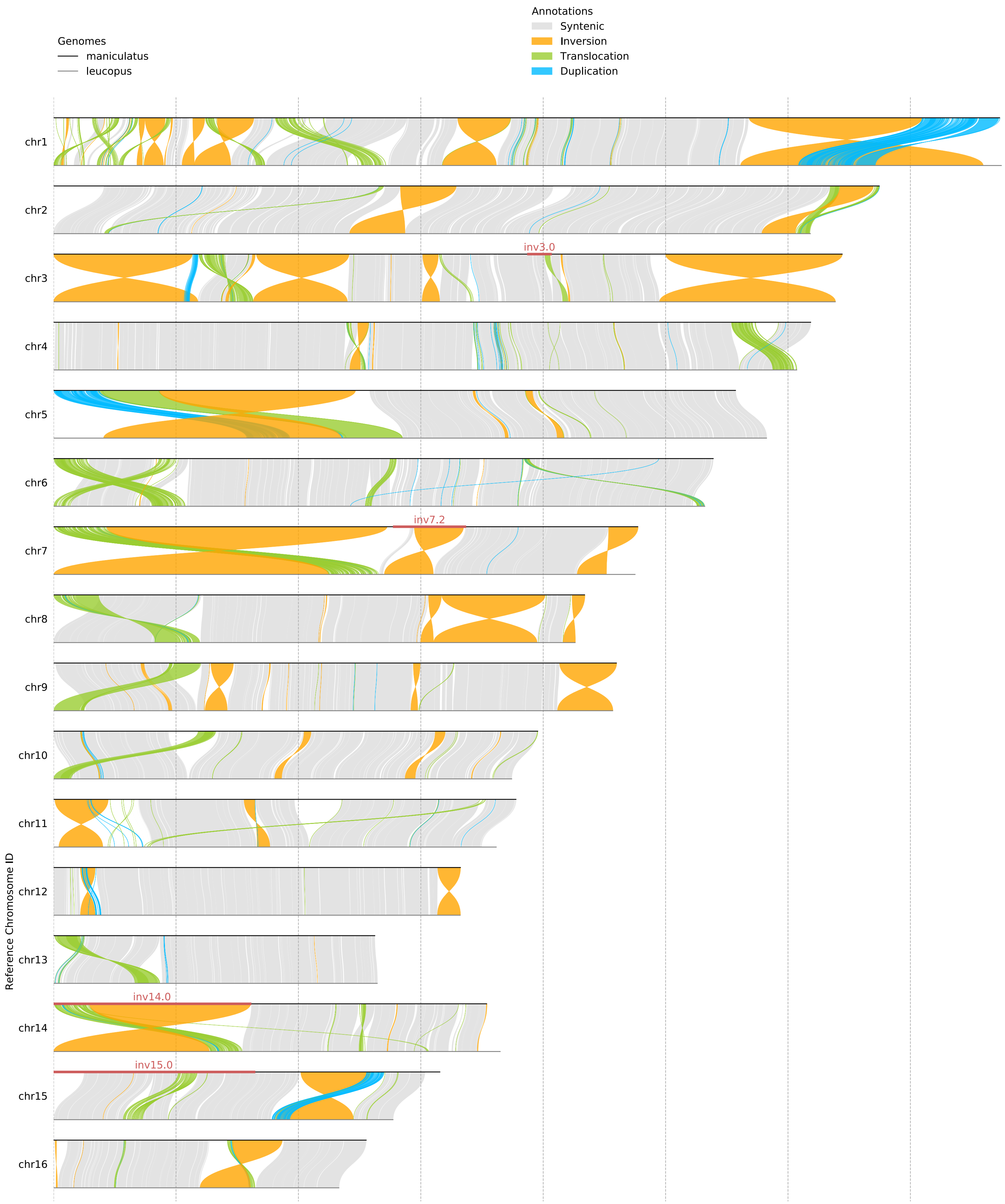

Annotations
Syntenic
Inversion
Translocation
Duplication
Genomes
maniculatus
leucopus
chr1
chr2
chr3
inv3.0
chr4
chr5
chr6
chr7
inv7.2
chr8
chr9
chr10
chr11
chr12
chr13
chr14
inv14.0
chr15
inv15.0
chr16
Reference Chromosome ID

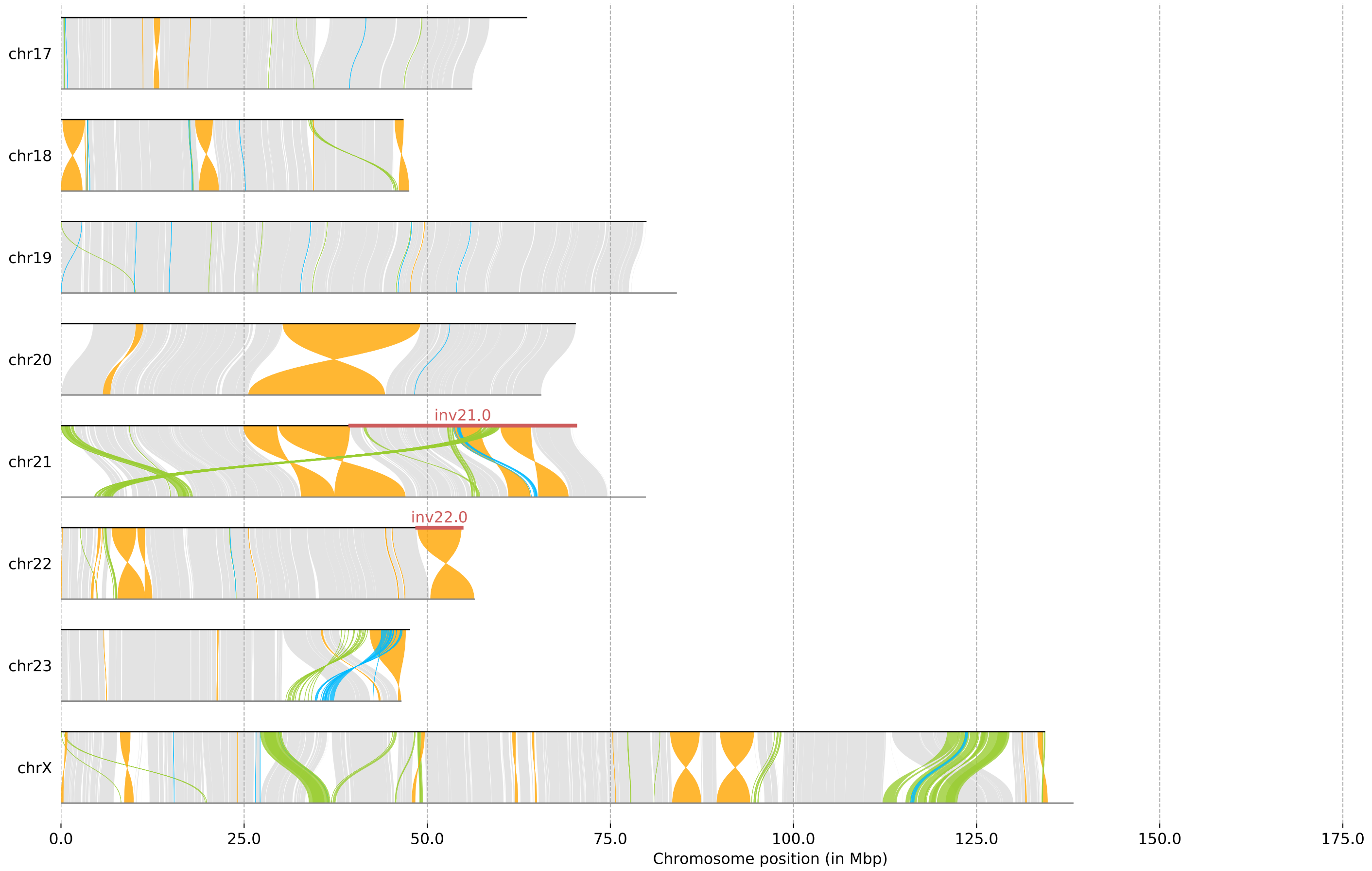


**Figure S6. Large-scale structural rearrangements between the *P. maniculatus* and *P. leucopus* reference assemblies.** Each horizontal facet represents a different chromosome. Chromosome IDs on the y-axis reflect chromosome assignments in the *P. maniculatus* reference. *P. maniculatus* chromosomes are represented at the top of each synteny plot by a black line and homologous *P. leucopus* chromosomes are represented at the bottom of each plot by a gray line. Gray ribbons between the two chromosomes represent aligned regions of synteny between the two species. Structural rearrangements are represented by orange ribbons (inversions), green ribbons (translocations), and blue ribbons (duplications). The genomic locations of the six inversions segregating within the Gulf Islands populations are represented by red bars along the *P. maniculatus* chromosome.

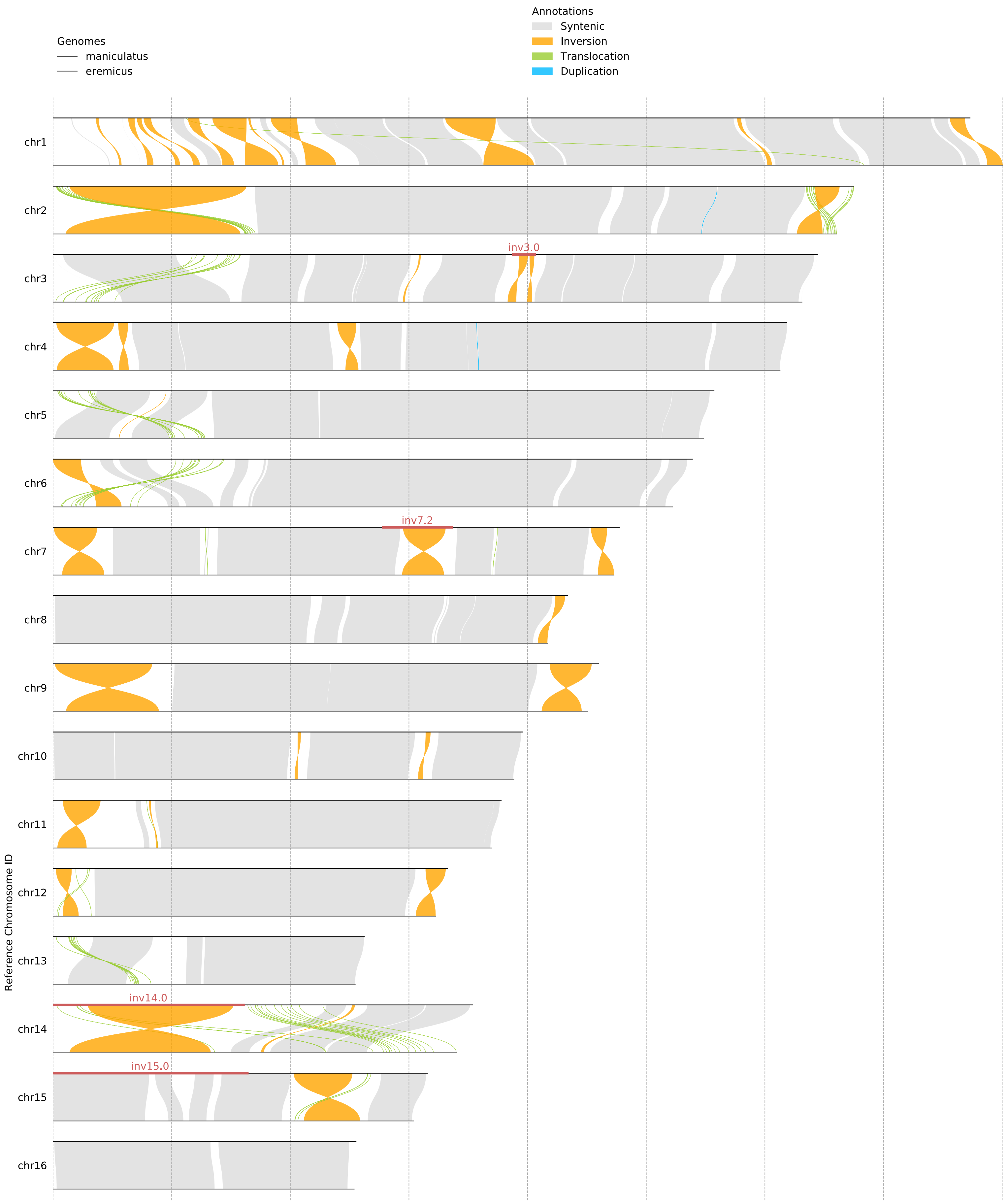
Genomes
maniculatus
eremicus
Annotations
Syntenic
Inversion
Translocation
Duplication
chr1
chr2
chr3
inv3.0
chr4
chr5
chr6
chr7
inv7.2
chr8
chr9
chr10
chr11
chr12
chr13
chr14
inv14.0
chr15
inv15.0
chr16
Reference Chromosome ID

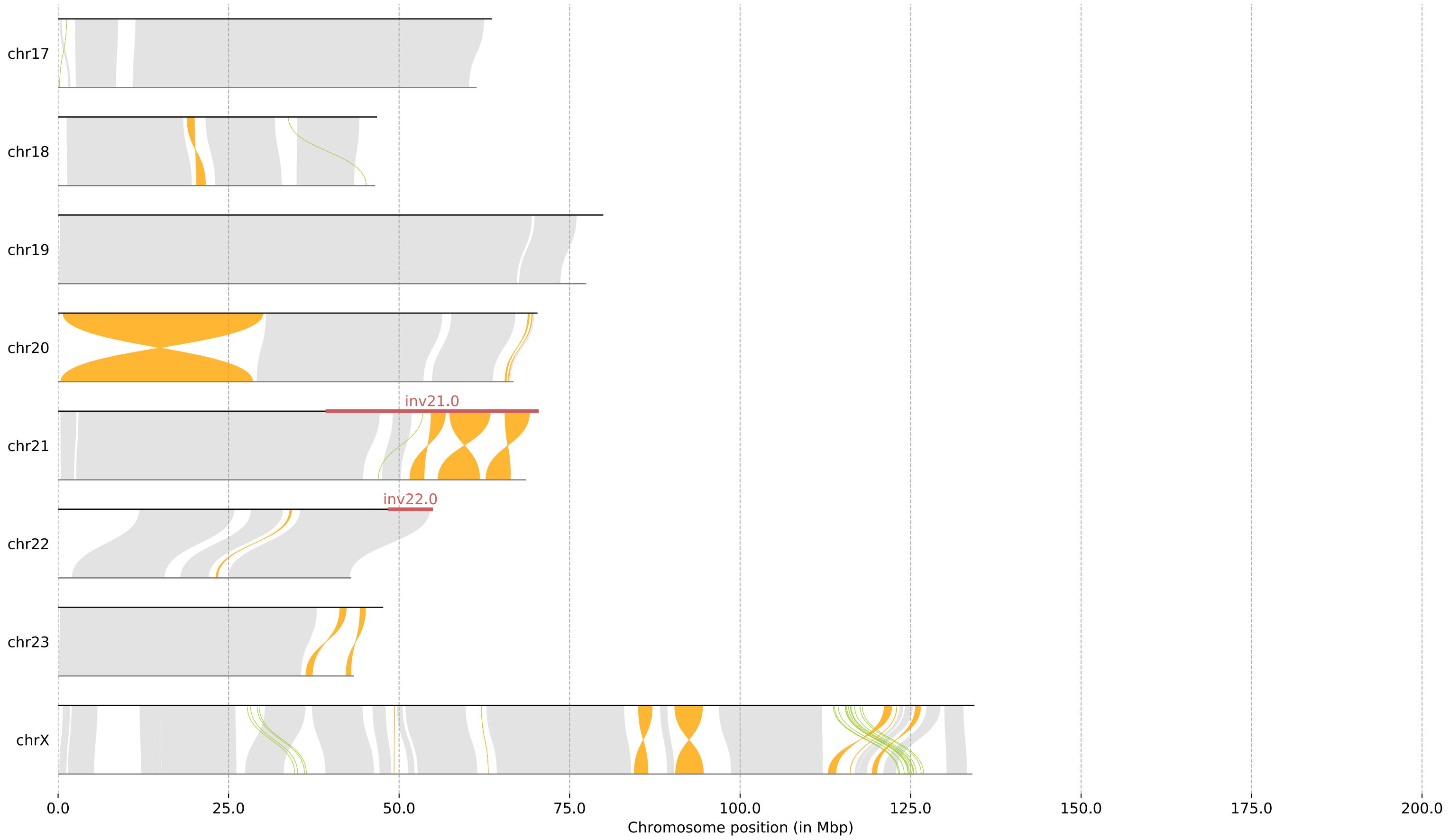


**Figure S7. Large-scale structural rearrangements between the *P. maniculatus* and *P. eremicus* reference assemblies.** Each horizontal facet represents a different chromosome. Chromosome IDs on the y-axis reflect chromosome assignments in the *P. maniculatus* reference. *P. maniculatus* chromosomes are represented at the top of each synteny plot by a black line and homologous *P. eremicus* chromosomes are represented at the bottom of each plot by a gray line. Gray ribbons between the two chromosomes represent aligned regions of synteny between the two species. Structural rearrangements are represented by orange ribbons (inversions), green ribbons (translocations), and blue ribbons (duplications). The genomic locations of the six inversions segregating within the Gulf Islands populations are represented by red bars along the *P. maniculatus* chromosome.

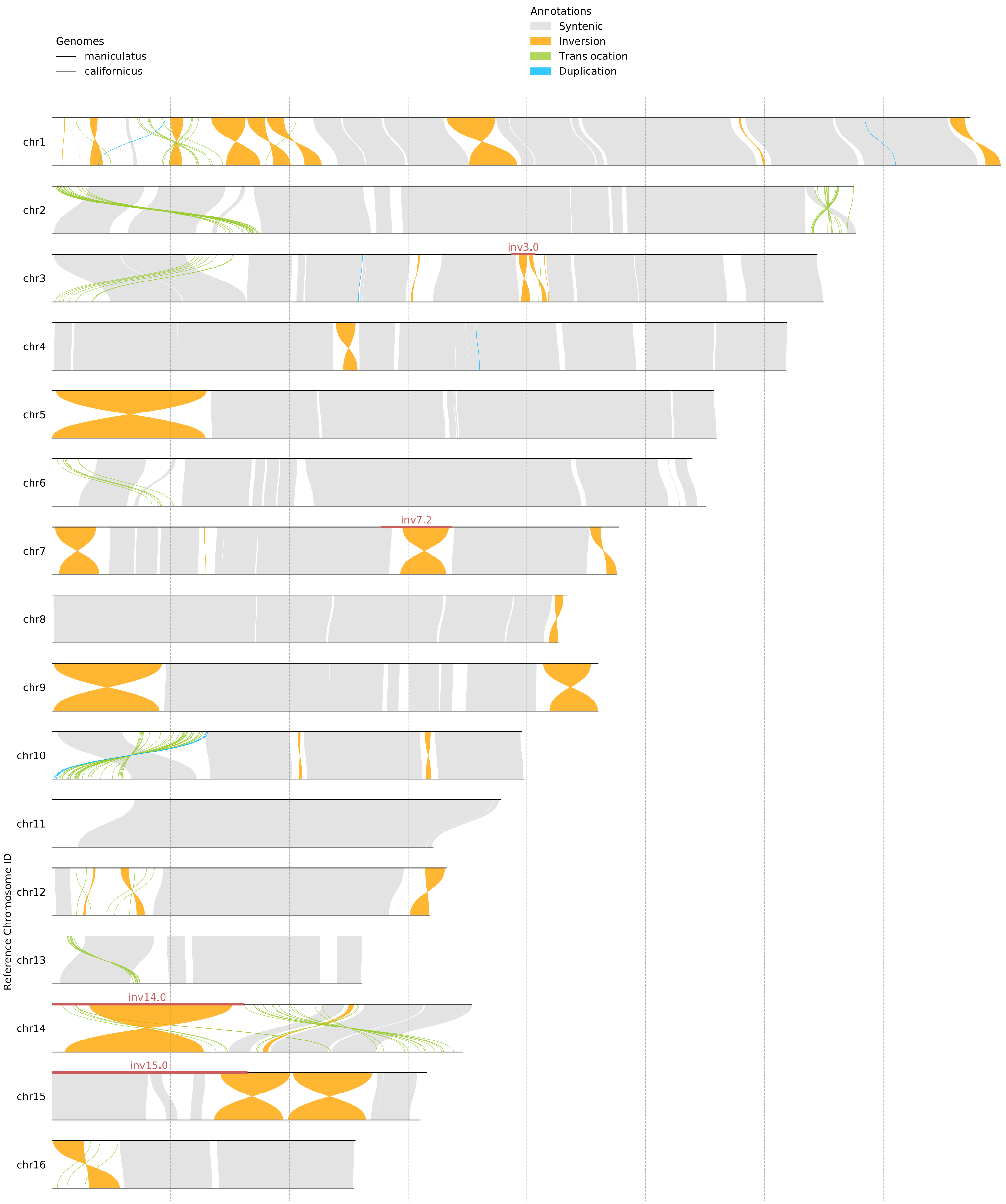

Genomes
maniculatus
californicus
Annotations
Syntenic
Inversion
Translocation
Duplication
Reference Chromosome ID
chr1
chr2
chr3
inv3.0
chr4
chr5
chr6
chr7
inv7.2
chr8
chr9
chr10
chr11
chr12
chr13
chr14
inv14.0
chr15
inv15.0
chr16

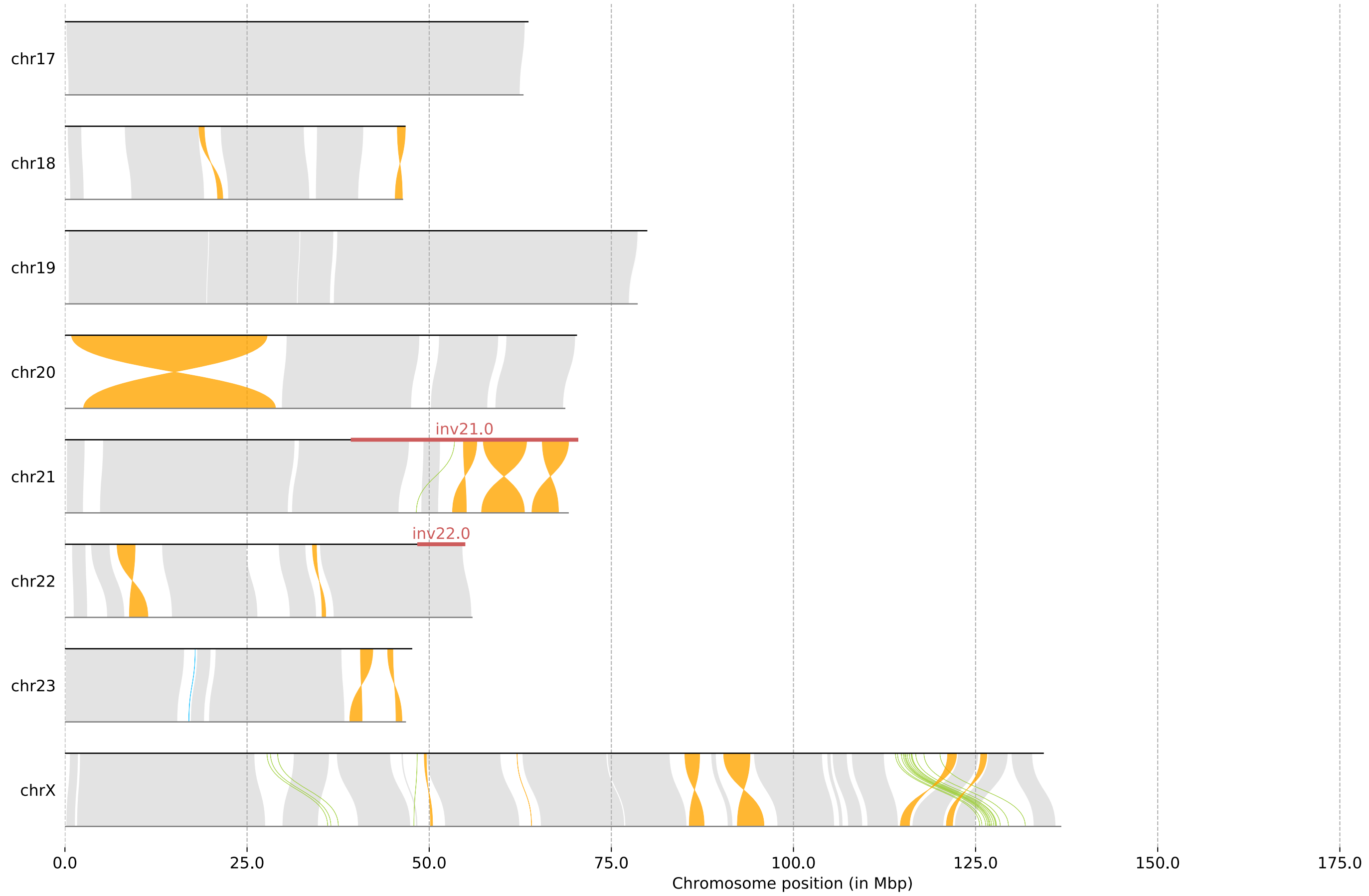


**Figure S8. Large-scale structural rearrangements between the *P. maniculatus* and *P. californicus* reference assemblies.** Each horizontal facet represents a different chromosome. Chromosome IDs on the y-axis reflect chromosome assignments in the *P. maniculatus* reference. *P. maniculatus* chromosomes are represented at the top of each synteny plot by a black line and homologous *P. californicus* chromosomes are represented at the bottom of each plot by a gray line. Gray ribbons between the two chromosomes represent aligned regions of synteny between the two species. Structural rearrangements are represented by orange ribbons (inversions), green ribbons (translocations), and blue ribbons (duplications). The genomic locations of the six inversions segregating within the Gulf Islands populations are represented by red bars along the *P. maniculatus* chromosome.

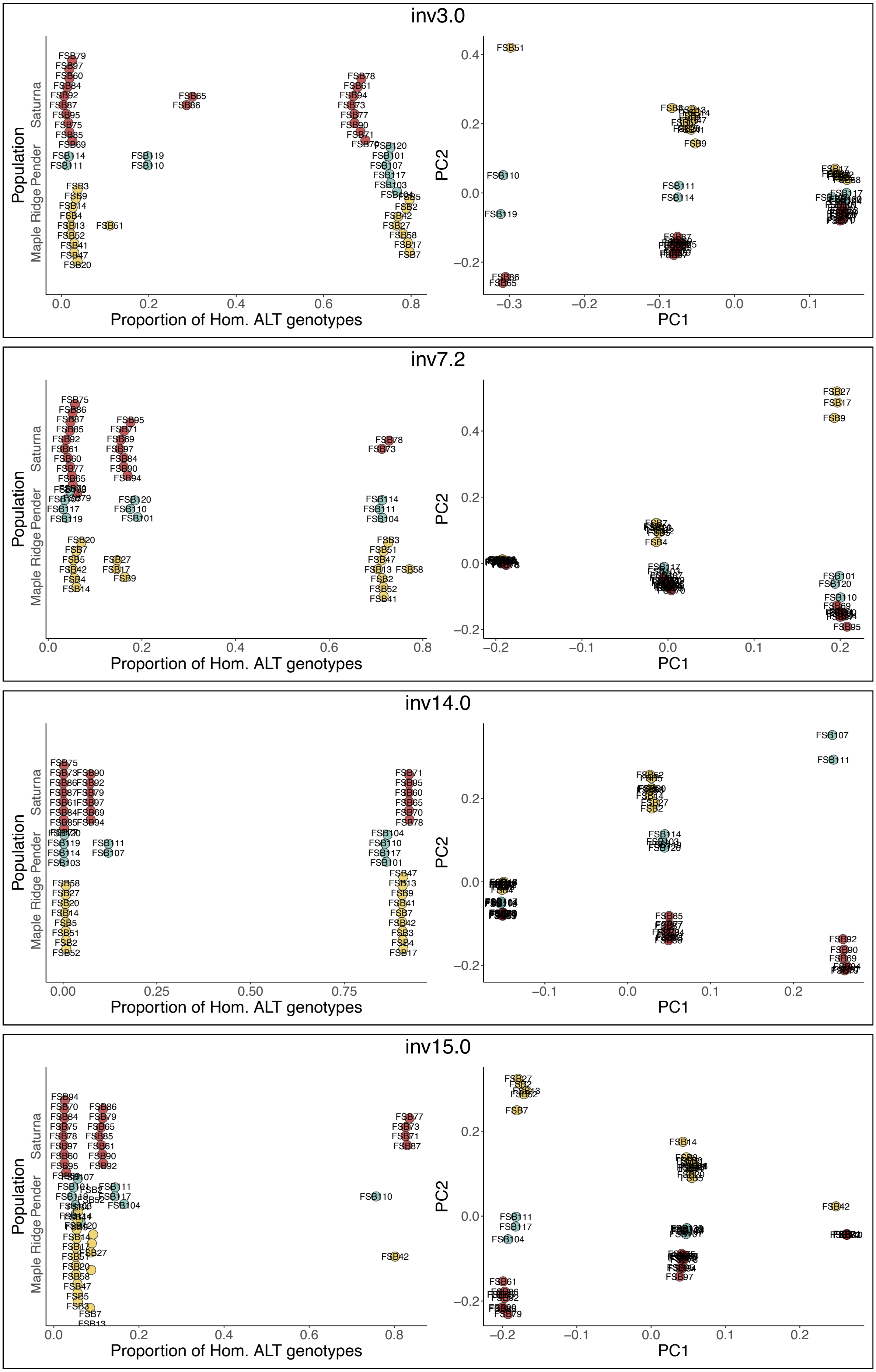

inv3.0
inv7.2
inv14.0
inv15.0
Population
Saturna
Pender
Maple Ridge
Proportion of Hom. ALT genotypes
PC1
PC2

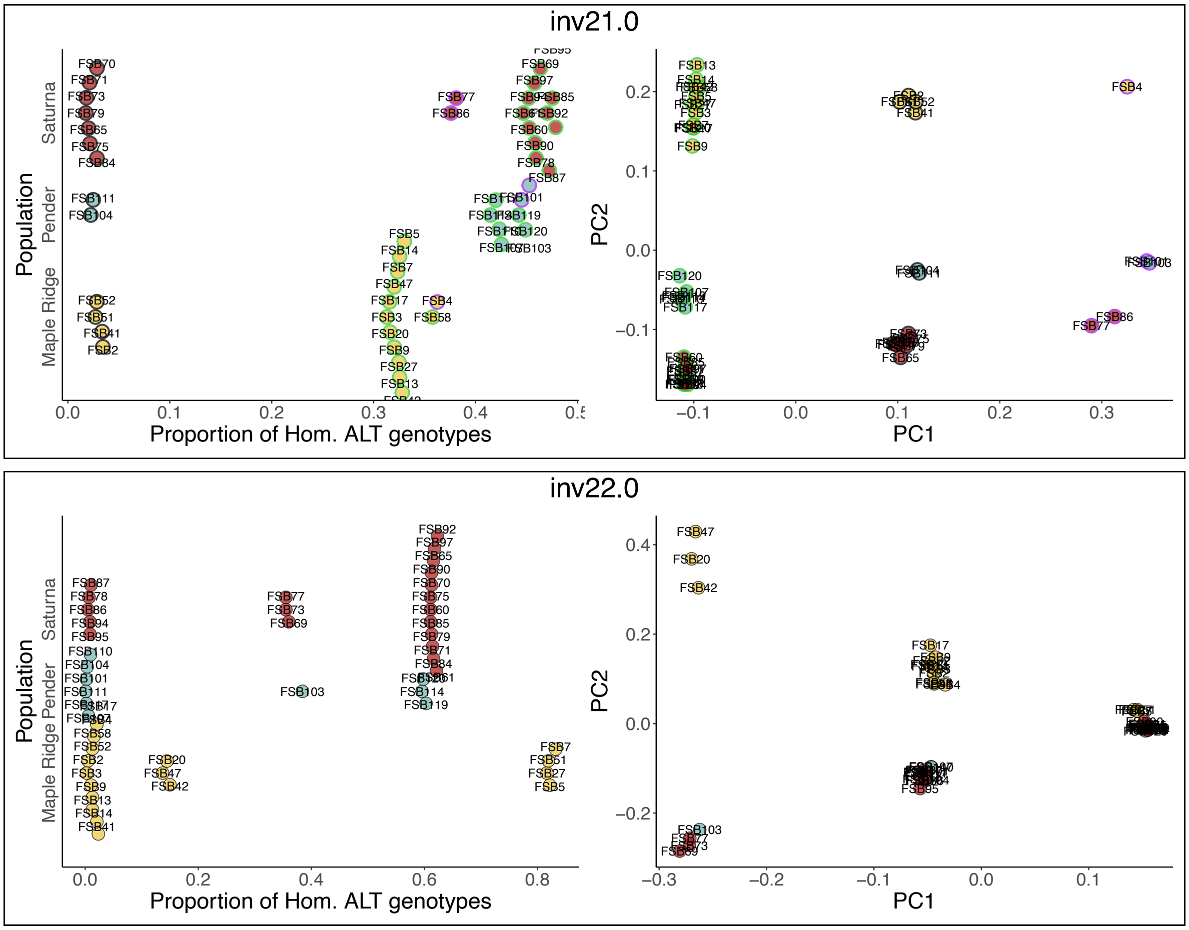


**Figure S9. Comparison between genotypic composition and PCA classifications at six polymorphic inversion loci.** Each panel reflects a different polymorphic inversion locus: inv3.0, inv7.2, inv14.0, inv15.0, inv21.0, and inv22.0 (ordered from top to bottom). Points within each pair of plots represent individual mice. Points are colored according to sampling location (red for Saturna, blue for Pender, and yellow for Maple Ridge) and point labels reflect sample IDs. Left-hand plots depict the proportion of homozygous alternate (ALT/ALT) genotypes among SNPs segregating at the minor allele frequency of the inversion polymorphism (i.e., the same SNPs illustrated in Figure 4 and Supplementary figures S2 and S3). Points are vertically jittered for clarity. Right-hand plots depict the same PCA results illustrated in Figure 5 and Supplementary figure S4 for each locus. With the exception of inv21.0, grouping based on proportion of ALT/ALT genotypes matches PCA clusters. At inv21.0, individuals comprising the right-most PCA cluster do not group consistently based on their proportion of ALT/ALT genotypes. To illustrate this, points for inv21.0 are outlined in both plots to reflect PCA cluster (left cluster: green, middle cluster: black, right cluster: purple). Whereas right-cluster individuals in Saturna (FSB77 and FSB86) have fewer ALT/ALT genotypes than left-cluster individuals, this pattern is reversed for right-cluster individuals in Maple Ridge (FSB4) and Pender (FSB101 and FSB103).

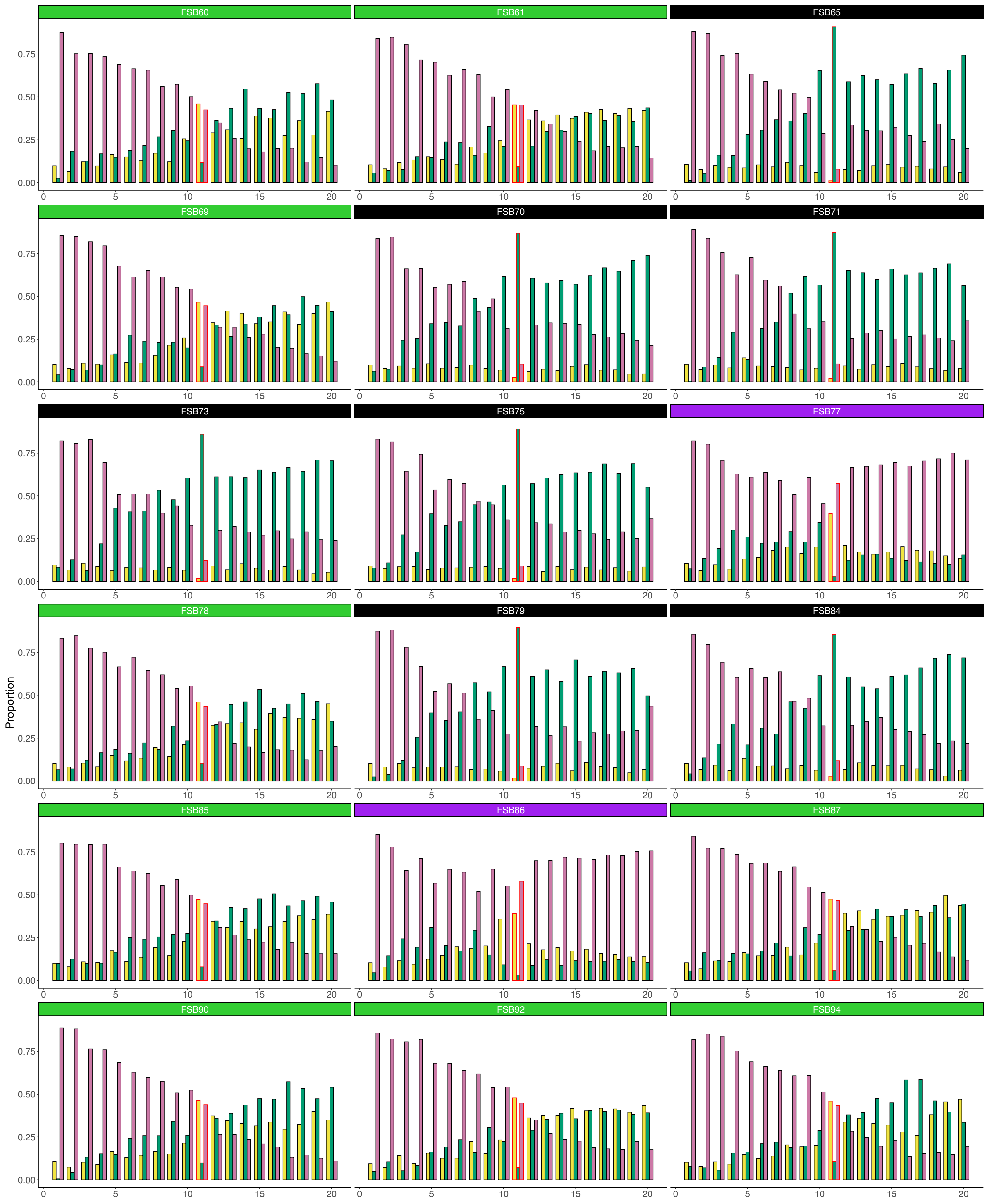
FSB60
FSB61
FSB65
FSB69
FSB70
FSB71
FSB73
FSB75
FSB77
FSB78
FSB79
FSB84
FSB85
FSB86
FSB87
FSB90
FSB92
FSB94
Proportion
0.00
0.25
0.50
0.75
0
5
10
15
20

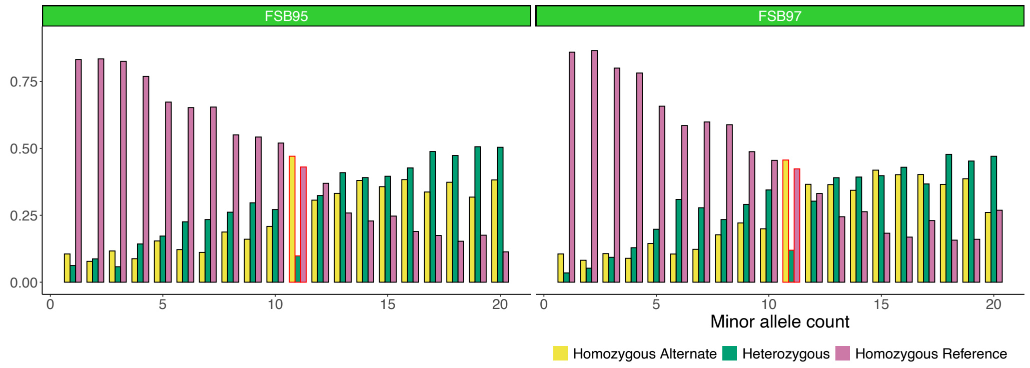


**Figure S10. Inversion homozygotes in Saturna show an enrichment of ALT alleles at minor allele frequencies approaching that of the inv21.0 polymorphism.** Each panel represents genotype proportions measured in a single Saturna individual for SNPs spanning the inv21.0 interval. Panel headers denote the individual's sample ID and are colored to match the PCA cluster assignments given in Supplementary Figure S9 for inv21.0 SNPs (left cluster homozygotes: green, middle cluster heterozygotes: black, right cluster homozygotes: purple). Bars represent the individual's proportion of each genotype (pink for REF/REF, green for HET, and yellow for ALT/ALT) among SNPs at each minor allele frequency bin (denoted by x-axes). The group of bars outlined in red in each facet denote the minor allele frequency at which inv21.0 is segregating the Saturna sample. For right cluster individuals (FSB77 and FSB86; in purple), there is an enrichment of REF alleles at minor allele frequencies exceeding that of the inv21.0 arrangement. In contrast, for left cluster individuals (FSB60, FSB61, FSB69, FSB78, FSB85, FSB87, FSB90, FSB92, FSB94, FSB95, and FSB97; in green), there is an enrichment of ALT alleles at these higher minor allele frequencies.

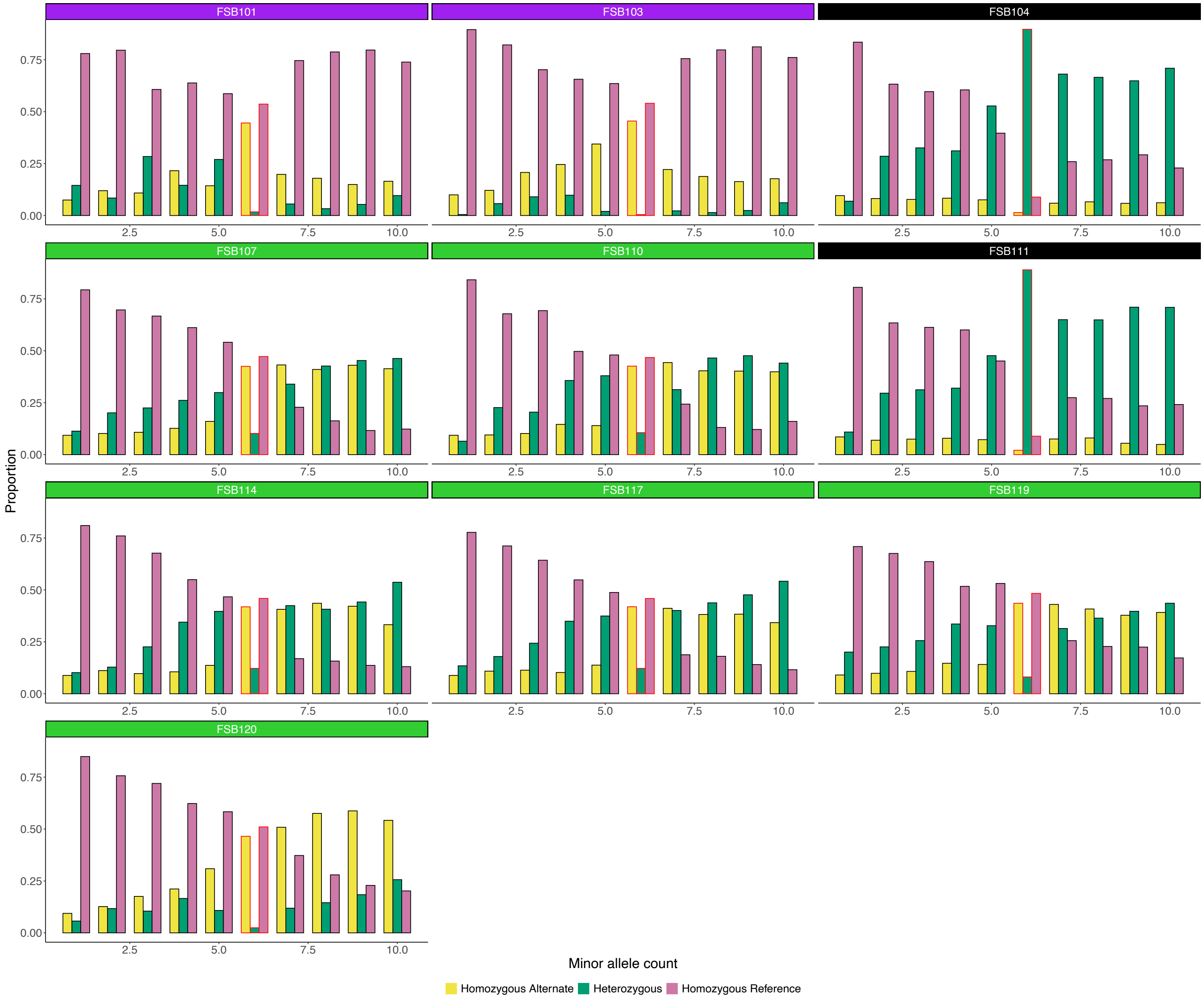


**Figure S11. Inversion homozygotes in Pender show an enrichment of ALT alleles at minor allele frequencies approaching that of the inv21.0 polymorphism.** Each panel represents genotype proportions measured in a single Pender individual for SNPs spanning the inv21.0 interval. Panel headers denote the individual's sample ID and are colored to match the PCA cluster assignments given in Supplementary Figure S9 for inv21.0 SNPs (left cluster homozygotes: green, middle cluster heterozygotes: black, right cluster homozygotes: purple). Bars represent the individual's proportion of each genotype (pink for REF/REF, green for HET, and yellow for ALT/ALT) among SNPs at each minor allele frequency bin (denoted by x-axes). The group of bars outlined in red in each facet denote the minor allele frequency at which inv21.0 is segregating the Pender sample. For right cluster individuals (FSB101 and FSB103; in purple), there is an enrichment of REF alleles at minor allele frequencies exceeding that of the inv21.0

arrangement. In contrast, for left cluster individuals (FSB107, FSB110, FSB114, FSB117, FSB119, and FSB120; in green), there is an enrichment of ALT alleles at these higher minor allele frequencies.

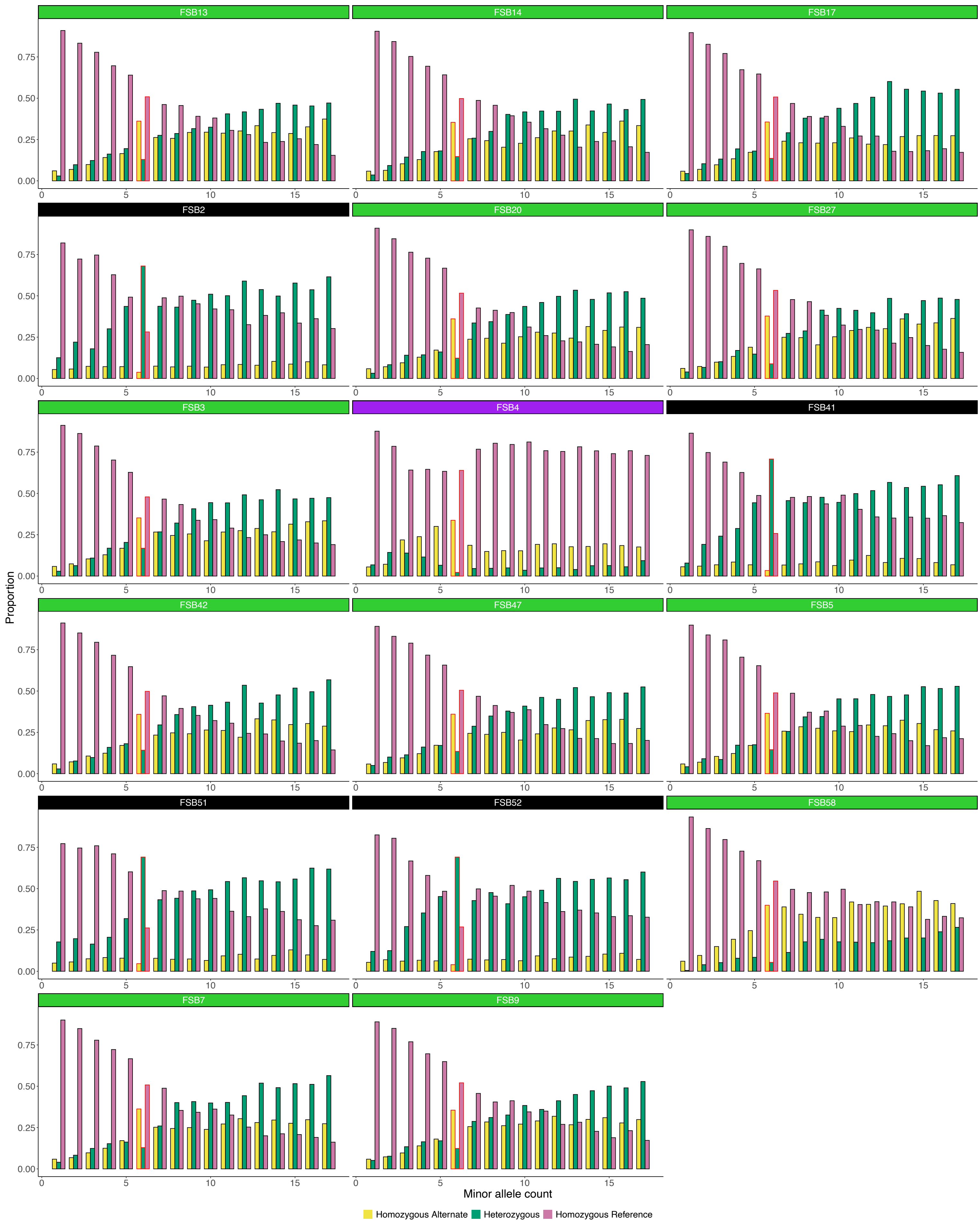
FSB13
FSB14
FSB17
FSB2
FSB20
FSB27
FSB3
FSB4
FSB41
FSB42
FSB47
FSB5
FSB51
FSB52
FSB58
FSB7
FSB9
Proportion
Minor allele count
Homozygous Alternate
Heterozygous
Homozygous Reference

**Figure S12. Inversion homozygotes in Maple Ridge show an enrichment of ALT alleles at minor allele frequencies approaching that of the inv21.0 polymorphism.** Each panel represents genotype proportions measured in a single Maple Ridge individual for SNPs spanning the inv21.0 interval. Panel headers denote the individual's sample ID and are colored to match the PCA cluster assignments given in Supplementary Figure S9 for inv21.0 SNPs (left cluster homozygotes: green, middle cluster heterozygotes: black, right cluster homozygotes: purple). Bars represent the individual's proportion of each genotype (pink for REF/REF, green for HET, and yellow for ALT/ALT) among SNPs at each minor allele frequency bin (denoted by x-axes). The group of bars outlined in red in each facet denote the minor allele frequency at which inv21.0 is segregating the Maple Ridge sample. For the right cluster individual (FSB4; in purple), there is an enrichment of REF alleles at minor allele frequencies exceeding that of the inv21.0 arrangement. In contrast, for left cluster individuals (FSB13, FSB14, FSB17, FSB20, FSB27, FSB3, FSB42, FSB47, FSB5, FSB58, FSB7, and FSB9; in green), there is an enrichment of ALT alleles at these higher minor allele frequencies.

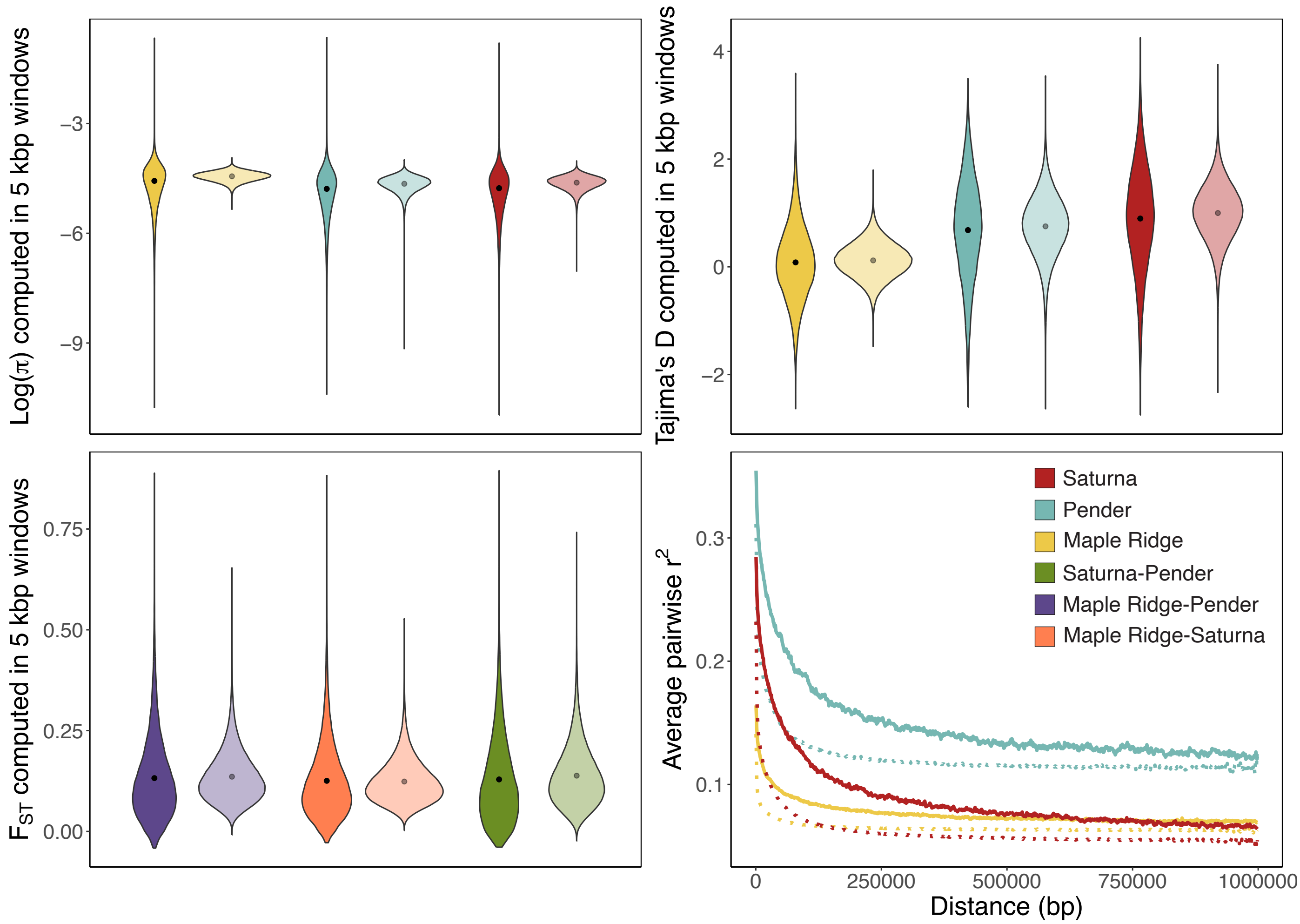


**Figure S13. Inferred demographic model provides a close fit to diversity-based summaries but underpredicts levels of LD.** Coalescent simulations conducted with the inferred demographic parameters were used to evaluate the fit of the model to empirical estimates of $\pi$, Tajima's D, $F_{ST}$, and LD. Violin plots compare simulated and empirical distributions of $\pi$ (top left), Tajima's D (top right), and $F_{ST}$ (bottom left) computed in non-overlapping 5 kbp windows. The color of each violin represents the population (yellow for Maple Ridge, blue for Pender Island, and red for Saturna Island) or population pair (purple for Maple Ridge-Pender, coral for Maple Ridge-Saturna, and green for Saturna-Pender) for which the given summary statistic was measured. Opaque distributions represent empirical results, and transparent distributions represent simulated results. Black points denote the mean value of each distribution. The bottom right panel compares simulated and observed LD decay. The y-axis plots average pairwise $r^2$ measured between SNPs separated by increasing distances (up to 1 Mbp; x-axis). Solid lines represent empirical results, and dashed lines represent simulated results. Curves are colored according to population.

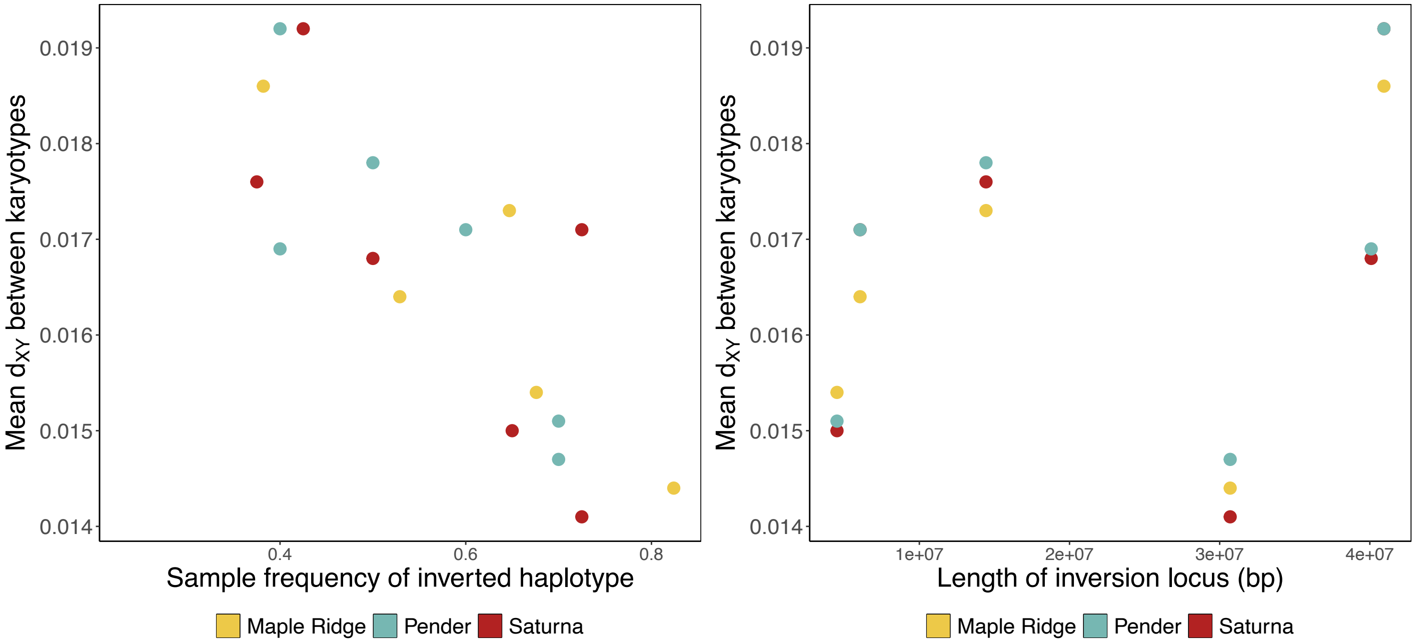


**Figure S14. Associations between inversion characteristics and the level of divergence between karyotypes.** Left panel depicts the relationship between mean $d_{XY}^{karyotype}$ (y-axis) and the observed frequency of the inversion in each population sample (x-axis). Right panel depicts the relationship between mean $d_{XY}^{karyotype}$ (y-axis) and the physical length of each inversion locus (x-axis). Each point represents a different polymorphic locus, colored according to population (yellow for Maple Ridge, blue for Pender Island, and red for Saturna Island). The inv14.0 locus is excluded for Maple Ridge in both panels due to a lack of inversion homozygotes. Although we find significant negative correlations between mean $d_{XY}^{karyotype}$ and inversion frequency in the Pender and Maple Ridge samples (Supplementary Table S2), we do not find a significant correlation between these variables in the Saturna sample, nor do we find significant correlations between $d_{XY}^{karyotype}$ and inversion length in any population sample. See Supplementary Table S2 for details.

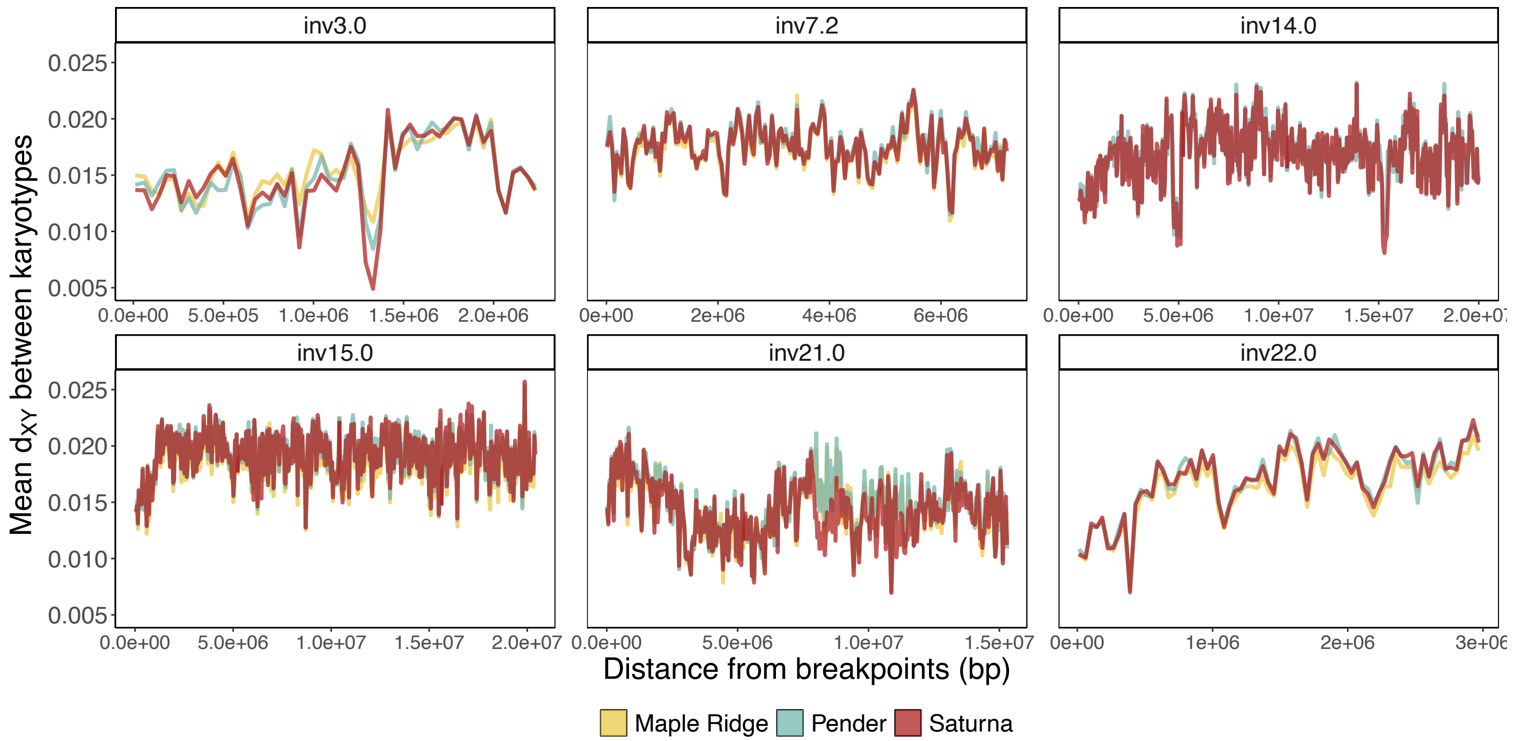


**Figure S15. Little evidence for genetic exchange between karyotypes at polymorphic inversions.** Panels plot divergence between karyotypes ($d_{XY}^{karyotype}$) computed in non-overlapping 5 kbp windows (y-axes) as a function of distance from inversion breakpoints (x-axes) for each of the six inversion loci segregating in the Gulf Islands populations. Lines are colored according to population (yellow for Maple Ridge, blue for Pender Island, and red for Saturna Island). Genetic exchange between karyotypes due to double crossovers and gene conversion events should occur most frequently near the center of inversion polymorphisms (i.e., furthest from the breakpoints), predicting a negative correlation between $d_{XY}^{karyotype}$ and distance from breakpoints. However, with the exception of the inv21.0 locus in Saturna, where there is a weakly negative correlation between $d_{XY}^{karyotype}$ and distance (r = -0.15; p-value = 3.24e-3), all other significant correlations are weakly positive. See Supplementary Table S3 for details.

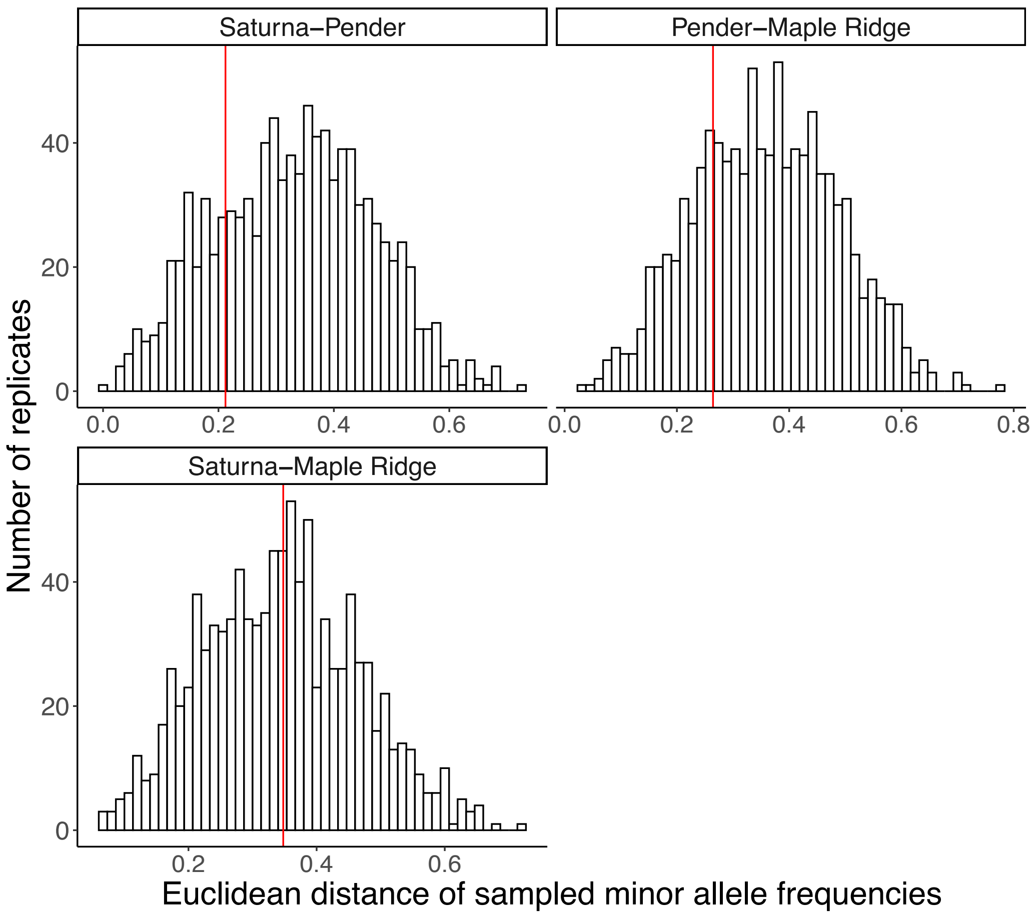


**Figure S16. Null sampling distributions of joint minor allele frequencies at high-confidence, putatively neutral, non-inverted SNP loci.** Histograms plot the distribution of Euclidean distances measured between each population (Saturna and Pender: top left, Pender and Maple Ridge: top right, Saturna and Maple Ridge: bottom left) at a random sample of six SNPs from the jSFS used for demographic inference. Euclidean distances for each sample of SNPs were computed based on each population's minor allele frequencies at the sampled SNPs. Vertical red lines denote where the same metric computed based on the minor allele frequencies of the six polymorphic inversions falls with respect to the null distribution.

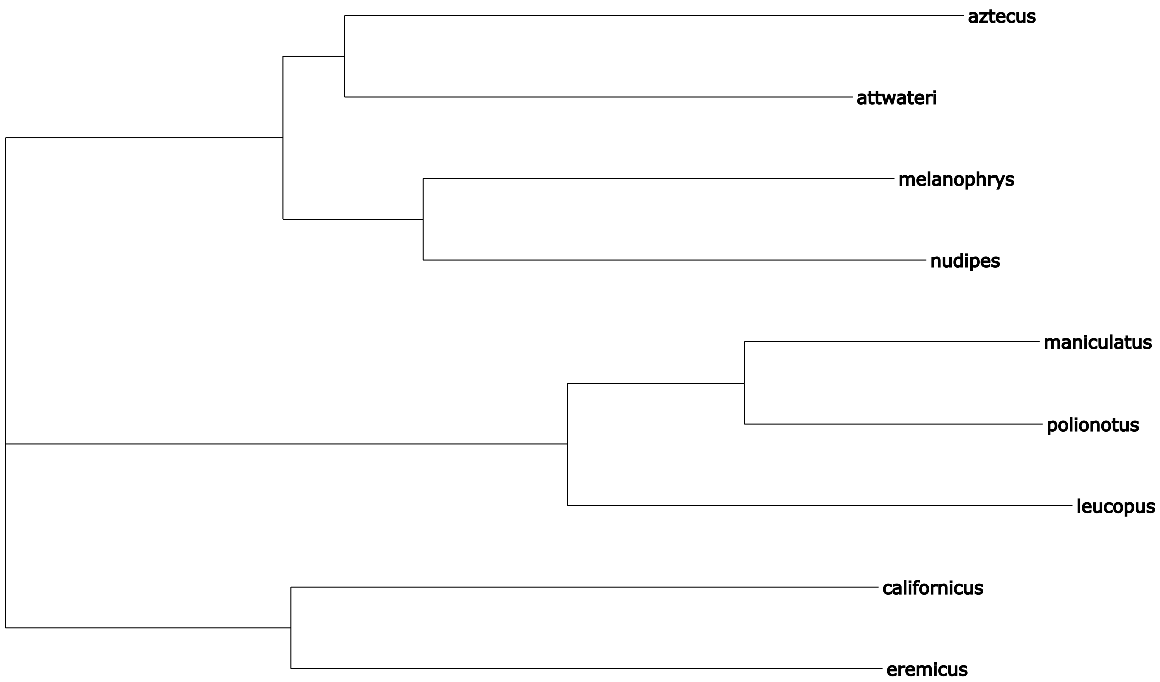


**Figure S17. Distance-based tree constructed from *Peromyscus* reference assembly recovers known phylogenetic relationships.** The Newick-formatted output from Mashtree (Katz et al. 2019) run on the FASTA files of nine NCBI RefSeq assemblies from diverse *Peromyscus* species was visualized with T-REX (Boc et al. 2012). Branch lengths are drawn proportional to the distance-based metric computed by Mashtree. Labels of leaves reflect species names. Four species: *P. polionotus*, *P. leucopus*, *P. californicus*, and *P. eremicus*, were used for downstream polarization of inversion polymorphisms in the *P. maniculatus* reference genome.

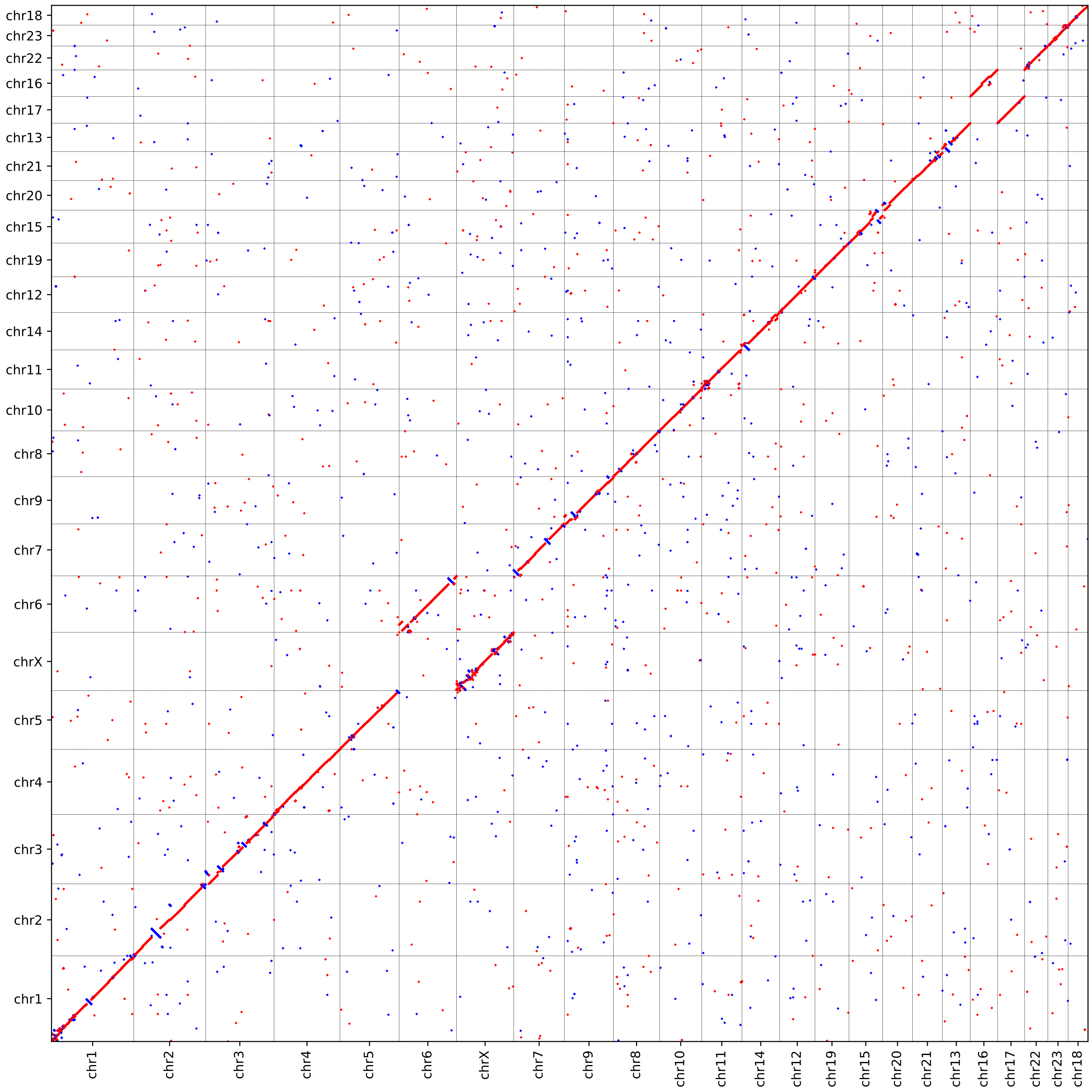


**Figure S18. Whole genome alignment between *P. maniculatus* and *P. polionotus* reference assemblies.** Dot plot represents aligned sequences between the two species. X-axis represents coordinates in the *P. maniculatus* genome. Y-axis represents coordinates in the *P. polionotus* genome. Chromosomes are named according to the assignments present in each assembly. Dots that lie on the anti-diagonal (in red) represent alignments with the same orientation in each assembly. Dots that lie on the main diagonal (in blue) represent alignments with the opposite orientation between assemblies

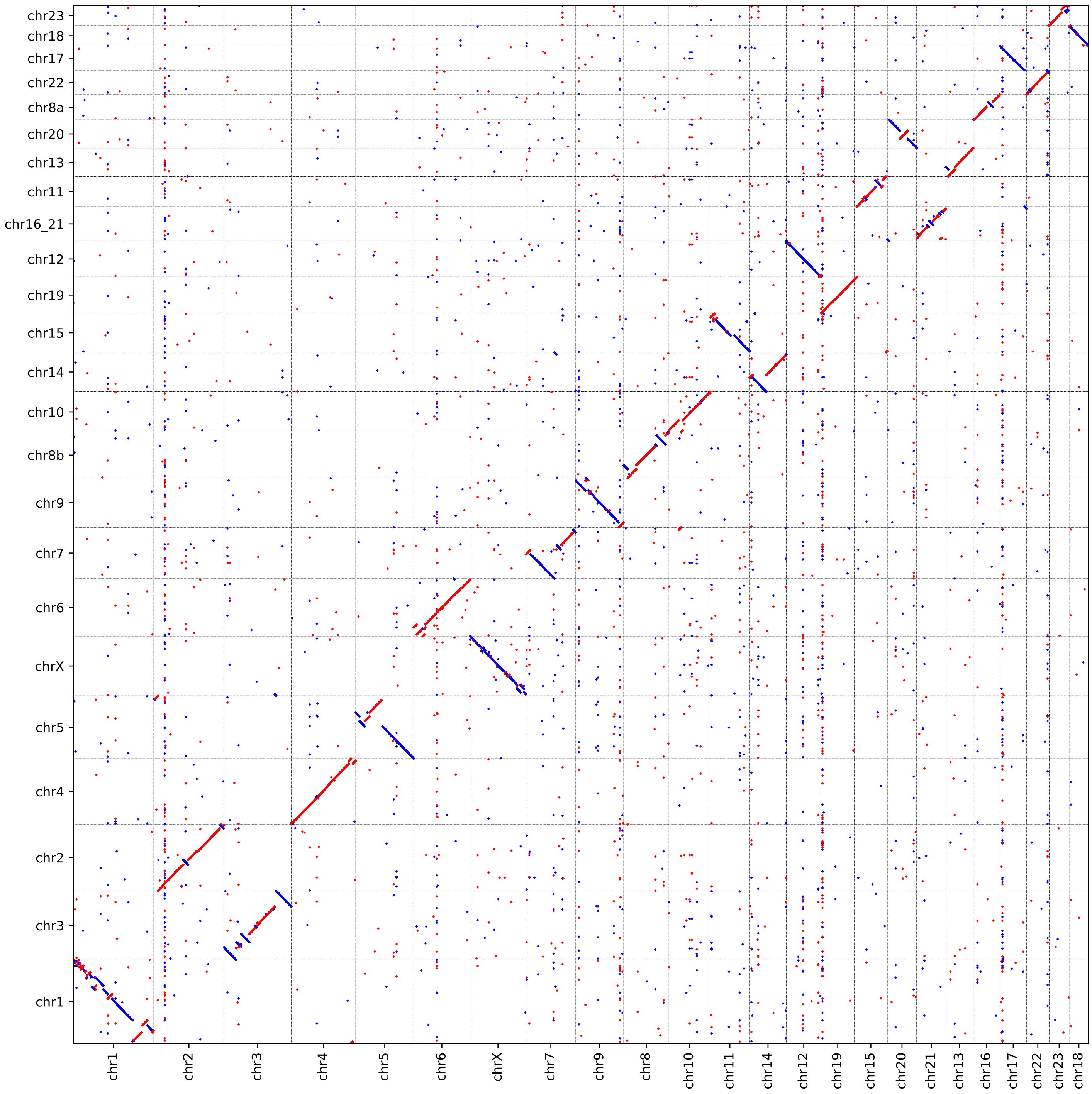


**Figure S19. Whole genome alignment between *P. maniculatus* and *P. leucopus* reference assemblies.** Dot plot represents aligned sequences between the two species. X-axis represents coordinates in the *P. maniculatus* genome. Y-axis represents coordinates in the *P. leucopus* genome. Chromosomes are named according to the assignments present in each assembly. Dots that lie on the anti-diagonal (in red) represent alignments with the same orientation in each assembly. Dots that lie on the main diagonal (in blue) represent alignments with the opposite orientation between assemblies.

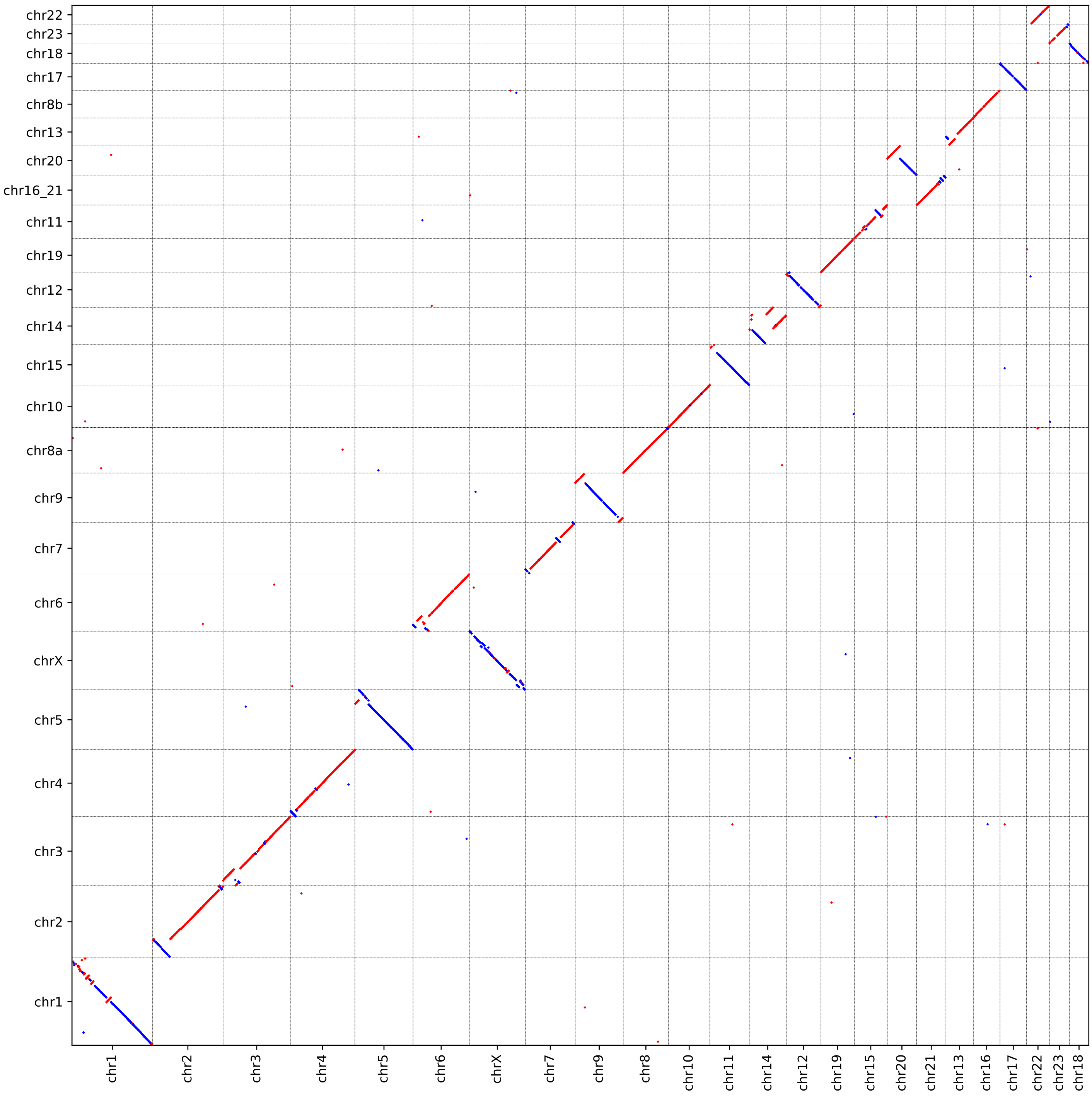


**Figure S20. Whole genome alignment between *P. maniculatus* and *P. eremicus* reference assemblies.** Dot plot represents aligned sequences between the two species. X-axis represents coordinates in the *P. maniculatus* genome. Y-axis represents coordinates in the *P. eremicus* genome. Chromosomes are named according to the assignments present in each assembly. Dots that lie on the anti-diagonal (in red) represent alignments with the same orientation in each assembly. Dots that lie on the main diagonal (in blue) represent alignments with the opposite orientation between assemblies.

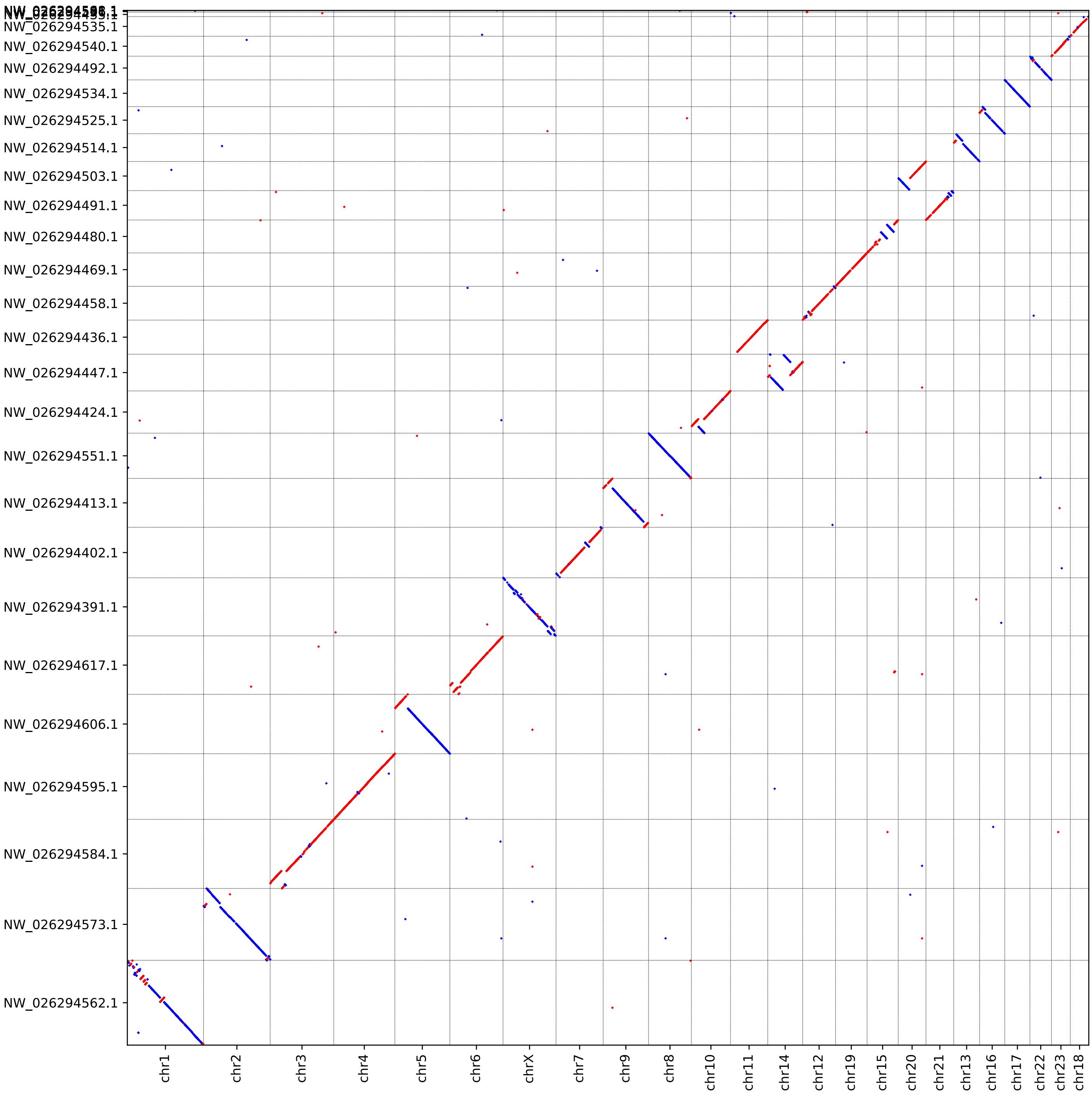


**Figure S21. Whole genome alignment between *P. maniculatus* and *P. californicus* reference assemblies.** Dot plot represents aligned sequences between the two species. X-axis represents coordinates in the *P. maniculatus* genome. Y-axis represents coordinates in the *P. californicus* genome. Chromosomes/contigs are named according to the assignments present in each assembly. Dots that lie on the anti-diagonal (in red) represent alignments with the same orientation in each assembly. Dots that lie on the main diagonal (in blue) represent alignments with the opposite orientation between assemblies.

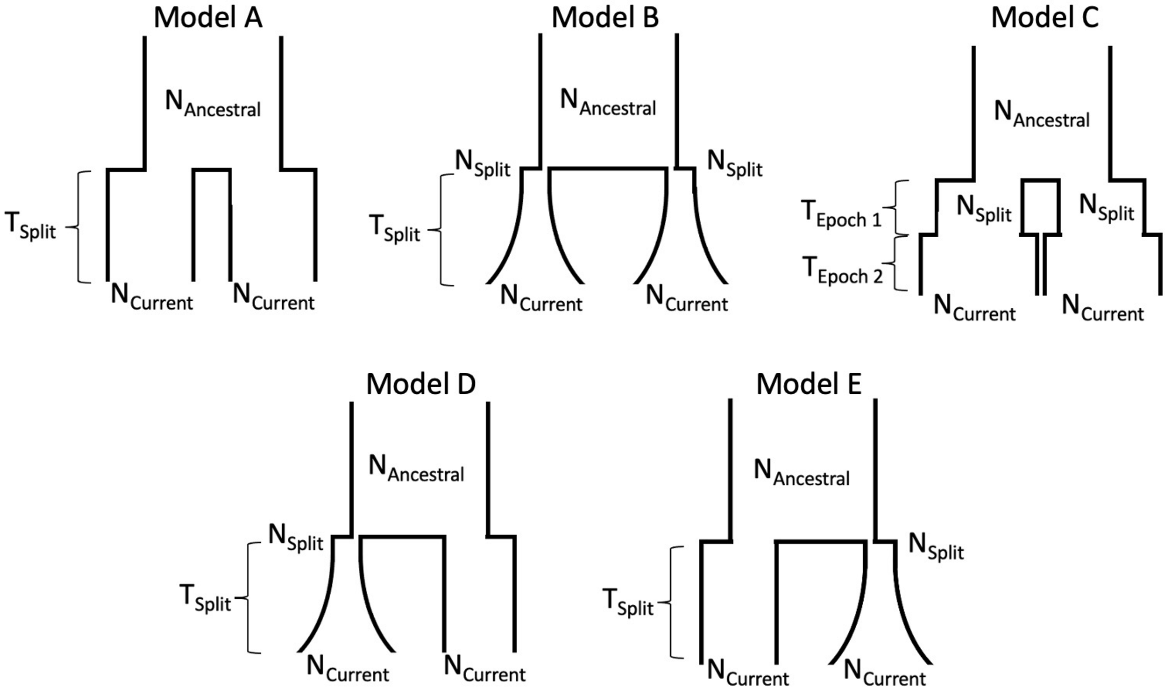


**Figure S22. Two-population demographic models.** Models A-E correspond to simplified models of population splits that involve either discrete (Models A and C), continuous (Model B), or a mixture of discrete and continuous (Model D and E) changes in effective population size following the split. Continuous changes in effective population size were modeled as exponential growth (or decline). Discrete changes were modeled as instantaneous growth (or decline). The direction of size change was not constrained by the model. All estimated parameters are labeled for each of the tested models.

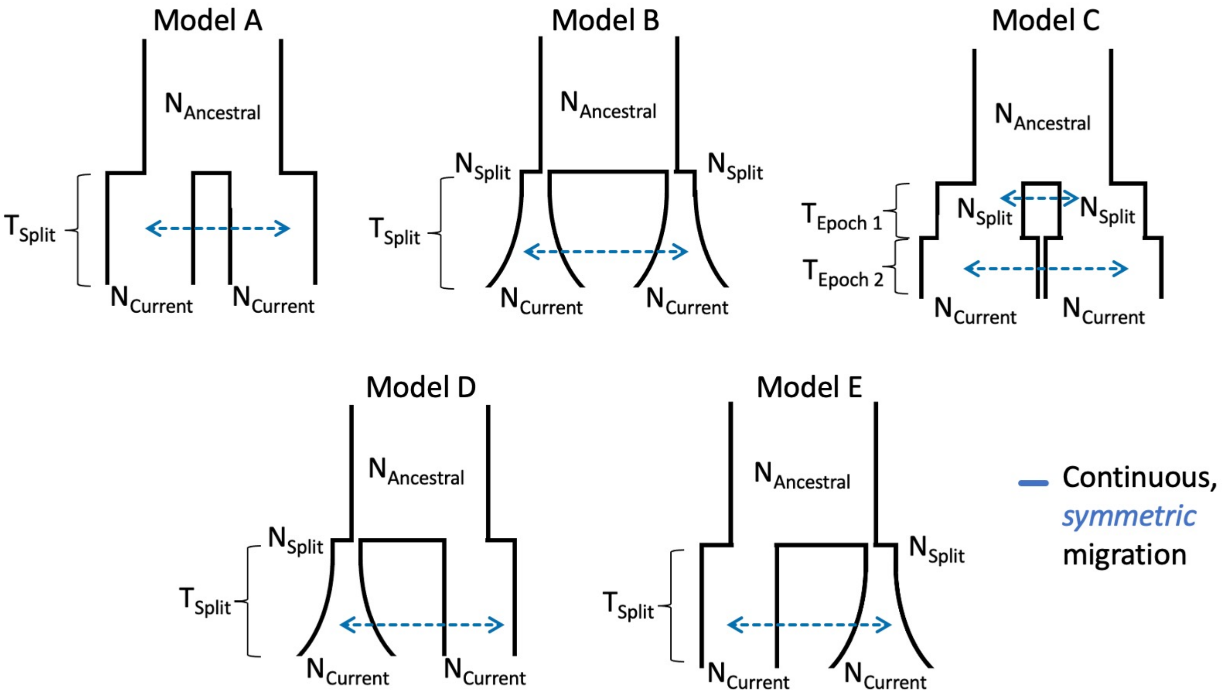


**Figure S23. Two-population demographic models with symmetric migration.** Models A-E are specified as in Supplementary Figure S22 with the addition of either one (Models A, B, D, and E) or two (Model C) migration rate parameters that model continuous, symmetric migration occurring between the two sampled populations after their divergence from a shared ancestor.

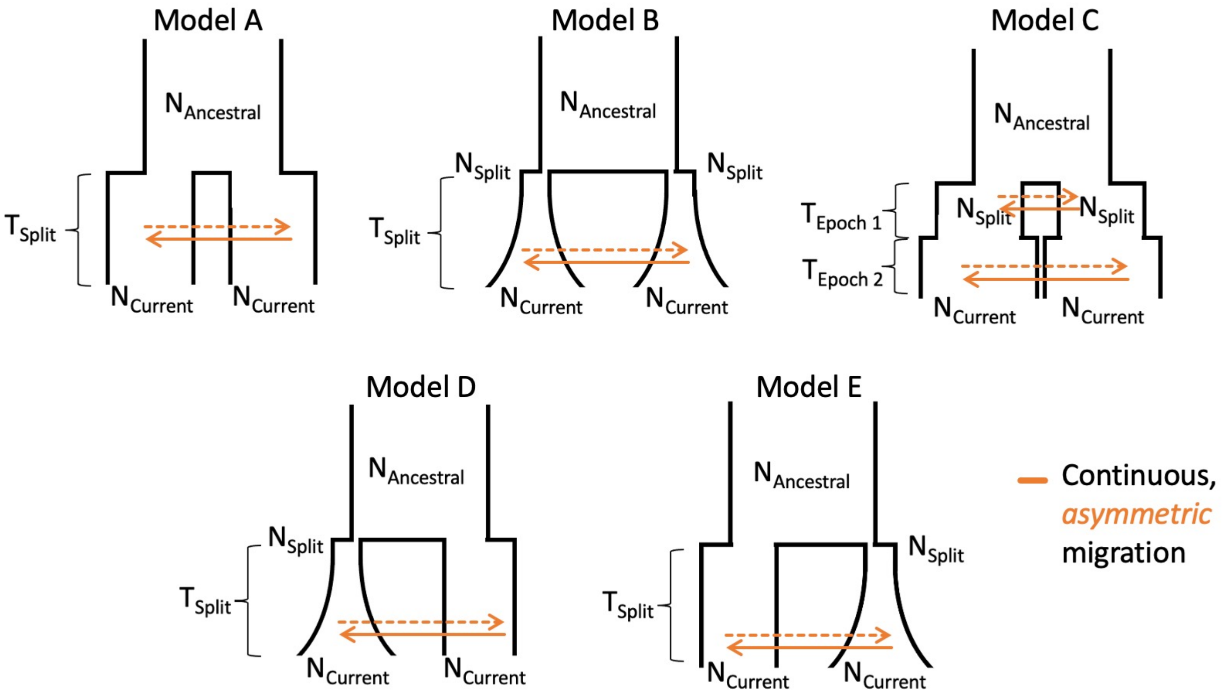


**Figure S24. Two-population demographic models with asymmetric migration.** Models A-E are specified as in Supplementary Figure S22 with the addition of either one (Models A, B, D, and E) or two (Model C) pairs of migration rate parameters that model continuous, asymmetric migration occurring between the two sampled populations after their divergence from a shared ancestor.

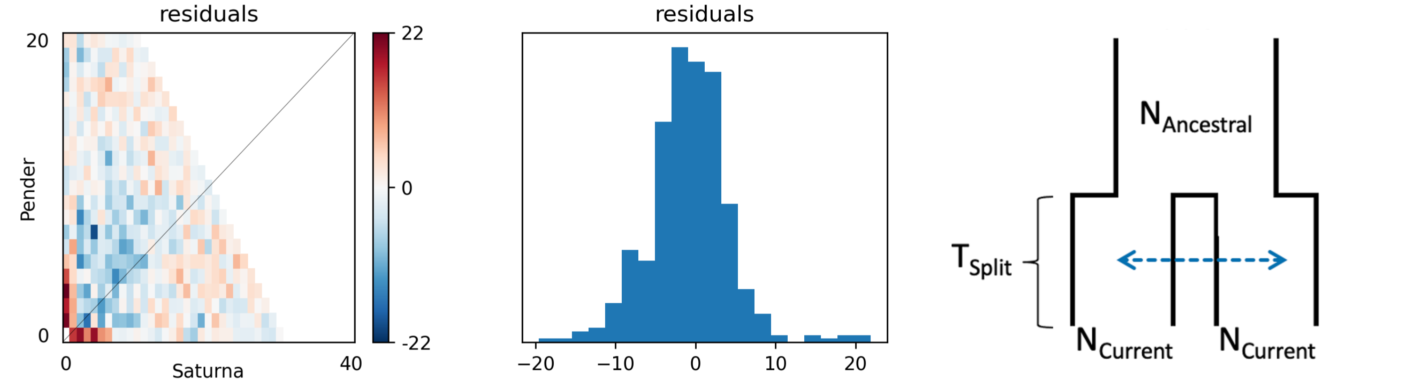


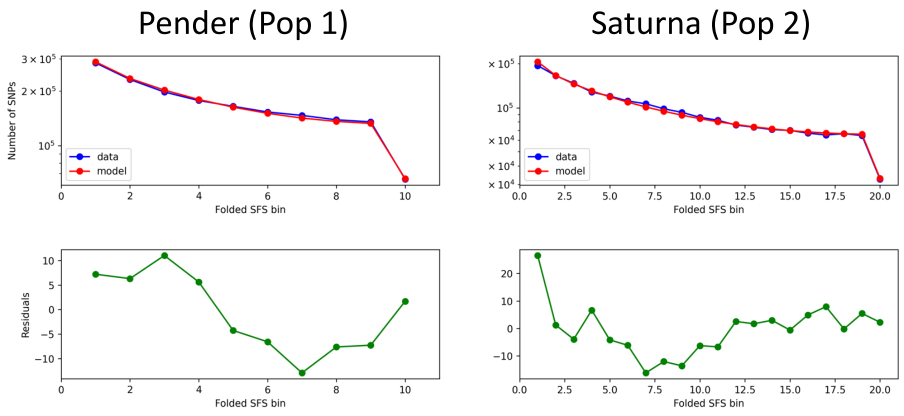


| Parameter | Estimate |
|---|---|
| Ancestral Size | 607,149 |
| Current Size (Pop 1) | 19,847 |
| Current Size (Pop 2) | 22,686 |
| Split Time | 7,360 |
| Migration | 8.23E-06 |
| Likelihood | -7,878 |

**Figure S25. Candidate two-population model fit for the Pender Island-Saturna Island comparison.** (Top right) Diagram of the best-fit model with estimated parameters labeled. (Bottom right) Table with maximum likelihood parameter estimates and model likelihood. (Top left) Residual differences between the expected joint site frequency spectrum (jSFS) and the observed jSFS. Each entry (*x*, *y*) in the jSFS represents the proportion of SNPs observed in *x* copies in the Saturna Island sample (x-axis) and *y* copies in the Pender Island sample (y-axis). The color of each cell represents the Anscombe residual difference, where cooler colors indicate that the model underpredicts the observed number of SNPs and warmer colors indicate an overprediction by the model. The accompanying histogram plots the distribution of residuals between the jSFS. (Bottom left) Residual differences (bottom panel, green) between the expected (top panel, red) and observed (top panel, blue) marginal SFS for the Pender Island (left) and Saturna Island (right) samples.

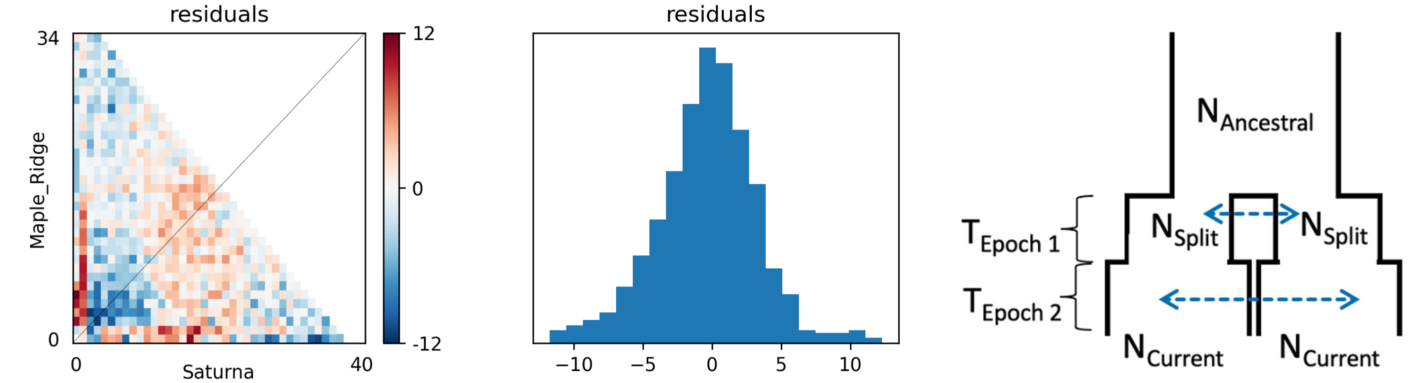


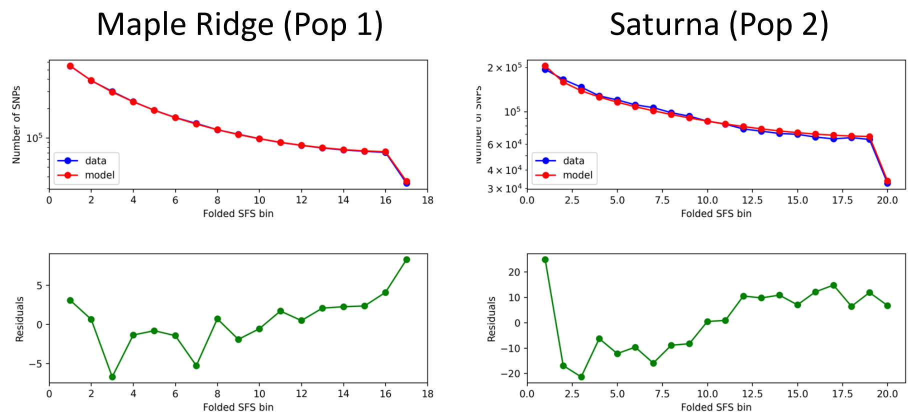


| Parameter | Estimate |
|---|---|
| Ancestral Size | 107,154 |
| Split Size (Pop 1) | 522,891 |
| Split Size (Pop 2) | 466,067 |
| Current Size (Pop 1) | 62,297 |
| Current Size (Pop 2) | 13,784 |
| Epoch 1 Duration | 1,976,998 |
| Epoch 2 Duration | 7,899 |
| Epoch 1 Migration | 4.13E-05 |
| Epoch 2 Migration | 1.26E-05 |
| Likelihood | -8,271 |

**Figure S26. Candidate two-population model fit for the mainland Maple Ridge-Saturna Island comparison.** (Top right) Diagram of the best-fit model with estimated parameters labeled. (Bottom right) Table with maximum likelihood parameter estimates and model likelihood. (Top left) Residual differences between the expected joint site frequency spectrum (jSFS) and the observed jSFS. Each entry (*x*, *y*) in the jSFS represents the proportion of SNPs observed in *x* copies in the Saturna Island sample (x-axis) and *y* copies in the mainland Maple Ridge sample (y-axis). The color of each cell represents the Anscombe residual difference, where cooler colors indicate that the model underpredicts the observed number of SNPs and warmer colors indicate an overprediction by the model. The accompanying histogram plots the distribution of residuals between the jSFS. (Bottom left) Residual differences (bottom panel, green) between the expected (top panel, red) and observed (top panel, blue) marginal SFS for the mainland Maple Ridge (left) and Saturna Island (right) samples.

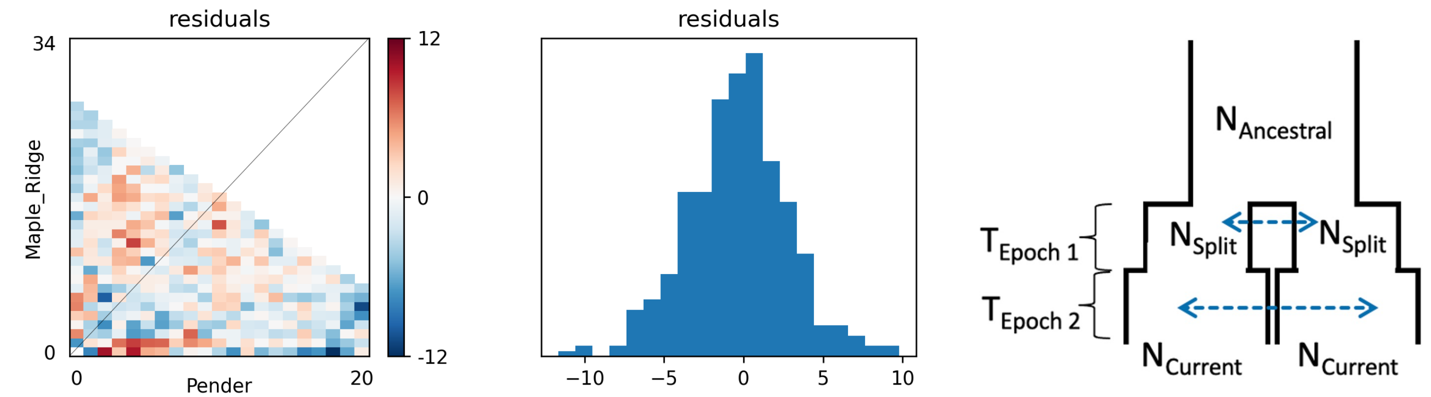


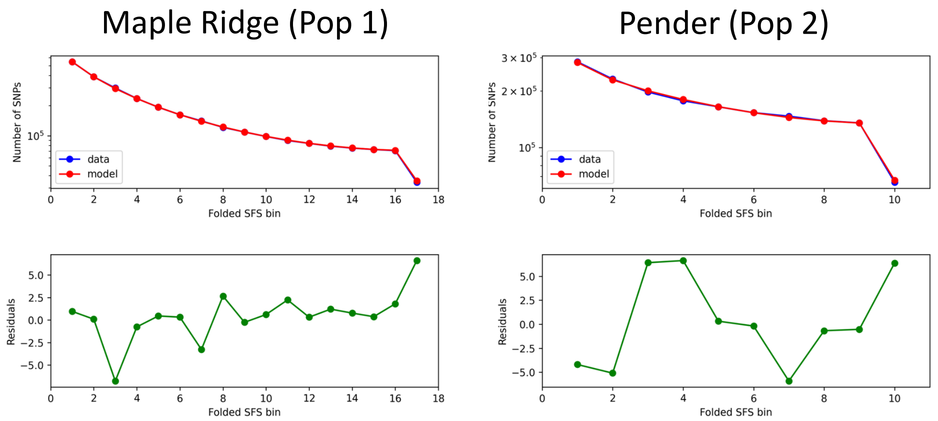


| Parameter | Estimate |
|---|---|
| Ancestral Size | 138,529 |
| Split Size (Pop 1) | 373,740 |
| Split Size (Pop 2) | 676,101 |
| Current Size (Pop 1) | 55,895 |
| Current Size (Pop 2) | 10,449 |
| Epoch 1 Duration | 1,796,672 |
| Epoch 2 Duration | 6,365 |
| Epoch 1 Migration | 3.58E-05 |
| Epoch 2 Migration | 1.42E-05 |
| Likelihood | -3,784 |

**Figure S27. Candidate two-population model fit for the mainland Maple Ridge-Pender Island comparison.** (Top right) Diagram of the best-fit model with estimated parameters labeled. (Bottom right) Table with maximum likelihood parameter estimates and model likelihood. (Top left) Residual differences between the expected joint site frequency spectrum (jSFS) and the observed jSFS. Each entry (*x*, *y*) in the jSFS represents the proportion of SNPs observed in *x* copies in the Pender Island sample (x-axis) and *y* copies in the mainland Maple Ridge sample (y-axis). The color of each cell represents the Anscombe residual difference, where cooler colors indicate that the model underpredicts the observed number of SNPs and warmer colors indicate an overprediction by the model. The accompanying histogram plots the distribution of residuals between the jSFS. (Bottom left) Residual differences (bottom panel, green) between the expected (top panel, red) and observed (top panel, blue) marginal SFS for the mainland Maple Ridge (left) and Pender Island (right) samples.

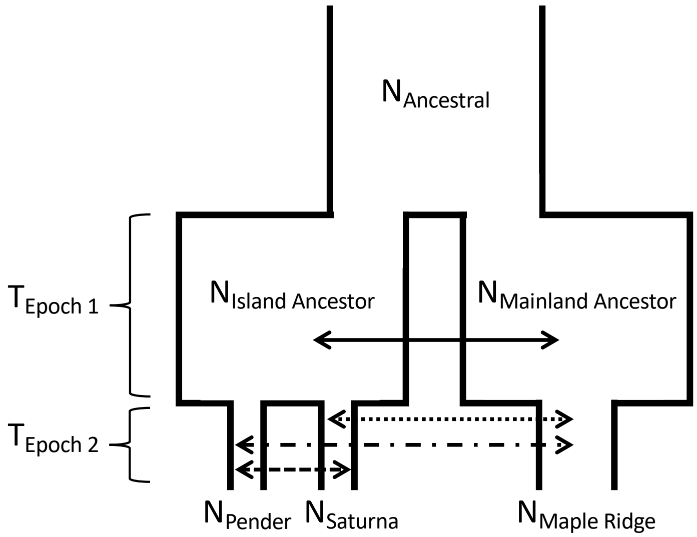


**Figure S28. Three-population demographic models.** Basic model involves an early epoch (Epoch 1) in which ancestral island and ancestral mainland populations diverge, followed by a recent epoch (Epoch 2) that involves the split of contemporary Saturna and Pender populations from the ancestral island population. All estimated population size and time interval parameters are labeled. Arrows denote symmetric migration between either the ancestral island and ancestral mainland population (solid), Saturna and Maple Ridge (dotted), Pender and Maple Ridge (dot dash), or Saturna and Pender (dashed). Tested models differ in the combination of symmetric migration parameters. We fit all combinations of migration parameters to the 3D jSFS. As with two-population models, the direction of size change was not constrained by the model.

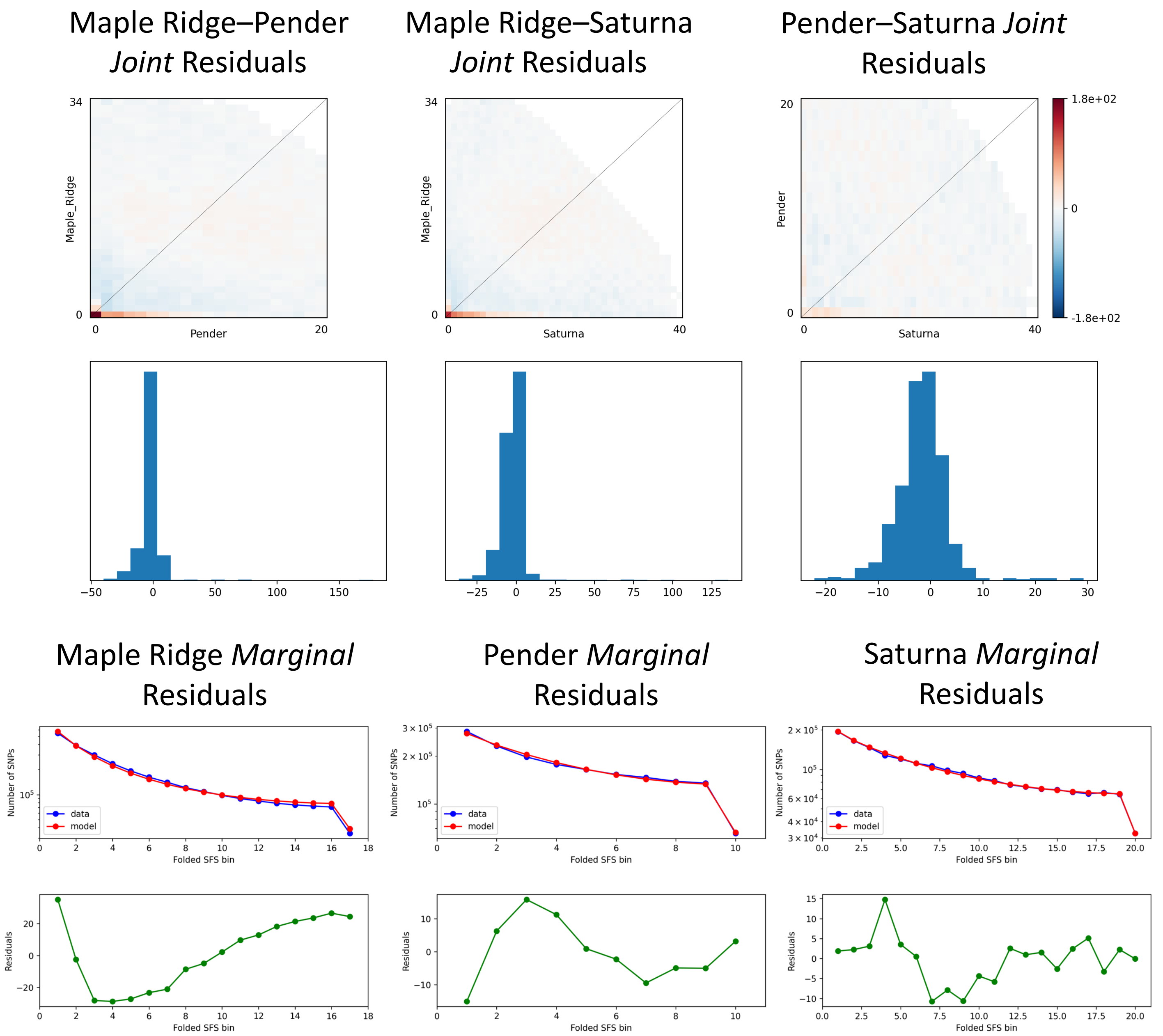


| Parameter | Estimate |
|---|---|
| Ancestral Size | 129,563 |
| Mainland-Ancestor Size | 579,644 |
| Island-Ancestor Size | 255,958 |
| Current Maple Ridge Size | 69,313 |
| Current Pender Size | 14,984 |
| Current Saturna Size | 18,607 |
| Epoch 1 Duration | 2,560,502 |
| Epoch 2 Duration | 5,090 |
| Mainland-Anc and Island-Anc Migration | 6.75E-06 |
| Pender-Saturna Migration | 1.63E-07 |
| Likelihood | -127,112 |

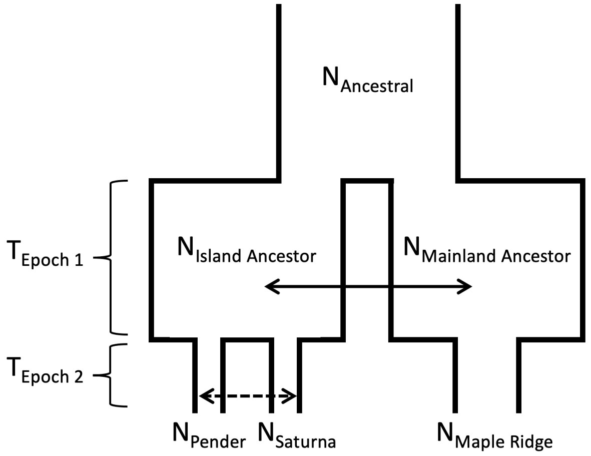

**Figure S29. Candidate three-population model fit.** (Top row) Residual differences between the expected 2D joint site frequency spectrum (jSFS) and the observed jSFS for each population pair (Maple Ridge-Pender: left, Maple Ridge-Saturna: middle, and Pender-Saturna: right). Each entry (*x*, *y*) in each jSFS represents the proportion of SNPs observed in *x* copies in the population sample represented on the x-axis and *y* copies in the population sample represented on the y-axis. The color of each cell represents the Anscombe residual difference, where cooler colors indicate that the model underpredicts the observed number of SNPs and warmer colors indicate an overprediction by the model. The accompanying histograms plot the distribution of residuals. (Middle row) Residual differences (bottom panels, green) between the expected (top panels, red) and observed (top panels, blue) marginal SFS for each population (Maple Ridge: left, Pender: middle, and Saturna: right). (Bottom row) Table with maximum likelihood parameter estimates and model likelihood (left) of the best-fit model (right).
